# Human Abdominal Subcutaneous-Derived Active Beige Adipocytes Carrying *FTO* rs1421085 Obesity-Risk Alleles Exert Lower Thermogenic Capacity

**DOI:** 10.1101/2023.01.30.525688

**Authors:** Attila Vámos, Rini Arianti, Boglárka Ágnes Vinnai, Rahaf Alrifai, Abhirup Shaw, Szilárd Póliska, Andrea Guba, Éva Csősz, István Csomós, Gábor Mocsár, Cecilia Lányi, Zoltán Balajthy, László Fésüs, Endre Kristóf

## Abstract

White adipocytes store lipids, have a large lipid droplet and few mitochondria. Brown and beige adipocytes, which produce heat, are characterized by high expression of uncoupling protein (UCP) 1, multilocular lipid droplets, and large amounts of mitochondria. The rs1421085 T-to-C single-nucleotide polymorphism (SNP) of the human *FTO* gene interrupts a conserved motif for ARID5B repressor, resulting in adipocyte type shift from beige to white. We obtained abdominal subcutaneous adipose tissue from donors carrying *FTO* rs1421085 TT (risk-free) or CC (obesity-risk) genotypes, isolated and differentiated their preadipocytes into beige adipocytes (driven by the PPARγ agonist rosiglitazone for 14 days), and activated them with dibutyryl-cAMP for 4 hours. Then, either the same culture conditions were applied for additional 14 days (active beige adipocytes) or it was replaced by a white differentiation medium (inactive beige adipocytes). White adipocytes were differentiated by their medium for 28 days. RNA-sequencing was performed to investigate the gene expression pattern of adipocytes carrying different *FTO* alleles and found that active beige adipocytes had higher brown adipocyte content and browning capacity compared to white or inactive beige ones when the cells were obtained from risk-free TT but not from obesity-risk CC genotype carriers. Active beige adipocytes carrying *FTO* CC had lower thermogenic gene (e.g., *UCP1, PM20D1, CIDEA*) expression and thermogenesis measured by proton leak respiration as compared to TT carriers. In addition, active beige adipocytes with CC alleles exerted lower expression of ASC1 neutral amino acid transporter (encoded by *SLC7A10*) and less consumption of Ala, Ser, Cys, and Gly as compared to risk-free carriers. We did not observe any influence of the *FTO* rs1421085 SNP on white and inactive beige adipocytes highlighting its exclusive and critical effect when adipocytes were activated for thermogenesis.

## 1 Introduction

In the last few decades, the prevalence of obesity dramatically increased across the world. Chronic obesity can lead to various cancers, type 2 diabetes, and cardiovascular disease. Therefore, obesity has been identified as one of the globally leading causes of mortality and disability, and is responsible for 10-13% of deaths in different regions worldwide [Frühbeck *et al*., 2013]. Imbalance of energy homeostasis, when energy intake is significantly greater than energy expenditure, has been identified as the main pathophysiological cause of obesity [Heymsfield and Wadden, 2017]. However, obesity is a multifactorial disease which can be the result of factors including social, lifestyle, behavioral networks, and genetic background of the individuals [Christakis and Fowler, 2007; Frühbeck *et al*., 2018; Lin and Li, 2021].

In rodents, adipocytes are classified into three types. The energy storing white adipocytes have one large unilocular lipid droplet and low mitochondrial density. The brown adipocytes located in the brown adipose tissue (BAT) are active thermogenic cells with high mitochondrial abundance, fragmentation, and uncoupling protein (UCP) 1 expression, as well as numerous small lipid droplet content in the cytoplasm. The “brown-like-in-white” (brite) or beige cells have cold-inducible thermogenic potential and multilocular lipid droplets [Sanchez-Gurmaches *et al*., 2016]. Beige adipocytes are interspersed in the white adipose tissue (WAT). In basal state, their gene expression pattern is similar to the white adipocytes, but upon extended stimuli (cold exposure, β-adrenergic stimulation, peroxisome proliferator-activated receptor (PPAR) γ activation) they exhibit a brown-like phenotype acquired in a process called browning [Petrovic *et al*., 2010; Wu *et al*., 2012]. Inguinal WAT has been discovered as a natural beige adipocyte depot, in which the adipocytes possess multilocular morphology and thermogenic gene expression profile in response to thermogenic cues [Chan *et al*., 2019].

In humans, BAT was primarily regarded as a tissue which is only present in infants and located at anatomical sites that are difficult to reach [Heaton, 1972]. Several studies using positron emission tomography (PET) provided evidence that adults have significant amounts of BAT, most commonly the cervical-supraclavicular depot was marked by high labeled glucose uptake especially after cold exposure [Cypess *et al*., 2009; Virtanen *et al*., 2009]. Using an elegant approach of the PET-Computed Tomography (CT) method, brownable adipose tissue could be found interspersed in several areas, such as cervical, supraclavicular, axillary, mediastinal, paraspinal, and abdominal [Leitner *et al*., 2017]. However, unlike in rodents, the molecular characteristics of human BAT remain elusive. Several studies reported that human BAT isolated from cervical-supraclavicular depots [Cypess *et al*., 2013] and primary adipocytes derived from fetal interscapular adipose tissue [Seiler *et al*., 2015] possess classical brown adipocyte characteristics marked by high expression of zinc finger protein-1 (ZIC1), whereas other studies using clonally derived human brown adipocytes isolated from supraclavicular depot reported the existence of a population of UCP1-positive cells displaying beige adipocyte features [Shinoda *et al*., 2015]. Another study using total RNA isolated from fat biopsies from various anatomical locations, including subcutaneous (SC) supraclavicular, posterior mediastinal, retroperitoneal, intra-abdominal, or mesenteric depots reported that beige-selective markers, such as *HOXC8, HOXC9*, and *CITED1* were highly expressed in human thermogenic adipose tissue, whereas classical BAT markers were not detectable [Sharp *et al*., 2012].

The activation of UCP1 to generate heat by brown/beige adipocytes drives a higher uptake of fuels, such as glucose and fatty acids, to sustain the metabolic substrates for tricarboxylic acid (TCA) cycle and generate NADH and FADH_2_ that subsequently enter the electron transport chain. Active brown/beige adipocytes also take up large amounts of TCA cycle intermediates, e.g. succinate, to enhance their proton leak respiration [Mills *et al*., 2018]. A recent study reported that the labeled glucose consumed by murine BAT during cold exposure is converted to pyruvate, which is further oxidized to acetyl-CoA catalyzed by pyruvate dehydrogenase [Panic *et al*., 2019]. In addition to glucose and fatty acids, active brown/beige adipocytes also catabolize branched-chain amino acids to fulfill the high demand of energy [Yoneshiro *et al*., 2019]. Our previous study also underlined the importance of alanine-serine-cysteine transporter 1 (ASC1)-mediated consumption of serine, cysteine, and glycine for efficient thermogenic response upon adrenergic stimulation in deep neck derived adipocytes [Arianti *et al*., 2021]. The capability of active brown/beige adipocytes as metabolic sink may contribute to the clearance of blood glucose and lipids, which indirectly improves glucose tolerance and insulin sensitivity [Cheng *et al*., 2020]. High rate of metabolic substrate utilization by active brown/beige adipocytes enhances the energy expenditure, therefore it may promote weight loss and become a potential pharmaceutical target to treat obesity and related metabolic diseases.

Recent publications reported the involvement of autophagy in the downregulation of beige adipocyte thermogenesis. In rodents, parkin-dependent selective mitochondrial clearance (mitophagy) drives the generation of inactive – morphologically white, but reactivation capable – masked beige adipocytes [Altshuler-Keylin *et al*., 2016]. In human abdominal SC adipocytes, both parkin-dependent and parkin-independent mitophagy related genes were upregulated upon *ex vivo* beige to white transition [Vámos *et al*., 2022]. In contrast, in differentiated human primary SC and Simpson– Golabi–Behmel syndrome (SGBS) adipocytes, cAMP-induced thermogenic activation downregulated mitophagy blocking beige to white transition [Szatmári-Tóth *et al*., 2020]. Preventing entry into this conversion might be a potential way to maintain elevated thermogenesis for combatting obesity.

Individual susceptibility to obesity is determined by interactions between genetic background and behavior. Genome-wide association studies identified the strong association between obesity and the Fat mass and obesity-associated (*FTO*) gene [Wang *et al*., 2011; Dina *et al*., 2007]. Among several identified single-nucleotide polymorphisms (SNPs) of *FTO*, an intronic rs1421085 T-to-C SNP has recently attracted attention. Studies in European and Japanese populations reported that the presence of the obesity-risk C allele increased the susceptibility for obesity, elevated fat mass and food intake [Wheeler *et al*., 2013; Felix *et al*., 2016; Imamura *et al*., 2016; Tanaka *et al*., 2013]. Claussnitzer *et al*. (2015) elucidated the molecular background for the association between *FTO* rs1421085 SNP and increased fat storage. In the presence of the risk-free allele (T), the AT-Rich Interaction Domain 5B (ARID5B) repressor protein can bind to the enhancer region of Iroquois Homeobox (IRX) 3 and 5, therefore the expression of IRX3 and 5 is suppressed [Claussnitzer *et al*., 2015]. When the obesity-risk allele is present, the conserved motif for ARID5B repressor is disrupted resulting in elevated expression of IRX3 and 5. As the consequence, the differentiation program is shifted from energy dissipation by beige adipocytes to lipid storage into white adipocytes [Claussnitzer *et al*., 2015; Herman and Rosen, 2015]. IRX5^-/-^ mice possessed reduced fat mass and did not develop obesity when fed on a high-fat diet. In addition, *IRX5* silencing increased the mitochondrial respiration in isolated mouse adipocytes [Bjune *et* al., 2005]. IRX3^-/-^ ME3 murine embryonic fibroblast line failed to differentiate to beige adipocytes, however, increased their capacity for chondrogenesis [Bjune *et al*., 2020]. On the other hand, it was also reported that IRX3 promotes the browning of white adipocytes as it can directly bind to the promoter of *UCP1* [Zou *et* al., 2017].

In this study, we employed transcriptomic and metabolomic data to investigate the effect of *FTO* rs1421085 SNP on the thermogenic capacity of three types of adipocytes: white, active and inactive beige, which were derived from human adipose-derived stromal cells (hASCs) isolated from abdominal SC fat of donors carrying *FTO* risk-free (TT) or obesity-risk (CC) genotypes (4 individuals of each genotype). RNA-sequencing (RNA-seq) analysis was performed to screen the global transcriptomic profiles of the differentiated adipocytes and we found that active beige adipocytes carrying risk-free alleles had higher brown adipocyte content, browning capacity, mitochondrial complex I, II, and IV subunit amount, proton leak respiration, extracellular acidification, expression of thermogenic markers (*UCP1, PM20D1, CITED1, CIDEA, CKMT1*, and *CKMT2*), and ASC1-mediated amino acid consumption, as compared to white or inactive beige adipocytes carrying the same genotypes. Intriguingly, we found that active beige adipocytes carrying *FTO* obesity-risk genotypes have less distinguishable characteristics as compared to white or inactive beige adipocytes. Our findings underline the critical effect of *FTO* rs1421085 SNP in human abdominal SC adipocytes when they are activated for thermogenesis.

## 2 Materials and methods

### 2.1 Materials

All chemicals were obtained from Sigma-Aldrich (Munich, Germany) unless stated otherwise.

### 2.2 Ethical Statement and Obtained hASCs

The human SC abdominal adipose tissue collection was approved by the Medical Research Council of Hungary (20571-2/2017/EKU) followed by the EU Member States’ Directive 2004/23/EC on presumed consent practice for tissue collection. All experiments were implemented in accordance with the Helsinki Declaration. All participants were informed about the experiments and written informed consent was obtained from them. hASCs were obtained and isolated from stromal vascular fractions of human SC abdominal adipose tissue of healthy donors undergoing plastic surgery, as previously described [Kristóf *et al*., 2019; Szatmári-Tóth *et al*., 2020].

### 2.3 Maintenance and Differentiation of hASCs

hASCs were seeded in 6-well plates and cultured in Dulbecco’s Modified Eagle’s Medium/Nutrient F-12 Ham (DMEM-F12) medium containing 17 µM pantothenic acid, 33 µM biotin, 100 U/ml penicillin/streptomycin and 10% fetal bovine serum (Thermo Fisher Scientific, Waltham, MA, USA) at 37°C in 5% CO_2_ until they reach complete confluence. The absence of mycoplasma was verified by polymerase chain reaction (PCR) analysis (PCR Mycoplasma Test Kit I/C, PromoKine, PromoCell, Heidelberg, Germany).

White adipogenic differentiation was induced for three days with serum-free DMEM-F12 medium supplemented with 17 µM pantothenic acid, 33 µM biotin, 100 U/ml penicillin/streptomycin, 100 nM cortisol, 10 µg/ml human apo-transferrin, 200 pM triiodothyronine, 20 nM human insulin, 2 µM rosiglitazone (Cayman Chemicals, Ann Arbor, MI, USA), 25 nM dexamethasone and 500 µM 3-isobutyl-l-methylxantine. After the third day, rosiglitazone, dexamethasone and 3-isobutyl-l-methylxantine were removed from the medium during the remaining 25 days of the differentiation. The medium was exchanged in every third day.

The active beige differentiation was induced for three days with serum-free DMEM-F12 medium supplemented with 17 µM pantothenic acid, 33 µM biotin, 100 U/ml penicillin/streptomycin, 10 µg/ml human apo-transferrin, 200 pM triiodothyronine, 20 nM human insulin, 2 µM rosiglitazone, 25 nM dexamethasone and 500 µM 3-isobutyl-l-methylxantine. After the third day, dexamethasone and 3-isobutyl-l-methylxantine were removed and 500 nM rosigliazone was added to the medium for the remaining 25 days of differentiation. On the fourteenth day, a 4 hours long 500 µM dibutyryl-cAMP treatment was carried out to mimic the *in vivo* cold-induced thermogenesis [Szatmári-Tóth *et al*., 2020]. After the treatment, the aforementioned beige differentiation medium was applied until the end of the differentiation.

The inactive beige differentiation was induced in the first fourteen days as in the case of the active beige adipocytes, but after the dibutyryl-cAMP treatment, the medium was replaced to the white cocktail without rosiglitazone, dexamethasone and 3-isobutyl-l-methylxantine for additional fourteen days.

### 2.4 RNA Isolation, Reverse-Transcription PCR (RT-PCR), Quantitative PCR (qPCR), and RNA-Seq analysis

Adipocytes were collected in TRIzol reagent (Thermo Fisher Scientific), followed by manual RNA isolation by chloroform extraction and isopropanol precipitation. The RNA quality was evaluated by Nanodrop (Thermo Fisher Scientific). cDNA generation was carried out by TaqMan reverse transcription reagent kit (Thermo Fisher Scientific) and followed by qPCR analysis [Szatmári-Tóth *et al*., 2020; Shaw *et al*., 2021]. Gene expressions were normalized to *GAPDH*. All TaqMan assays are listed in Supplementary Table 1.

Total RNA sample quality was checked on Agilent BioAnalyzer using Eukaryotic Total RNA Nano Kit according to the Manufacturer’s protocol. Samples with RNA integrity number (RIN) value > 7 were accepted for the library preparation process. RNA-Seq libraries were prepared from total RNA using MGIEasy RNA Library Prep Set V3.0 (MGI, Shenzhen, China) according to the manufacturer’s protocol. Briefly, poly-A RNAs were captured by oligo-dT conjugated magnetic beads then the mRNAs were eluted and fragmented at 94 °C. First-strand cDNA was generated by random priming reverse transcription, then in the second strand synthesis step, double-stranded cDNA was generated. After repairing ends, A-tailing and adapter ligation steps, adapter-ligated fragments were amplified in enrichment PCR and finally sequencing libraries were generated. In the next step double-stranded libraries denatured and single strand circularization performed, after enzymatic digestion step circularized single-stranded library was generated for DNA nano ball (DNB) generation. After making DNB step, single-end 100 cycles sequencing run was executed on MGI DNBSEQ G400 instrument. After sequencing, the reads were aligned to the GRCh38 reference genome (with EnsEMBL 95 annotation) using STAR aligner (version 2.7.0a). FeatureCounts was used to quantify our reads to genes. Significantly differentially expressed genes (DEGs) were defined based on adjusted p values < 0.05 and log2 fold change threshold > 0.85. Heatmap was generating by using GraphPad 8.0 Software and interactome map was constructed by using Gephi 0.9 based on interaction from STRING (https://string-db.org/). Pathway analysis was performed by subjecting the list of DEGs to STRING and overrepresented KEGG pathways were selected based on the false discovery rate (FDR) <0.05. Brown adipocyte content and browning capacity was estimated by BATLAS and ProFAT webtools, respectively, by subjecting the transcriptomic data of both groups of markers [Perdikari *et al*., 2018; Cheng *et al*., 2018].

### 2.5 *FTO* allele Genotyping

DNA isolation was performed as previously described [Klusóczki *et al*., 2019]. Rs1421085 SNP was genotyped by qPCR using TaqMan SNP Genotyping assay (Thermo Fisher Scientific, 4351379) according to the Manufacturer’s instructions.

### 2.6 Antibodies and immunoblotting

The separation of investigated proteins was performed by SDS-PAGE, followed by transfer to a PVDF membrane, which was blocked by 5% skimmed milk solution [Szatmári-Tóth *et al*., 2020; Shaw *et al*., 2021]. The primary antibodies were utilized in the following dilutions: anti-β-actin (1:5000, A2066), anti-UCP1 (1:750, R&D Systems, Minneapolis, MN, USA, MAB6158), anti-OXPHOS (1:1000, Abcam, Cambridge, MA, USA, ab110411), anti-SLC7A10 (1:500, Abnova, Taipei City, Taiwan, H00056301-B01P), anti-PGC1α (1:1000, Novus Biologicals, Centennial, CO, USA, NBP1-04676) anti-GPT2 (1:2000, Thermo Fisher Scientific, PA5-62426), and anti-SHMT1 (1:2000, Thermo Fisher Scientific, PA5-88581). The following species corresponding secondary antibodies were used: HRP-conjugated goat anti-mouse (1:5000, Advansta, San Jose, CA, USA, R-05071-500) or anti-rabbit (1:5000, Advansta, R-05072-500) IgG. The expression of the visualized immunoreactive proteins were quantified by densitometry using the FIJI ImageJ software (National Institutes of Health (NIH), Bethesda, MD, USA) as previously described [Szatmári-Tóth *et al*., 2020].

### 2.7 Immunofluorescent Staining

hASCs were seeded and differentiated on Ibidi eight-well µ-slides as described in 2.3. Cells were washed once with PBS and fixed by 4% paraformaldehyde, followed by permeabilization with 0.1% saponin and blocking with 5% skimmed milk. Primary antibody incubations were kept overnight with anti-TOM20 (1:75, WH0009804M1) and anti-LC3 (1:200, Novus Biologicals, NB100-2220). Secondary antibody incubation was for 3 h with Alexa Fluor 647 goat anti-mouse IgG (1:1000, Thermo Fisher Scientific, A21236) and Alexa Fluor 488 goat anti-rabbit IgG (1:1000, Thermo Fisher Scientific, A11034). Propidium iodide (PI, 1.5 μg/mL, 1 h) was used for nuclei labeling. Images were obtained with an Olympus FluoView 1000 (Olympus Scientific Solutions, Tokyo, Japan) confocal microscope and FluoView10-ASW (Olympus Scientific Solutions) software version 3.0, as previously described [Szatmári-Tóth *et al*., 2020; Vámos *et al*., 2022]. LC3 and TOM20 immunostaining images were converted to binary form, followed by processing with FIJI. The LC3 punctae count was determined by size (pixel^2^) 50–infinity AU with circularity 0–1 AU. Fragmented mitochondria were analyzed from the binary TOM20 immunostaining images with size (pixel^2^) 0– 100 AU and circularity 0–1 AU. The optimum size values for the LC3 punctae and fragmented mitochondria were determined based on an analysis of all immunostaining images and manual verification of the counting accuracy by checking the outlines of counts. Both LC3 punctae and fragmented mitochondria content were normalized to per nucleus for individual images. Co-localization of LC3 and TOM20 was evaluated by calculation of the Pearson correlation coefficient (PCC) [Szatmári-Tóth *et al*., 2020; Vámos *et al*., 2022]. 30 cells per 3 donors were recorded and analyzed.

### 2.8 Quantification of amino acid fluxes of adipocytes

Supernatants of the cells were collected at the end of the differentiation process and examined as previously described [Guba *et al*., 2022; Nokhoijav *et al*., 2022]. Briefly, the media were filtered using a 3 kDa filter (Pall Corporation, Port Washington, NY, USA) and 10 µl of this filtrate was derivatized with AccQ·Tag Ultra Derivatization Kit (Waters, Milford, MA, USA). Chromatographic separation was executed on H-class UPLC (Waters) using AccQ·Tag Ultra Column (2.1 × 100 mm), AccQ·Tag Eluent A and B, and gradient was ensured by the AccQ·Tag Ultra Chemistry Kit (Waters). Amino acid derivatives were detected at 260 nm in the PDA detector of the UPLC. Detection of the amino acid concentrations were calculated by Empower software (Waters) using a 7-point calibration curve. Flux of amino acids into or from adipocytes was calculated by comparing concentration differences measured in the conditioned media, which was administered to the cells at day 25 and collected at the end point of 28 days differentiation, and the unconditioned medium. The concentration of amino acids was normalized to the number of cells as described previously [Arianti *et al*., 2021].

### 2.9 Determination of Real-Time Cellular Oxygen Consumption (OCR) and Extracellular Acidification Rate (ECAR)

Cells were seeded and differentiated on XF96 assay plates (Seahorse Biosciences, North Billerica, MA, USA) to white, active beige, or inactive beige using the protocols as described in 2.3. The OCR and ECAR were measured with XF96 oximeter (Seahorse Biosciences). Dibutyryl-cAMP stimulated OCR and ECAR, etomoxir-resistant (ETO-R) OCR, and stimulated proton leak OCR were quantified using previously utilized protocols [Arianti *et al*., 2021; Nagy *et al*., 2022]. 10 µM antimycin A was used for baseline collection (measuring non-mitochondrial respiration). The OCR was normalized to protein content of each well.

### 2.10 Statistical Analysis

All results are expressed as mean ± SD. The normality of the obtained data was tested by Shapiro-Wilk test. Datasets with a normal distribution was analyzed using one-way ANOVA with a Tukey’s post hoc test. Data was analyzed and visualized by GraphPad Prism 8 Software.

## 3. Results

### 3.1 Active beige adipocytes derived from abdominal SC exert high browning capacity

Primarily, RNA-seq analysis was performed to investigate the global gene expression patterns of the three types of differentiated SC adipocytes, white, active, and inactive beige (see Methods for their differentiation protocol). We found that general adipocyte markers, such as *SLC2A4, FABP4, LPL, ADIPOQ, AGPAT2, PLIN1, LEP*, and *LEPR* were not expressed differentially among the three types of adipocytes (Fig. 1A) suggesting that their differentiation rate was similar. The thermogenic markers, such as *UCP1, ELOVL3, PGC1a, CIDEA, CITED1, AQP3, GK, CKMT1a/b*, and *PM20D1* had higher expression in active beige as compared to white or inactive beige adipocytes (Fig. 1B). Next, we subjected our transcriptomic data to open source webtools to estimate brown adipocyte content by BATLAS [Perdikari *et al*., 2018] and browning capacity by ProFAT [Cheng *et al*., 2018]. We did not find significant differences in brown adipocyte content (Fig. 1C), however, active beige adipocytes showed higher browning capacity score as compared to white or inactive beige ones (Fig, 1D). The expression pattern of BATLAS and ProFAT markers are displayed as heatmaps on Fig. 1E and F. We found that 211 and 147 genes had higher expression in the comparison of active beige vs white and active beige vs inactive beige, respectively; out of those, 100 genes were common (Fig. 1G, top panel and Supplementary Tables 2 and 3). Among the commonly highly expressed 100 genes, thermogenic markers, such as *GK, PM20D1, PLIN5, CITED1*, and *AQP3* were found (Fig. 1H). Interestingly, *SLC7A10*, encoding ASC1, which was described as an important transporter during thermogenic activation [Arianti *et al*., 2021; Jersin *et al*., 2021] was also commonly upregulated in both comparisons (Fig. 1H). 248 and 226 genes had lower expression in the comparison of active beige vs white and active beige vs inactive beige, respectively, and 164 genes had commonly lower expression (Fig. 1G, bottom panel, Supplementary Tables 2 and 3). We did not find any DEGs when we compared the gene expression profile of white and inactive beige adipocytes. We also analyzed the mitophagy rate and mitochondrial morphology by co-immunostaining of microtubule-associated protein 1 light chain 3 (LC3) and translocase of outer mitochondrial membrane 20 (TOM20) [Szatmari-Toth *et al*., 2020] (Supplementary Fig. 1A). Confocal images were used to quantify the co-localization of LC3 and TOM20 by measuring the correlation between pixel intensities of two detection channels [Szatmari-Toth *et al*., 2020]. We observed lower LC3 puncta counts per cell and PCC values in active beige as compared to white or inactive beige adipocytes (Supplementary Fig. 1B-C). We also found that active beige adipocytes had higher amounts of fragmented mitochondria, which were shown to support uncoupled respiration and enhanced energy expenditure [Pisani *et* al., 2018], as compared to white or inactive beige cells (Fig. 1I). Altogether, these data suggests that thermogenesis-related genes were upregulated, the mitophagy rate was lowered, and mitochondria were more fragmented when human abdominal SC adipocytes were activated for thermogenesis.

**Figure 1.**
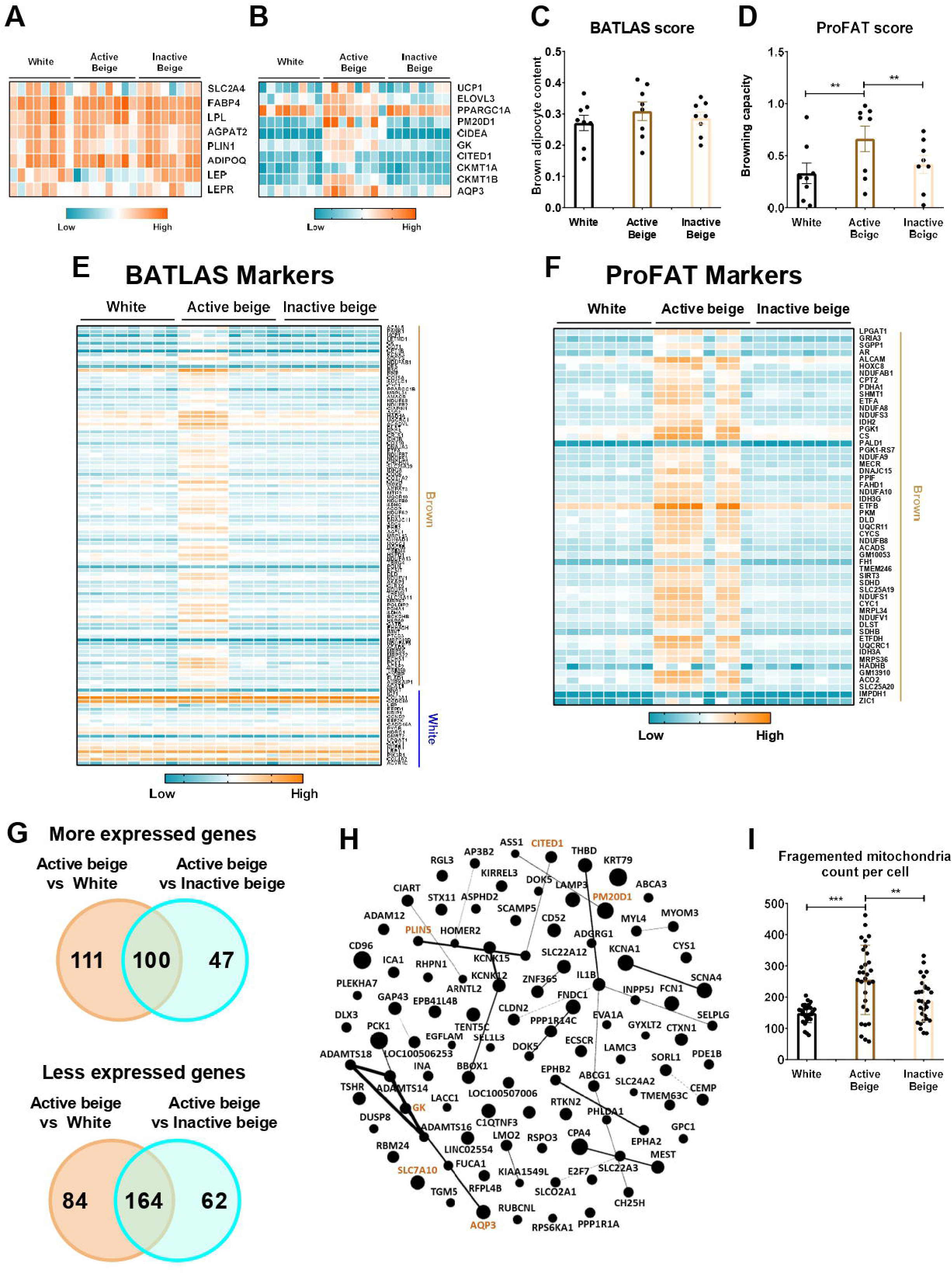
Differentially expressed genes and thermogenic capacity of human abdominal subcutaneous adipocytes. (A) Heatmap displaying the expression of general markers of adipocytes. (B) Heatmap displaying the expression of brown/beige adipocyte markers. (C) Brown adipocyte content quantified by BATLAS open source webtool, n=8. (D) Browning capacity quantified by ProFAT open source webtool, n=8. (E) Heatmap displaying the expression pattern of BATLAS markers. (F) Heatmap displaying the expression of ProFAT markers. (G) Venn diagram displaying the numbers of more (top panel) and less (bottom panel) expressed genes in comparison of active beige vs white and active beige vs inactive beige adipocytes. (H) Overlapping upregulated genes between active beige vs white and active beige vs inactive beige in interactome map generated by Gephi. The size of the nodes correlates with fold change in the expression values. Gene names marked in brown are known thermogenic markers. (I) Quantification of fragmented mitochondria count per cell, n=30 cells of 3 donors. Statistical analysis by ANOVA. **p<0.01 and ***p<0.001.

### 3.2 Active beige adipocytes carrying *FTO* rs1421085 obesity-risk alleles had lower brown adipocyte content and expressed lower level of genes involved in metabolic pathways

Next, we investigated whether *FTO* rs1421085 SNP affected the browning capacity of human abdominal SC adipocytes, which were differentiated into white, active, or inactive beige. We genotyped the hASCs for *FTO* rs1421085 SNP by using SNP genotyping assay and obtained the allelic discrimination plot (Fig. 2A). Then, we selected samples from 4-4 individuals with either homozygous TT (risk-free) or CC (obesity-risk) genotypes for further analysis. We found that active beige adipocytes carrying *FTO* obesity-risk alleles exerted lower brown adipocyte content estimated by BATLAS [Perdikari *et al*., 2018], but no effect of the allelic distribution was observed in case of white or inactive beige adipocytes (Fig. 2B). We also found that the risk-free genotype carrier active beige adipocytes had higher BATLAS and ProFAT [Cheng *et al*., 2018] scores as compared to white ones that carried the same TT genotype (Fig. 2B-C). Interestingly, active beige adipocytes carrying *FTO* obesity-risk genotype showed similar estimated brown adipocyte content and browning capacity as compared to white adipocytes (Fig. 2B-C) suggesting that active beige differentiation could not overcome the browning inhibitory effect of the CC alleles. Active beige adipocytes with *FTO* risk-free genotype expressed the BATLAS marker genes at the highest level, whereas ones with the obesity-risk allele carriers expressed them at a level similar to those observed in the white or inactive beige adipocytes (Fig. 2D). The expression of ProFAT markers was high in risk-free carrier active beige adipocytes and showed large donor variability in the obesity-risk allele carrier ones (Fig. 2E).

**Figure 2.**
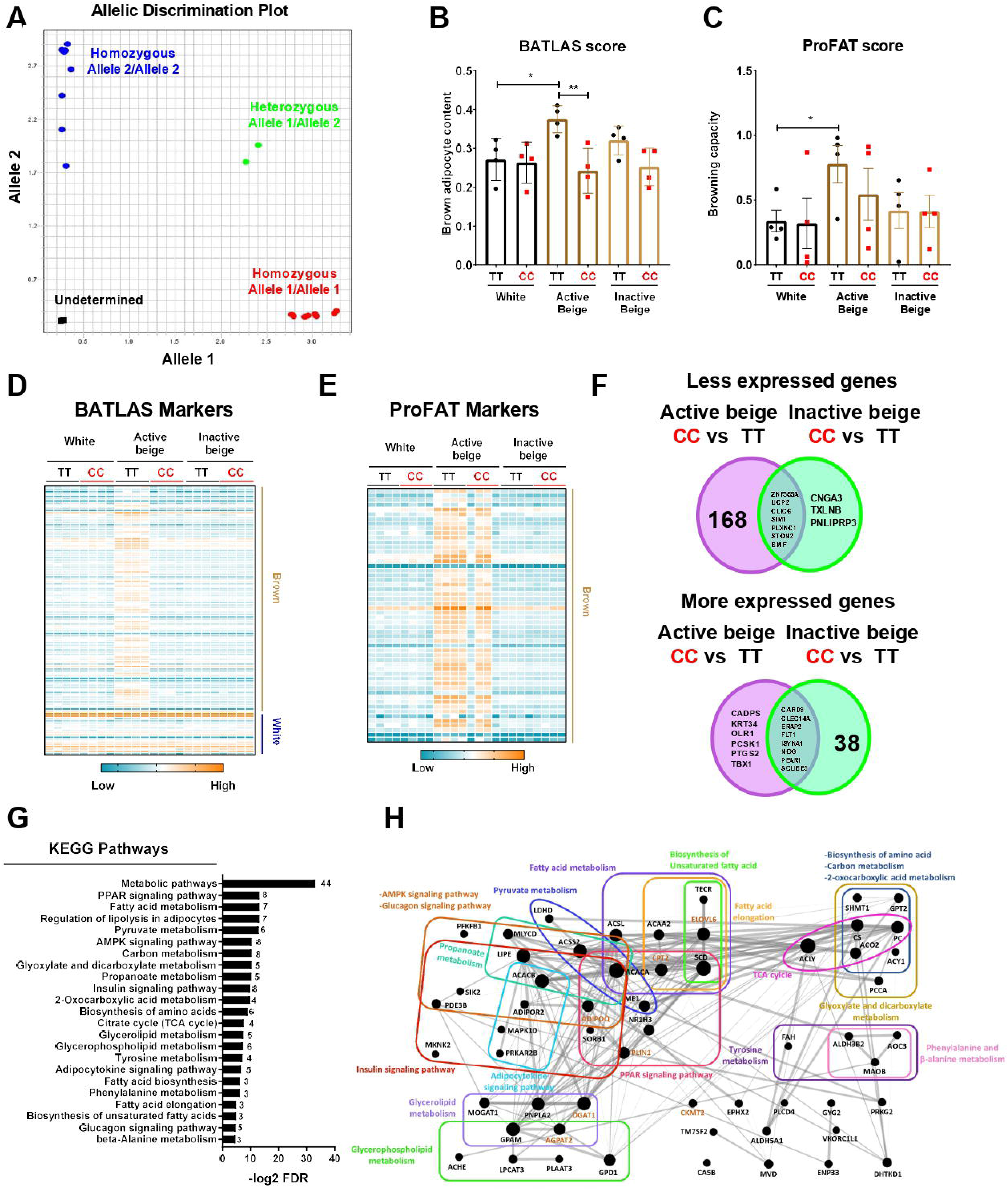
The effect of *FTO* rs1421085 SNP on the thermogenic capacity and gene expression pattern of differentiated abdominal subcutaneous (SC) adipocytes. (A) Allelic distribution of *FTO* rs1421085 SNP. Blue: SC progenitors with TT alleles, red: SC progenitors with CC alleles, green: heterozygous SGBS adipocytes, black: no template negative control. (B) Brown adipocyte content quantified by BATLAS, n=4 of each genotype. (C) Browning capacity quantified by ProFAT, n=4 of each genotype, *p<0.05, **p<0.01, statistical analysis by ANOVA. (D) Heatmap displaying the expression pattern of BATLAS markers. (E) Heatmap displaying the expression of ProFAT markers. (F) Venn diagram displaying the numbers of less (top panel) and more (bottom panel) expressed genes in comparison of *FTO* rs1421085 CC vs TT in both active and inactive beige differentiated adipocytes. (G) Overrepresented pathways which are less expressed in active beige differentiated-adipocytes carrying *FTO* CC/obesity-risk variant as compared to TT/risk-free variant. Numbers on the right side indicate the number of genes involved in the pathways. (H) Genes involved in less expressed pathways shown in figure (G) in interactome map generated by Gephi. The size of the nodes correlates with fold change of the expression values. Gene names marked in brown are known thermogenic markers.

A total of 175 genes including thermogenic markers (*UCP2, CKMT2*, and *CITED1*) and 5 BATLAS markers (*PPARGC1B, ACO2, ACSF2, NNAT*, and *DMRT2*) were expressed less in active beige adipocytes carrying *FTO* obesity-risk variant as compared to risk-free carriers (Figure 2F, top panel, Supplementary Table 4). Only 10 genes (7 of them were common in both comparisons) were expressed at a lower extent in CC as compared to TT carrier inactive beige adipocytes (Fig. 2F, top panel, Supplementary Table 5). We found only 14 and 46 genes (8 of them were common in both comparisons) which were expressed more in active or inactive beige adipocytes, respectively, carrying the *FTO* obesity-risk as compared to the risk-free variant (Fig. 2F, bottom panel, Supplementary Tables 4 and 5). In white adipocytes, we did not find any DEGs which was affected by the *FTO* rs1421085 SNP. Next, we investigated the gene expression pathways affected by the *FTO* rs1421085 SNP and found that genes, which were less expressed in active beige adipocytes that carried the obesity-risk genotype, were overrepresented in several pathways, such as metabolic, PPAR signaling, lipolysis, fatty acid metabolism, or TCA cycle (Fig. 2G-H). More expressed genes in active beige adipocytes with CC alleles were not significantly overrepresented in any of the pathways. We did not find any overrepresented pathway with respect to the DEGs found in inactive beige adipocytes either.

Because we observed significant effects of *FTO* rs1421085 SNP on the gene expression pattern of active beige SC adipocytes, we also compared the transcriptomic data of active beige and white or inactive beige adipocytes that carried *FTO* risk-free or obesity-risk genotypes, respectively. We found approximately four times more genes that were expressed in a larger extent in active beige vs white adipocytes in the case of *FTO* risk-free genotype as compared to obesity-risk carrier samples. Only 25 genes were overlapping in the two aforementioned comparisons (Supplementary Fig. 2A, Supplementary Tables 6 and 7). Approximately similar number of genes were expressed less in active beige as compared to white adipocytes carrying *FTO* risk-free or obesity-risk alleles, of which almost half of them were common (Supplementary Fig. 2B, Supplementary Table 6 and 7). The PPAR signaling pathway was the only one in which more expressed genes in active beige as compared to white adipocytes carrying *FTO* risk-free variant were overrepresented. The less expressed genes in active beige adipocytes as compared to white ones were overrepresented in several pathways, such as axon guidance or longevity regulating only in *FTO* obesity-risk genotype carriers (Supplementary Fig. 2C). A similar trend was observed when more or less expressed genes were analyzed in the comparison of active vs inactive beige adipocytes carrying *FTO* risk-free or obesity-risk variants, respectively (Supplementary Fig. 2D-E, Supplementary Tables 8 and 9). Genes expressed in a lower extent in active as compared to inactive beige adipocytes were overrepresented in several pathways, such as AMPK signaling and TGF-beta signaling only in *FTO* obesity-risk allele carriers (Supplementary Fig. 2F). When adipocytes carried the risk-free genotype, we did not find any overrepresented pathway in comparison of active with white or inactive beige adipocytes. These results suggest that *FTO* rs1421085 SNP only affects the gene expression profile, particularly that of the thermogenesis-related genes, in active beige but not in white or inactive beige adipocytes. In addition, the applied differentiation protocols resulted in more pronounced differences in the gene expression patterns of adipocytes with *FTO* risk-free alleles which suggest their significant browning potential when thermogenic cues are constantly present.

### 3.3 Thermogenic marker genes were less expressed in active beige adipocytes carrying *FTO* obesity-risk alleles

Since we observed that the allelic discrimination at *FTO* rs1421085 SNP influences the expression of thermogenic and BATLAS markers, we went further to investigate the expression of thermogenic genes at mRNA and protein levels in abdominal SC adipocytes. Our results showed that the mRNA expression of *UCP1* was higher in active beige as compared to white or inactive beige adipocytes with *FTO* risk-free alleles, however, this difference was not observed in obesity-risk carriers (Fig. 3A). At the protein level, we observed that active beige adipocytes expressed more UCP1 as compared to white or inactive beige adipocytes regardless the *FTO* rs1421085 genotypes, however, less UCP1 protein amount was detected in obesity-risk than in risk-free allele carrier active beige adipocytes. (Fig. 3B). Other thermogenic genes, such as *UCP2, PM20D1, CIDEA, CITED1, CKMT1* and *2, CPT2*, and *PLIN1* were also expressed higher in active beige adipocytes carrying risk-free alleles as compared to white or inactive beige adipocytes with the same TT variant, however, we did not observe these differences in *FTO* obesity-risk carrier samples (Fig. 3D-J). As compared to risk-free carriers, active beige adipocytes with *FTO* obesity-risk genotype had lower expression of these thermogenic genes and also that of the neurotrophic factor *S100b*, which stimulates sympathetic axon growth and plays important role in BAT innervation through calsyntenin (CLSTN) 3β axis [Zeng *et al*., 2019] (Fig. 3A-K). These results are in accordance with our RNA-seq data suggesting the critical importance of *FTO* rs1421085 SNP in active beige adipocytes and the compromised effect of active beige differentiation protocol in the presence of obesity-risk alleles.

**Figure 3.**
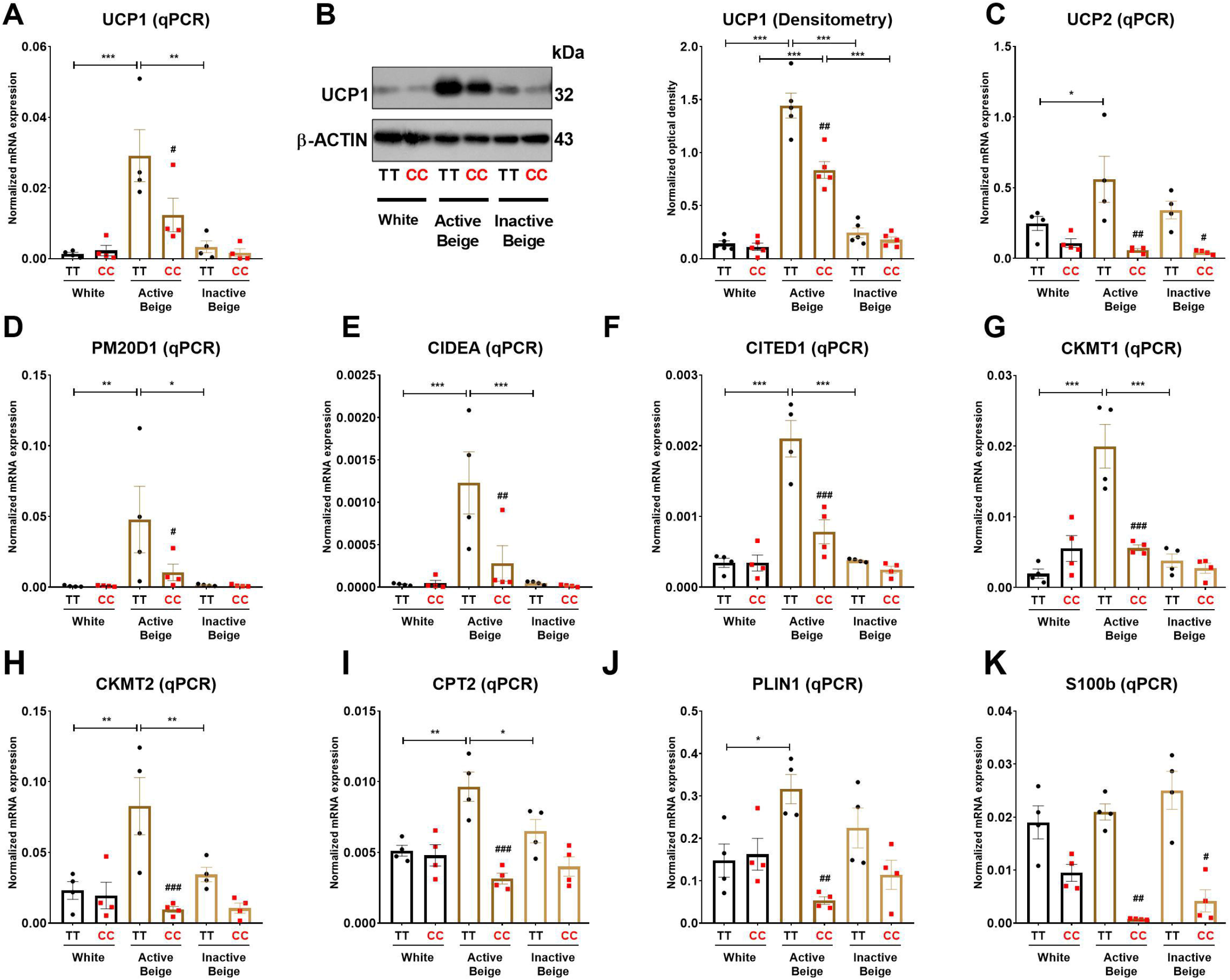
The effect of the differentiation protocols and alleles at *FTO* rs1421085 on the expression of thermogenesis markers in differentiated abdominal subcutaneous adipocytes. (A-B) The mRNA (A) and protein (B) expression of UCP1 measured by qPCR or immunoblotting. (C-K) The mRNA expression of *UCP2, PM20D1, CIDEA, CITED1, CKMT2* and *2, CPT2, PLIN1* and *S100B* was analyzed by RT-qPCR. Statistical analysis was performed by ANOVA, n=4 of each genotypes, */#p<0.05, **/##p<0.01, and ***/###p<0.001. *analysis was performed to compare the effect of the applied differentiation protocol in the same genotype. #analysis was performed to compare *FTO* rs1421085 TT and CC genotypes within the same differentiation protocol.

### 3.4 Active beige adipocytes with *FTO* obesity-risk genotype expressed less amount of mitochondrial complex subunits and had lower proton leak respiration

Having observed that *FTO* rs1421085 SNP affected the expression of thermogenic genes, we investigated whether the expression of mitochondrial complex subunits and cellular respiration were also suppressed in adipocytes with obesity-risk alleles. We found that active beige adipocytes carrying *FTO* risk-free genotype had higher amounts of mitochondrial complex subunits I, II, and IV as compared to white or inactive beige adipocytes carrying the same TT genotype (Fig. 4A-C, E). However, no difference was found between the three types of differentiation programs when adipocytes carried the obesity-risk variant (Fig. 4A-C, E). Active beige adipocytes with *FTO* obesity-risk alleles had lower expression of mitochondrial complex subunits I, II, and IV as compared to the risk-free carriers (Fig. 4A-C, E). We observed a similar but statistically not significant trend in the case of mitochondrial complex subunit III (Fig. 4D), while the expression of complex V subunit was similar for all types of adipocytes regardless the *FTO* rs1421085 genotype (Fig 4F). Next, we measured the cellular respiration of the three types of adipocytes carrying *FTO* risk-free or obesity-risk alleles. In accordance with the mitochondrial complex subunit expression, we found that active beige adipocytes carrying risk-free genotype had higher respiration (at both basal and maximal stimulated conditions), proton leak respiration, and extracellular acidification as compared to white or inactive beige adipocytes, but this difference was not pronounced when the adipocytes carried the risk variant (Fig. 4G-H). Active beige adipocytes with *FTO* obesity-risk variant showed lower cellular respiration, especially proton leak respiration which reflects UCP1-dependent heat production, and extracellular acidification, which associates with glycolytic activity, as compared to risk-free carriers (Fig. 4G-H). Intriguingly, the effect of *FTO* obesity-risk alleles on cellular respiration was observed in active beige but not in white or inactive beige adipocytes highlighting its exclusive effect in human abdominal SC adipocytes only when they are activated for thermogenesis.

**Figure 4.**
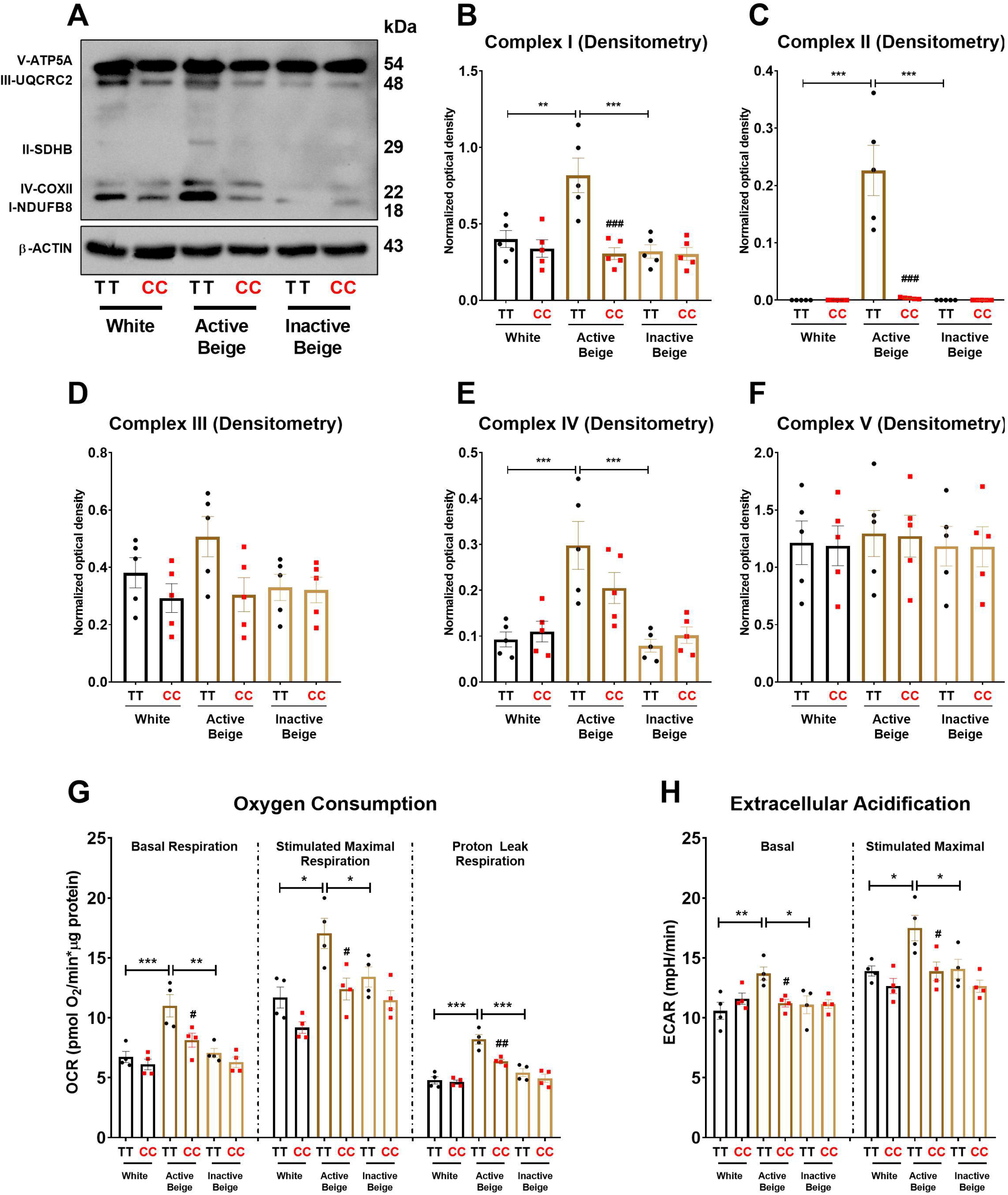
The effect of the differentiation protocols and alleles at *FTO* rs1421085 on the expression of mitochondrial complex subunits, oxygen consumption, and extracellular acidification. (A-F) Protein expression of mitochondrial complex subunits detected by immunoblotting. (G-H) Oxygen consumption at basal, dibutyryl-cAMP stimulated maximal, and proton leak respiration (G) and extracellular acidification (H) were quantified in white, active beige, and inactive beige adipocytes carrying *FTO* risk-free or obesity risk genotypes by Seahorse extracellular flux analysis. Statistical analysis was performed by ANOVA, n=4 of each genotypes, */#p<0.05, **/##p<0.01, and ***/###p<0.001. *analysis was performed to compare the effect of the applied differentiation protocol in the same genotype. #analysis was performed to compare *FTO* rs1421085 TT and CC genotypes within the same differentiation protocol.

### 3.5 Adipocytes carrying *FTO* obesity-risk genotype consume lower amounts of neutral amino acids when activated for thermogenesis

Active thermogenic adipocytes utilize metabolic substrates, such as carbohydrates, fatty acids, or amino acids to generate heat [Onogi and Ussar, 2022]. ASC1, which is encoded by *SLC7A10*, plays an important role in mediating amino acid consumption in human adipocytes derived from abdominal SC and deep neck regions [Jersin *et al*., 2021; Arianti *et al*., 2021]. To evaluate the preferable energy sources during thermogenic activation, we monitored the oxygen consumption of adipocytes upon etomoxir (inhibitor of carnitine palmitoyltransferase-1) administration. ETO-R respiration, which reflects the activity of carbohydrate and amino acid oxidation [Nagy *et al*., 2022], was higher in active beige adipocytes with *FTO* risk-free genotype than in white or inactive beige adipocytes carrying the same TT genotype. Active beige adipocytes with *FTO* obesity-risk alleles had lower level of ETO-R oxygen consumption as compared to risk-free carriers (Fig. 5A). These observations suggest less pronounced carbohydrate and/or amino acid utilization in active beige adipocytes of CC carriers at *FTO* rs1421085.

**Figure 5.**
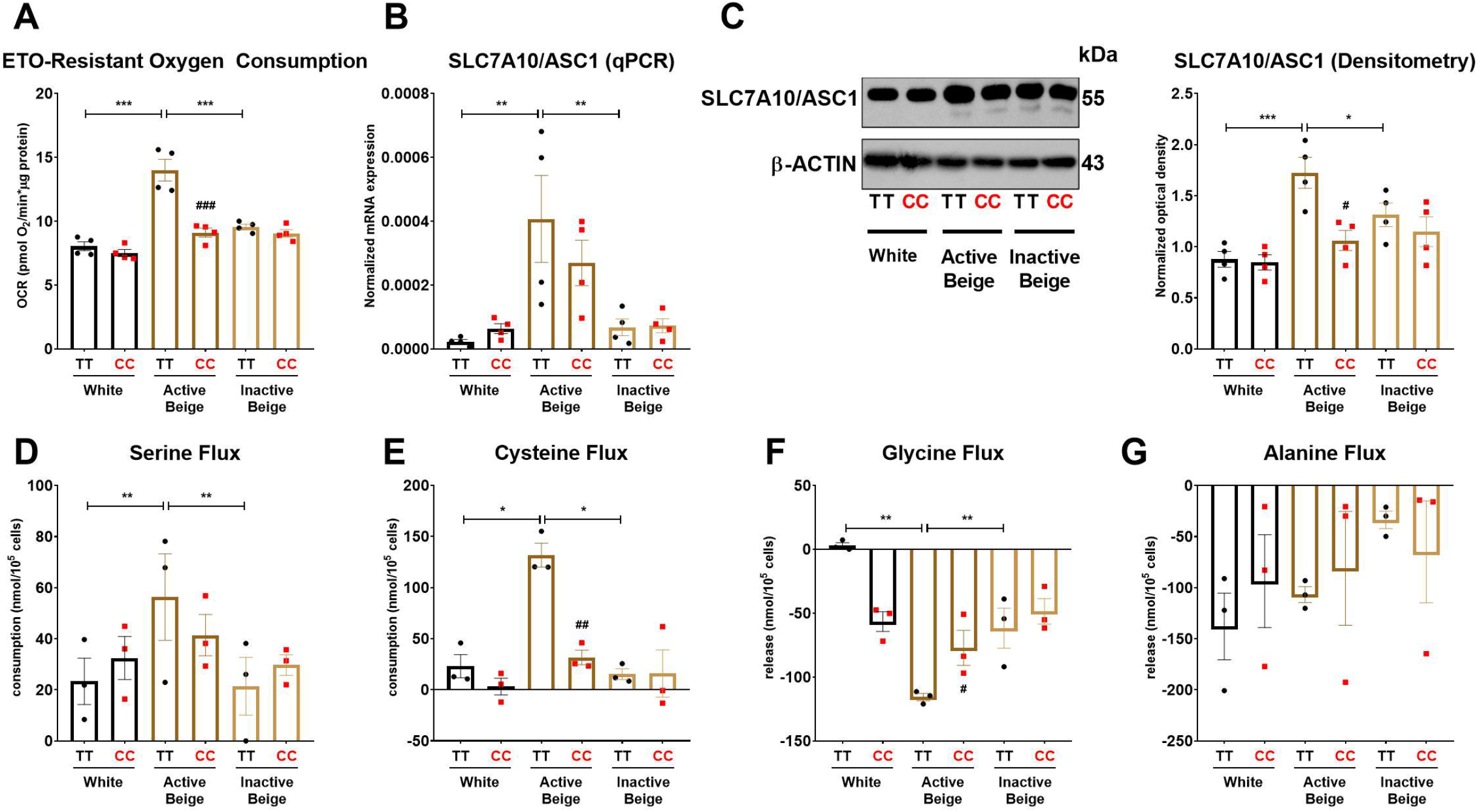
The effect of the differentiation protocols and alleles at *FTO* rs1421085 on the ASC1-mediated amino acid flux of differentiated abdominal subcutaneous adipocytes. (A) Etomoxir-resistant oxygen consumption was quantified in white, active beige, and inactive beige adipocytes carrying risk-free or obesity-risk genotypes by Seahorse extracellular flux analysis. (B-C) mRNA (B) and protein (C) expression of *SLC7A10*/ASC1 by RT-qPCR and immunoblotting, n=4 of each genotype. (D-G) Amino acids flux measured in the conditioned media of differentiated abdominal subcutaneous adipocytes, n=3 of each genotype. Statistical analysis was performed by ANOVA. */#p<0.05, **/##p<0.01, and ***/###p<0.001. *analysis was performed to compare the effect of the applied differentiation protocol in the same genotype. #analysis was performed to compare *FTO* rs1421085 TT and CC genotypes within the same differentiation protocol.

Because we found *SLC7A10* (encoding alanine-serine-cysteine transporter, ASC-1) as a DEG among the most abundantly expressed genes in active beige adipocytes with *FTO* risk-free genotype (Fig. 1H, Supplementary Table 6), we decided to investigate the effect of the applied differentiation protocols and *FTO* rs1421085 SNP on the expression of ASC1 and the consumption of ASC1 cargos by the adipocytes. We found that active beige adipocytes with risk-free alleles expressed higher mRNA level of *SLC7A10* as compared to white or inactive beige ones that carried the same TT variant (Fig. 5B), which could be confirmed at ASC1 protein level (Fig. 5C). The presence of the *FTO* rs1421085 SNP resulted in lower expression of *SLC7A10* in active beige adipocytes; this effect was statistically significant at protein level but not at mRNA level (Fig. 5B-C). Next, we measured the consumption of amino acids in the conditioned media obtained from the three types of differentiated adipocytes with CC or TT alleles, respectively. In the case of adipocytes with *FTO* risk-free genotype, we found that active beige ones consumed higher amounts of serine (Fig. 5D) and cysteine (Fig. 5E) and released more glycine as compared to white or inactive beige adipocytes (Fig. 5F). In accordance with the aforementioned gene expression and ETO-R oxygen consumption results, we did not observe any differences in the fluxes of these amino acids between the three types of differentiation protocols in adipocytes with obesity-risk alleles. The applied differentiation programs did not affect the release of alanine regardless of the *FTO* allele status (Fig. 5G). Active beige adipocytes with obesity-risk genotype consumed lower amount of cysteine as compared to risk-free carriers (Fig. 5E), but significant effect of the SNP was not observed on serine consumption (Fig. 5D) suggesting that other amino acid transporters might compensate for the reduced expression of ASC1.

Our RNA-seq data showed that serine hydroxymethyltransferase (SHMT) 1, which catalyzes the conversions of L-serine to glycine and tetrahydrofolate (THF) to 5,10-methylene-THF (5,10-CH_2_-THF), was expressed lower in active beige adipocytes with *FTO* obesity-risk as compared to risk-free carriers (Supplementary Table 4). We validated our RNA-seq data of SHMT1 expression by RT-qPCR (Fig. 6A) and immunoblotting (Fig. 6B). We also found that active beige adipocytes had higher protein content of SHMT1 as compared to white or inactive beige adipocytes in the presence of the *FTO* risk-free variant, but this difference was not observed in obesity-risk carrier samples (Fig. 6B).

**Figure 6.**
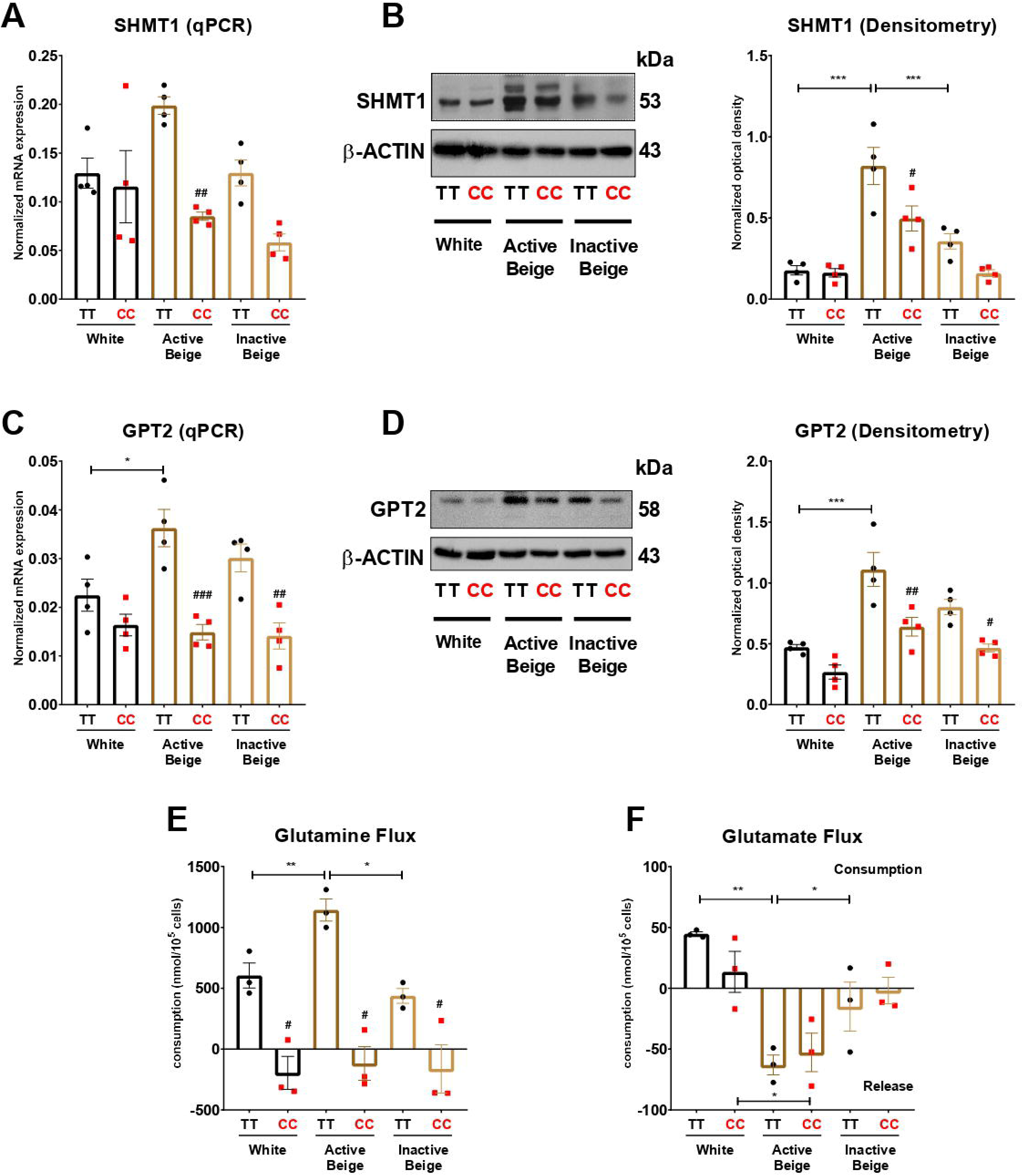
The effect of differentiation protocols and alleles at *FTO* rs1421085 on the expression of SHMT1 and GPT2, and glutamine-glutamate flux of differentiated abdominal subcutaneous adipocytes. (A-B) mRNA (A) and protein (B) expression of SHMT1 by RT-qPCR and immunoblotting, n=4 of each genotype. (C-D) mRNA (C) and protein (D) expression of GPT2 by RT-qPCR and immunoblotting, n=4 of each genotype. (E-F) Glutamine (E) and glutamate (F) flux measured in the conditioned media of differentiated abdominal subcutaneous adipocytes, n=3 of each genotypes. Statistical analysis was performed by ANOVA. */#p<0.05, **/##p<0.01, and ***/###p<0.001. *analysis was performed to compare the effect of the applied differentiation protocol in the same genotype. #analysis was performed to compare *FTO* rs1421085 TT and CC genotypes within the same differentiation protocol.

According to the RNA-seq data, active beige adipocytes carrying obesity-risk genotype expressed lower mRNA level of glutamic pyruvic transaminase (GPT) 2 as compared to risk-free allele carriers (Supplementary Table 4). In the case of adipocytes with *FTO* risk-free alleles, active beige cells expressed more GPT2 both at mRNA (Fig. 6C) and protein level (Fig. 6D) as compared to white ones. We also observed lower expression of GPT2 in active and inactive beige adipocytes with FTO obesity-risk genotype as compared to risk-free allele carriers (Fig. 6C-D). Active beige adipocytes carrying risk-free genotypes also consumed higher amount of glutamine as compared to white or inactive beige adipocytes carrying the same TT genotype, but no difference was observed among the three types of differentiation protocols in the case of the samples with obesity-risk alleles. On the contrary to risk-free carrier cells, adipocytes with *FTO* obesity-risk genotype did not consume glutamine irrespective to the applied differentiation protocols (Fig. 6E). In contrast, we found that active beige adipocytes released higher amount of glutamate as compared to inactive ones, while white adipocytes rather consumed that amino acid. The consumption of glutamate did not depend on the allelic discrimination at *FTO* rs1421085 locus (Fig. 6F). In the case of adipocytes with TT alleles, active beige cells consumed less aspartate than white or inactive beige ones, however, the consumption of other amino acids was not affected by either the applied protocols or the *FTO* genotypes (Supplementary Fig. 3).

## 4 Discussion

Abdominal fat is classified into SC and intraabdominal fat, which is mainly composed of visceral or intraperitoneal WAT [Marin *et al*., 1992; Wajchenberg, 2000]. Several studies reported that accumulation of visceral WAT is strongly associated with the risk of metabolic disorders [Matsuzawa *et al*., 1995; Fujimoto *et al*., 1994; Banerji *et al*., 1999], whereas other studies claimed that abdominal SC WAT may possess a protective role [McLaughlin *et al*., 2011; Patel and Abate, 2013]. A more recent study using an elegant PET/CT technique, Leitner *et al*. (2017) reported that human BAT or brownable adipose tissue can be found in cervical, supraclavicular, axillary, mediastinal, paraspinal, and abdominal depots. In this study, we performed RNA-seq on human abdominal SC derived adipocytes with *FTO* rs1421085 risk-free or obesity-risk genotypes, which were differentiated by applying three types of protocols: white, active, or inactive beige. Irrespective to the *FTO* genotypes, we found that active beige adipocytes exerted greater thermogenic potential, marked by higher expression of thermogenic genes (Fig. 1B) and browning capacity quantified by ProFAT (Fig. 1D), as compared to white or inactive beige cells. Our results suggest that human abdominal SC adipocytes have a significant thermogenic potential when they are activated using active beige differentiation protocol driven by a PPARγ agonist and adrenergic stimulation. However, this potential subsides when adipocytes are inactivated through beige to white transition. This is in line with our previous study which has reported that beige to white transition leads to increased mitophagy resulting in the appearance of white-like phenotype and reduced thermogenesis [Vámos *et al*., 2022]. Other studies also revealed that beige adipocytes in mice gradually lose their thermogenic morphology and capacity after the external stimuli, such as β-adrenergic agonist or cold exposure, were withdrawn [Altshuler-Keylin *et al*., 2016; Rosenwald *et al*., 2013]. The loss of thermogenic characteristics in murine beige adipocytes was coupled with increased mitophagy, which was mediated by parkin [Lu *et al*., 2018; Sarraf and Youle, 2018]. The comparison of active beige and white or inactive beige adipocytes found 100 and 164 genes, which were commonly more and less expressed, respectively, in active beige adipocytes. Notably, several well-known thermogenesis markers, such as *CITED1, PM20D1, PLIN5, GK*, and *AQP3* were commonly upregulated in active beige as compared to white or inactive beige adipocytes. No DEGs were found in the comparison of white and inactive beige adipocytes indicating a high similarity of the gene expression profiles in these two differentiation programs.

Dibutyryl-cAMP is extensively used to mimic *in vivo* thermogenesis due to its ability to penetrate the cell membrane [Cypess *et al*., 2013]. In contrast to cAMP, which can be hydrolyzed by phosphodiesterase (PDE), dibutyryl-cAMP is resistant to degradation by PDE [Jarett *et al*., 1974; Blecher *et al*., 1971]. Dibutyryl-cAMP directly inhibits PDE in a competitive manner resulting in a significant elevation of endogenous cAMP levels [Jarett *et al*., 1974]. cAMP activates protein kinase A (PKA), which phosphorylates various proteins and initiate consecutive cascades of additional protein kinases [Daniel *et al*., 1998]. More recently, a role has emerged for PKA in the regulation of gene transcription [Daniel *et al*., 1998; London and Stratakis, 2022]. When we administered dibutyryl-cAMP in the middle of active and inactive beige differentiation programs (at day 14 for 4 hours) we found that the effect of the compound on the thermogenic gene expression was sustained until the end of the differentiation of active beige, but not in inactive beige (undergoing beige to white transition) adipocytes. This suggests that the effect of dibutyryl-cAMP can be maintained for a long period of time in beige adipocytes. Activation of cAMP response element binding protein (CREB) is one of the most studied links between PKA and gene expression regulation [Daniel *et al*., 1998]. CREB binding protein (CBP) associates specifically with CREB when it is phosphorylated at serine 133 and enhances cAMP-induced transcription [Kwok *et al*., 1994]. Our RNA-seq data showed that the expression of Cbp/p300 interacting transactivator with Glu/Asp rich carboxy-terminal domain 1 (CITED1) was higher in active beige as compared to white or inactive beige adipocytes (Fig. 1H). CITED1, a beige-selective transcription factor, is highly expressed in human thermogenic adipocytes derived from various depots, including SC supraclavicular, posterior mediastinal, retroperitoneal, intraabdominal, mesenteric, and inguinal [Sharp *et al*., 2012]. Daniel *et al*., (1998) described that cAMP increased the mRNA level of PDE, although the molecular mechanism of this regulation remained elusive. We also found that the expression of *PDE1B* was high in active beige adipocytes (Fig. 1H) suggesting active cAMP-driven signaling. In another set of experiments, we found that the mRNA expression of *UCP1* was higher in active beige as compared to regularly differentiated beige adipocytes (28 days differentiation driven by rosiglitazone without dibutyryl-cAMP administration at day 14) (Supplementary Fig. 4) suggesting that dibutytyl-cAMP treatment at the middle of beige differentiation program enhances the thermogenic capacity of abdominal subcutaneous adipocytes at a sustained manner.

We also evaluated the effect of rs1421085 T-to-C SNP of the *FTO* gene, which interrupts a conserved motif for ARID5B repressor, resulting in elevated expression of IRX3 and IRX5 during the early stage of adipocyte differentiation. As the consequence, the commitment of the cells diverts from beige towards the white program and lipid storage increases [Claussnitzer *et al*., 2015]. When the gene expression profiles of the three types of adipocytes were analyzed by segregating the *FTO* rs1421085 risk-free (TT) and obesity-risk (CC) allele carrier samples, intriguingly, we found that the SNP affected the gene expression profile, in particular the expression of thermogenic markers (*CITED1, CIDEC, PLIN1, LIPE, CKMT2*, and *S100b*), in active beige adipocytes, but not in white or inactive beige adipocytes (Fig. 2F-H). CIDEC, PLIN1, and LIPE are lipid droplet-associated proteins, which regulate triglyceride accumulation and lipolysis [Wolin *et al*., 2006; Puri *et al*., 2007]. Decreased expression of these genes in active beige adipocytes with *FTO* obesity-risk alleles may contribute to the downregulation of lipolysis in the SC adipose tissue of affected individuals. CKMT1a/b and CKMT2 mitochondrial creatine kinases phosphorylate creatine generating phosphocreatine and contribute to UCP1-independent heat generation via creatine futile cycle [Kazak *et al*., 2015]. S100b protein plays a role in the sympathetic innervation of thermogenic adipose tissue by stimulating neurite outgrowth from sympathetic neurons [Zeng *et al*., 2019]. Reduced expression or loss of function of S100b resulted in disrupted sympathetic innervation leading to reduced thermogenesis in brown or beige adipocytes. Decreased expression of S100b in abdominal SC adipocytes with *FTO* obesity-risk carriers might partially contribute to lower thermogenic capacity in abdominal SC WAT even when the adipocytes are activated for heat production. Importantly, genes overrepresented in metabolic, especially in energy metabolism-related pathways, such as TCA cycle, lipolysis, pyruvate metabolism, and PPAR signaling were downregulated in active beige adipocytes with obesity-risk genotypes as compared to risk-free allele carriers (Fig. 2G), indicating lower energy dissipation in active beige adipocytes with CC alleles. Our findings suggest that C alleles at *FTO* rs1421085 suppress the thermogenic activation of human abdominal SC adipocytes; even long-term rosiglitazone treatment could not compensate for the effect of the obesity-risk genotype. In addition, we also observed that active beige adipocytes carrying *FTO* obesity-risk alleles exerted similar transcriptomic profiles as white or inactive beige adipocytes (Fig. 2). This is in association with our previous study, which reported that human neck derived adipocytes carrying *FTO* obesity-risk genotype had lower expression of thermogenic genes, such as *CKMT1A/B, CITED1, PPARGC1A/B*, and *CPT1B* and genes involved in respiratory electron transport, fatty acid metabolism, and the signaling by retinoic acid pathways [Tóth *et al*., 2020].

Active heat producing adipocytes utilize higher amounts of nutrients, such as glucose, fatty acids, and amino acids to provide sufficient fuel for thermogenesis and solute carrier (SLC) transporters play a crucial role in mediating the transport of these molecules [Cypess *et al*., 2009; Virtanen *et al*., 2009; Wu *et al*., 2006; Yoneshiro *et al*., 2019]. Our data showed that active beige adipocytes carrying risk-free genotype had higher ETO-R oxygen consumption that reflects carbohydrate and amino acid utilization and expressed higher level of the neutral amino acid transporter, ASC1 (encoded by *SLC7A10*) as compared to white or inactive beige adipocytes with the same TT genotypes, whereas there was no difference when the three types of adipocytes with FTO obesity-risk alleles were compared (Fig. 5A-C). ASC1 has been recently identified as a novel regulator of energy metabolism in human SC adipose tissue elevating mitochondrial respiration and preventing development of adipocyte hypertrophy and insulin resistance [Jersin *et al*., 2021]. Our previous study reported that ASC1-mediated uptake of serine, cysteine, and glycine is essential for efficient thermogenic response upon adrenergic stimulation in human neck derived adipocytes [Arianti *et al*., 2021]. The role of ASC1 in adipose tissue has been comprehensively reviewed by Jersin *et al*., (2022) highlighting its beneficial effects in enhancing mitochondrial activity and lowering reactive oxygen species production in white adipocytes. We also found that the consumption of serine and cysteine was higher in active beige as compared to white or inactive beige adipocytes with *FTO* risk-free genotype. Lower consumption of these amino acids was observed in active beige adipocytes with *FTO* obesity-risk genotype as compared to risk-free allele carriers. In addition, we observed release of less glycine by active beige adipocytes carrying obesity-risk genotypes as compared to those with risk-free alleles (Fig. 5D-F). Serine is an important metabolic source to generate one-carbon units in mammalian cells [de Koning *et al*., 2003], which are produced by both isoforms of SHMT enzymes, SHMT1 (cytosolic) and SHMT2 (mitochondrial), resulting in glycine. Our data showed that active beige adipocytes carrying *FTO* obesity-risk genotype expressed lower level of SHMT1 as compared to risk-free allele carrier ones (Fig. 6A-B) suggesting lower generation of one-carbon units in thermogenic adipocytes with *FTO* obesity-risk genotype. We also revealed that one-carbon metabolism pathway, in which SHMT1 and GPT2 participate, was less expressed in active beige adipocytes with *FTO* obesity-risk alleles (Fig. 2G-H). One-carbon unit metabolism forms a functional interaction with mitochondrial oxidative phosphorylation that is crucial for ATP or heat generation in mammalian cells [Lucas *et al*., 2018]. Lower serine influx that can result in the decrease of one-carbon unit levels may lead to lower amounts of mitochondrial complex subunits I, II and, IV in active beige adipocytes carrying *FTO* obesity-risk genotype (Fig. 4A-F). As a consequence, stimulated maximal and proton leak respiration, which positively correlates with UCP1 activity, and extracellular acidification were suppressed in active beige adipocytes originated from *FTO* obesity-risk genotype carriers. Active beige adipocytes with risk-free genotype also consumed higher amounts of glutamine as compared to white or inactive beige adipocytes carrying the same alleles. The *FTO* obesity-risk carrier adipocytes did not consume glutamine irrespective of the differentiation protocol used (Fig. 6E). Glutamine is the most abundant free amino acid in the circulation [Hall and McCauley, 1996] and in the applied DMEM-F12 cell culture medium. It is one of the main fuel resources for cells supplying carbon atoms to drive the TCA cycle and generate ATP (or heat) [Scalise *et al*., 2016]. Lower expression of *GPT2* and glutamine consumption may contribute to the downregulation of pyruvate metabolism and TCA cycle pathway in active beige adipocytes with obesity-risk alleles (Fig. 2G). Altogether, our findings suggest that adipocytes derived from abdominal SC tissues of *FTO* obesity-risk carriers exert lower uptake of several amino acids as substrates of cellular metabolic processes contributing to compromised energy dissipation.

The positive correlation between *FTO* rs1421085 SNP and obesity or increased body mass index has been reported in several populations such as Estonian children [Katus *et al*., 2020], Chinese children [Wang *et al*., 2013], Iranian adults [Hasan-Bonab *et al*., 2022], Arabic [Hebbar *et al*., 2020], Pakistani [Rana and Bhatti, 2020], Balinese [Priliani *et al*., 2020], and Mexican Mayan girls [Gonzales-Herrera *et al*., 2019]. Through genome-wide association meta-analyses of more than 100000 individuals of European ancestry without diabetes, *FTO* rs1421085 SNP was found to be significantly associated with fasting insulin levels [Scott *et al*., 2012]. A recently published study by Laber *et al*. (2021) showed that an engineered deletion of the rs1421085 conserved cis-regulatory module in mice prevented high fat diet-induced obesity, decreased whole-body fat mass, and elevated the number of mitochondria in WAT. Our presented data highlight the critical effect of *FTO* rs1421085 SNP on human abdominal SC adipocytes only when they are activated for thermogenesis. Leitner *et al*., (2017) reported that large amounts of brownable adipocytes can be found in abdominal SC fat. However, the activation of these adipocytes in humans to reduce adiposity remains challenging. Although these cells can be potentially activated, our previous [Tóth *et al*., 2020] and present results have pointed to a strong effect of obesity-risk genotype at *FTO* rs1421085 SNP, which has a high prevalence in the European population [Dina *et al*., 2007; Babenko *et al*., 2019; Hudek *et al*., 2018] and Mexican children (12.93% to 18.67%) [González-Herrera *et al*., 2018] that must be overcome to enable efficient thermogenesis and weight loss. Our findings further support the importance of genetic background not only in the pathogenesis of obesity but also in the potential effectivity of novel therapeutic approaches which target thermogenesis-related energy dissipation.

## 6 Author Contributions

AV, RAr, BÁV, RAl, AS performed the experiments. EK, RAr, and AV conceptualized the research, with inputs from LF. SP performed the RNA-seq. AG and ÉC carried out measurements of amino acid concentration. IC and GM acquired and analyzed confocal microscopy images. CL and ZB provided adipose tissue samples. EK and LF supervised the research and acquired funding. RAr and AV wrote the original draft of the manuscript. EK and LF edited the final version of the manuscript.

## 8 Acknowledgments

We thank Dr. Zsuzsa Szondy for her exceptional help in reviewing the manuscript before its submission and Jennifer Nagy for technical assistance.

## 10 Data Availability Statement

The RNA-seq datasets generated and analyzed for this study can be found in the Sequence Read Archive (SRA) database [https://www.ncbi.nlm.nih.gov/sra] (access date 26.01.2023.) under accession number PRJNA928240.

