## Supplementary material for "Human Abdominal Subcutaneous-Derived Active Beige Adipocytes Carrying *FTO* rs1421085 Obesity-Risk Alleles Exert Lower Thermogenic Capacity"

### 1 Supplementary Figures and Tables

#### 1.1 Supplementary Figures

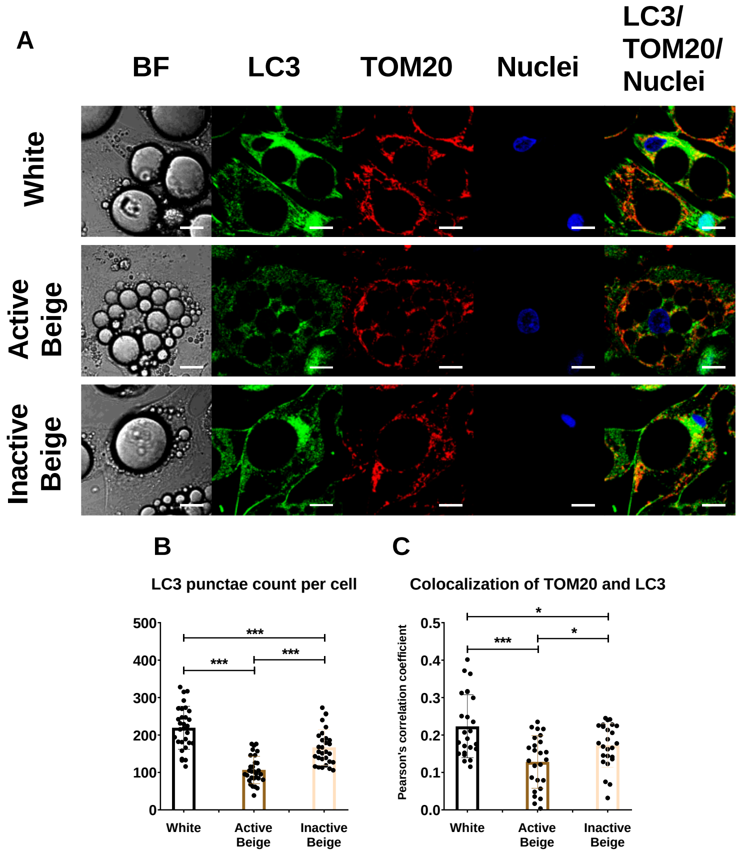


**Supplementary Figure 1.** Confocal microscopy images show decreased mitophagy rate in active beige abdominal subcutaneous derived adipocytes. (A) Representative confocal microscopy images of microtubule-associated protein 1 light chain 3 (LC3) and Translocase of outer mitochondrial membrane (TOM) 20 immunostaining of white, active beige, and inactive beige adipocytes. (B) Quantification of LC3 punctae, n=30. (C) Co-localization of TOM20 and LC3, n=25 per cells. Statistical analysis by ANOVA, *p<0.05 and ***p<0.001. BF: bright field, scalebars represent 10 μm.

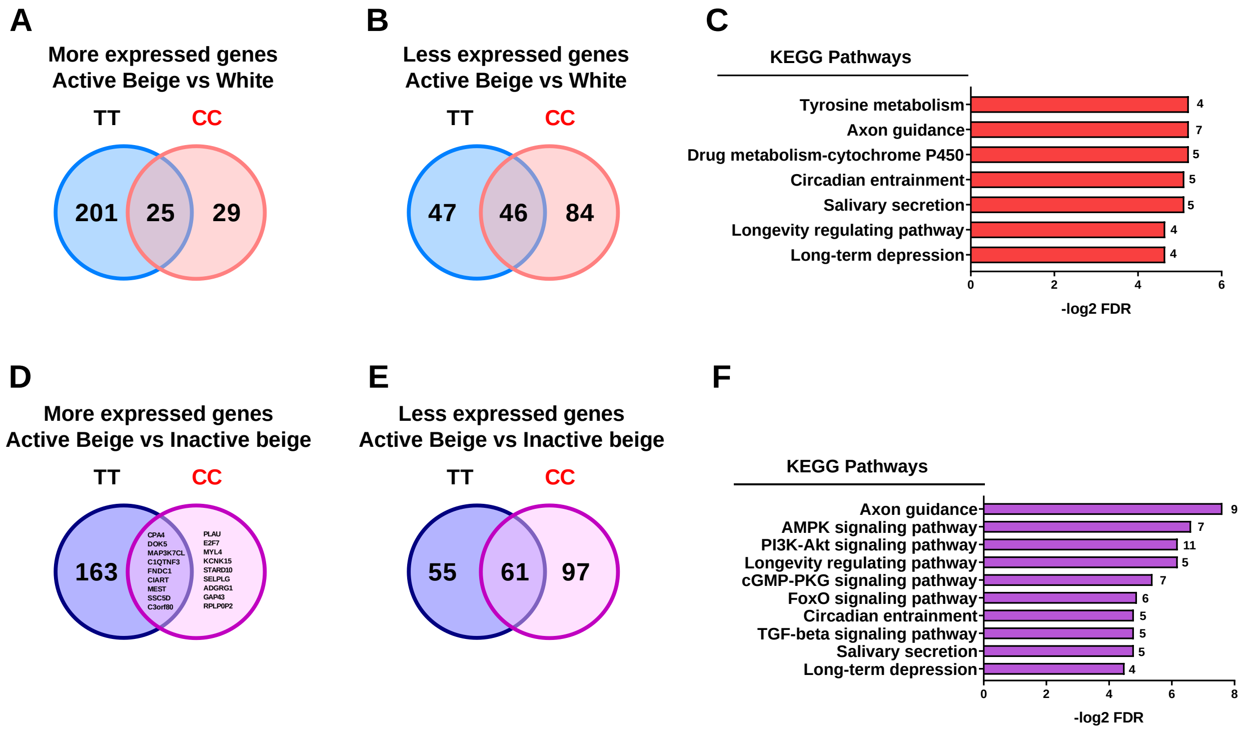


**Supplementary Figure 2.** The effect of *FTO* rs1421085 on the gene expression pattern of active beige adipocytes as compared to white or inactive beige adipocytes. (A-B) Higher (A) and lower (B) expressed genes in active beige as compared to white adipocytes carrying *FTO* TT or CC. (C) Overrepresented pathways which are less expressed in active beige as compared to white differentiated adipocytes carrying *FTO* CC/obesity-risk variant. Numbers on the right side indicate the number of genes involved in the pathways. (D-E) Higher (D) and lower (E) expressed genes in active beige as compared to inactive beige adipocytes carrying *FTO* TT or CC. (F) Overrepresented pathways which are less expressed in active beige as compared to inactive beige differentiated adipocytes carrying *FTO* CC/obesity-risk variant. Numbers on the right side indicate the number of genes involved in the pathways.

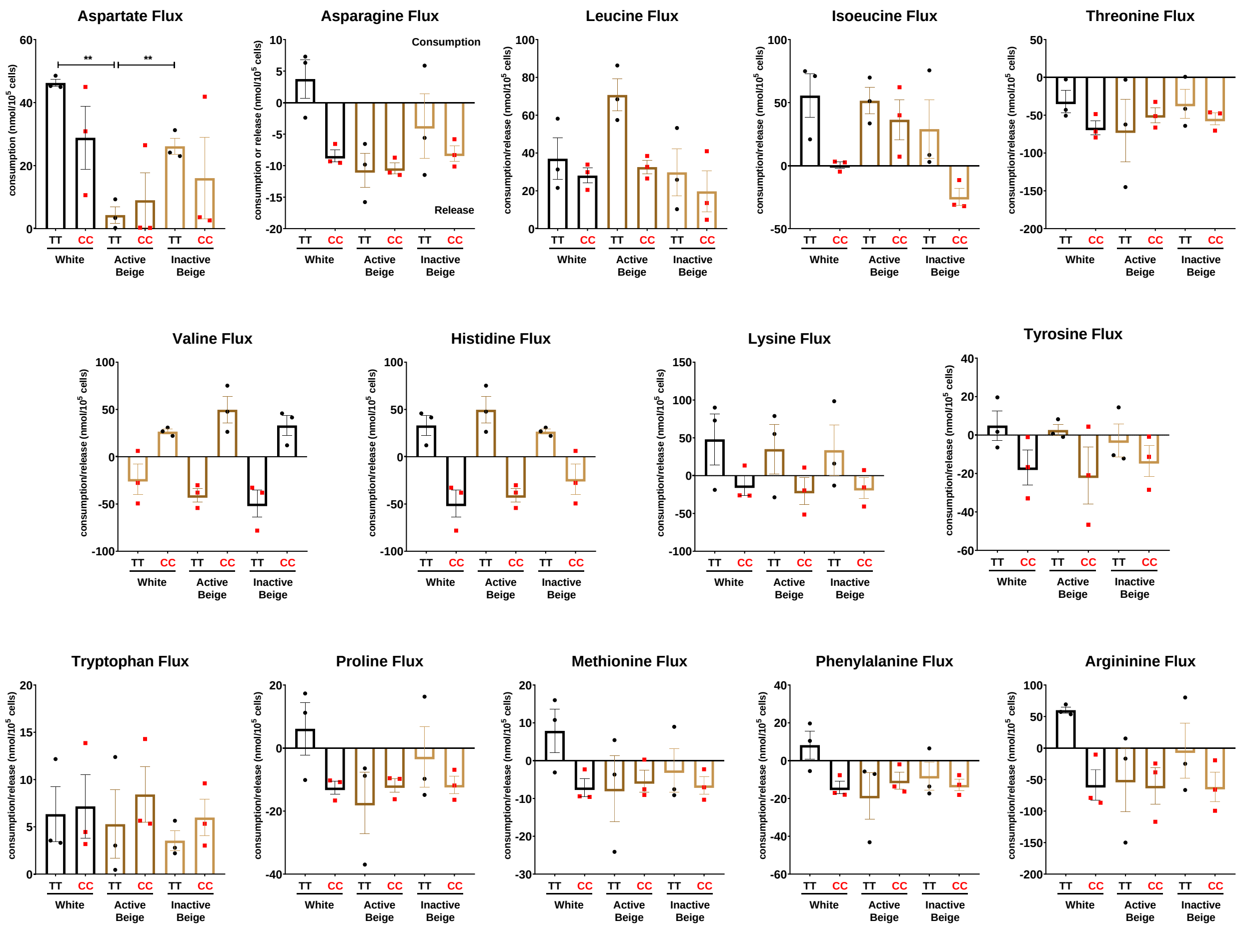


**Supplementary Figure 3.** The effect of differentiation protocols and alleles at *FTO* rs1421085 on the amino acid fluxes of differentiated abdominal subcutaneous adipocytes. Amino acid fluxes were measured in the conditioned media of differentiated abdominal subcutaneous adipocytes, n=3 of each genotype. Statistical analysis was performed by ANOVA, $^{**}p<0.01$.

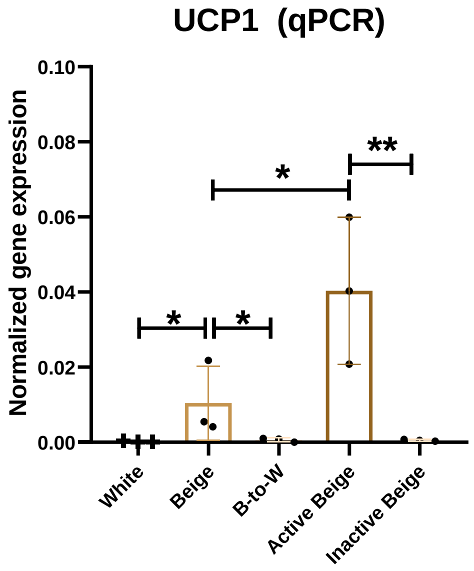


**Supplementary Figure 4.** The mRNA expression of *UCP1* in abdominal subcutaneous adipocytes differentiated under white, beige, beige to white transition (B-to-W), active beige, or inactive beige protocols for 28 days, analyzed by RT-qPCR. In the case of B-to-W and inactive beige protocols, the beige differentiation cocktail was replaced to white at day 14. In the case of active beige and inactive beige protocols, adipocytes received a single bolus treatment of dibutyryl-cAMP at 500 μM concentration for 4 hours at day 14. Statistical analysis was performed by ANOVA, n=3, *p<0.05 and **p<0.01.

### 1.2 Supplementary Tables

Supplementary Table 1. Gene primers and probes

| GENES | ASSAY ID |
|---|---|
| *UCP1* | Hs00222453_m1 |
| *UCP2* | Hs01075227_m1 |
| *PM20D1* | Hs00399438_m1 |
| *CIDEA* | Hs00154455_m1 |
| *CITED1* | Hs00918445_g1 |
| *CKMT1A/B* | Hs00179727_m1 |
| *CKMT2* | Hs00176502_m1 |
| *CPT2* | Hs00988962_m1 |
| *PLIN1* | Hs00160173_m1 |
| *S100b* | Hs00902901_m1 |
| *SLC7A10* | Hs00219813_m1 |
| *SHMT1* | Hs00541043_g1 |
| *GPT2* | Hs00370287_m1 |
| *GAPDH* | Hs99999905_m1 |

Supplementary Table 2. Differentially expressed genes (DEGs) from comparison of active beige and white adipocytes; n=8

| Symbol | Gene Name | Log2 Fold Change | Adj. P value |
|---|---|---|---|
| *KRT79* | keratin 79 | 2.43 | 2.37E-11 |
| *CD96* | CD96 molecule | 2.40 | 5.82E-11 |
| *PCK1* | phosphoenolpyruvate carboxykinase 1 | 2.40 | 1.55E-12 |
| *PM20D1* | peptidase M20 domain containing 1 | 2.22 | 1.38E-08 |
| *CPA4* | carboxypeptidase A4 | 2.20 | 3.95E-10 |
| *KCNA1* | potassium voltage-gated channel subfamily A member 1 | 2.10 | 2.08E-10 |
| *SCN4A* | sodium voltage-gated channel alpha subunit 4 | 2.04 | 2.49E-08 |
| *LAMP3* | lysosomal associated membrane protein 3 | 1.94 | 1.27E-06 |
| *FNDC1* | fibronectin type III domain containing 1 | 1.94 | 2.94E-08 |
| *FCN1* | ficolin 1 | 1.91 | 2.24E-06 |
| *AQP3* | aquaporin 3 (Gill blood group) | 1.81 | 2.71E-07 |
| *C1QTNF3* | C1q and TNF related 3 | 1.78 | 2.11E-06 |
| *THBD* | thrombomodulin | 1.77 | 5.37E-08 |
| *SLC7A10* | solute carrier family 7 member 10 | 1.77 | 1.37E-05 |
| *SLC22A12* | solute carrier family 22 member 12 | 1.73 | 3.58E-05 |
| *TENT5C* | terminal nucleotidyltransferase 5C | 1.72 | 1.08E-07 |
| *SLC24A2* | solute carrier family 24 member 2 | 1.69 | 6.53E-05 |
| *CD52* | CD52 molecule | 1.66 | 4.82E-05 |
| *GAP43* | growth associated protein 43 | 1.64 | 0.000107485 |
| *CTXN1* | cortexin 1 | 1.64 | 2.26E-08 |
| *BBOX1* | gamma-butyrobetaine hydroxylase 1 | 1.64 | 7.62E-05 |
| *LINC02554* | long intergenic non-protein coding RNA 2554 | 1.62 | 0.000156997 |
| *TSHR* | thyroid stimulating hormone receptor | 1.59 | 0.000215184 |
| *KCNK12* | potassium two pore domain channel subfamily K member 12 | 1.58 | 0.000242958 |
| *MEST* | mesoderm specific transcript | 1.55 | 1.45E-10 |
| *ZNF365* | zinc finger protein 365 | 1.54 | 2.13E-07 |
| *CPA2* | carboxypeptidase A2 | 1.54 | 0.000314379 |
| *MAP3K7CL* | MAP3K7 C-terminal like | 1.53 | 3.53E-10 |
| *IL1B* | interleukin 1 beta | 1.52 | 0.000156997 |
| *EPB41L4B* | erythrocyte membrane protein band 4.1 like 4B | 1.52 | 5.83E-08 |

| | | | |
|---|---|---|---|
| *RTKN2* | rhotekin 2 | 1.50 | 6.91E-08 |
| *TMEM200C* | transmembrane protein 200C | 1.50 | 8.95E-05 |
| *TMEM130* | transmembrane protein 130 | 1.48 | 0.000751043 |
| *PDZD2* | PDZ domain containing 2 | 1.47 | 1.72E-05 |
| *LOC100507560* | uncharacterized LOC100507560 | 1.46 | 0.000189853 |
| *PPP1R14C* | protein phosphatase 1 regulatory inhibitor subunit 14C | 1.45 | 0.000889281 |
| *KCNK15* | potassium two pore domain channel subfamily K member 15 | 1.45 | 1.38E-05 |
| *ECSCR* | endothelial cell surface expressed chemotaxis and apoptosis regulator | 1.44 | 0.000937479 |
| *CEMIP* | cell migration inducing hyaluronidase 1 | 1.43 | 1.94E-05 |
| *LINGO1* | leucine rich repeat and Ig domain containing 1 | 1.41 | 0.000178003 |
| *KCNMA1* | potassium calcium-activated channel subfamily M alpha 1 | 1.41 | 0.000622242 |
| *RBM24* | RNA binding motif protein 24 | 1.41 | 5.75E-08 |
| *STX11* | syntaxin 11 | 1.41 | 5.37E-08 |
| *CDH6* | cadherin 6 | 1.39 | 0.000402281 |
| *CLDN2* | claudin 2 | 1.39 | 0.001821264 |
| *SORL1* | sortilin related receptor 1 | 1.38 | 0.001236163 |
| *ADAMTS14* | ADAM metallopeptidase with thrombospondin type 1 motif 14 | 1.37 | 0.000177898 |
| *LOC100506253* | uncharacterized LOC100506253 | 1.37 | 0.002219581 |
| *CIART* | circadian associated repressor of transcription | 1.36 | 7.13E-15 |
| *TAGAP* | T cell activation RhoGTPase activating protein | 1.35 | 0.002422638 |
| *CA3* | carbonic anhydrase 3 | 1.34 | 0.003390067 |
| *KIRREL3* | kirre like nephrin family adhesion molecule 3 | 1.34 | 0.001332712 |
| *GK* | glycerol kinase | 1.32 | 3.32E-05 |
| *CYS1* | cystin 1 | 1.31 | 5.88E-07 |
| *RGL3* | ral guanine nucleotide dissociation stimulator like 3 | 1.30 | 0.000607669 |
| *CASZ1* | castor zinc finger 1 | 1.30 | 0.001332712 |
| *PRAG1* | PEAK1 related, kinase-activating pseudokinase 1 | 1.30 | 0.003661237 |
| *ABCG1* | ATP binding cassette subfamily G member 1 | 1.30 | 0.000100674 |
| *RFPL4B* | ret finger protein like 4B | 1.30 | 0.003092843 |
| *ICA1* | islet cell autoantigen 1 | 1.29 | 0.000281869 |

| | | | |
|---|---|---|---|
| *LOC100507006* | uncharacterized LOC100507006 | 1.29 | 0.001493056 |
| *NA* | NA | 1.29 | 0.005606153 |
| *KLHDC7B* | kelch domain containing 7B | 1.28 | 0.006574118 |
| *NA* | NA | 1.26 | 0.006876331 |
| *LMO2* | LIM domain only 2 | 1.25 | 0.000269485 |
| *ADAMTS18* | ADAM metallopeptidase with thrombospondin type 1 motif 18 | 1.23 | 0.0104071 |
| *EPHB2* | EPH receptor B2 | 1.23 | 1.97E-05 |
| *SCAMP5* | secretory carrier membrane protein 5 | 1.22 | 0.001166631 |
| *TMEM63C* | transmembrane protein 63C | 1.22 | 0.011482877 |
| *SLC22A3* | solute carrier family 22 member 3 | 1.21 | 0.000821422 |
| *ITGA8* | integrin subunit alpha 8 | 1.20 | 0.005161367 |
| *MYOM3* | myomesin 3 | 1.18 | 0.014018549 |
| *ADAM12* | ADAM metallopeptidase domain 12 | 1.17 | 0.001221323 |
| *MYL4* | myosin light chain 4 | 1.17 | 0.01316891 |
| *NREP* | neuronal regeneration related protein | 1.16 | 0.005021214 |
| *HOXD1* | homeobox D1 | 1.16 | 0.008846877 |
| *IQCA1* | IQ motif containing with AAA domain 1 | 1.15 | 0.00189403 |
| *STAC* | SH3 and cysteine rich domain | 1.15 | 0.001409092 |
| *CITED1* | Cbp/p300 interacting transactivator with Glu/Asp rich carboxy-terminal domain 1 | 1.14 | 0.010822663 |
| *KRT80* | keratin 80 | 1.14 | 0.013421498 |
| *INA* | internexin neuronal intermediate filament protein alpha | 1.14 | 0.009387775 |
| *CIDEA* | cell death inducing DFFA like effector a | 1.14 | 0.022804912 |
| *CYP26A1* | cytochrome P450 family 26 subfamily A member 1 | 1.13 | 0.023417525 |
| *CSDC2* | cold shock domain containing C2 | 1.13 | 0.000730888 |
| *PPP1R1A* | protein phosphatase 1 regulatory inhibitor subunit 1A | 1.13 | 0.019974952 |
| *AP3B2* | adaptor related protein complex 3 subunit beta 2 | 1.12 | 0.011744388 |
| *FGF13* | fibroblast growth factor 13 | 1.12 | 0.010973719 |
| *E2F7* | E2F transcription factor 7 | 1.12 | 0.001400194 |
| *CXADRP3* | CXADR pseudogene 3 | 1.12 | 0.020497723 |
| *EPHA2* | EPH receptor A2 | 1.12 | 0.004129368 |

| | | | |
|---|---|---|---|
| *RAB39B* | RAB39B, member RAS oncogene family | 1.12 | 0.020181492 |
| *NDUFA4L2* | NDUFA4 mitochondrial complex associated like 2 | 1.10 | 0.001531789 |
| *NA* | NA | 1.10 | 0.028586024 |
| *HAS3* | hyaluronan synthase 3 | 1.10 | 0.002527803 |
| *RSPO3* | R-spondin 3 | 1.09 | 0.028518081 |
| *RTN4R* | reticulon 4 receptor | 1.09 | 0.008569867 |
| *DLX3* | distal-less homeobox 3 | 1.09 | 0.022055084 |
| *PLIN5* | perilipin 5 | 1.09 | 0.017677372 |
| *SELPLG* | selectin P ligand | 1.09 | 0.000121056 |
| *APLP1* | amyloid beta precursor like protein 1 | 1.09 | 0.002758266 |
| *PCDH19* | protocadherin 19 | 1.08 | 0.008475504 |
| *RGS7BP* | regulator of G protein signaling 7 binding protein | 1.08 | 0.020727321 |
| *HRH2* | histamine receptor H2 | 1.08 | 0.035283159 |
| *CKMT1B* | creatine kinase, mitochondrial 1B | 1.08 | 0.021991002 |
| *CX3CR1* | C-X3-C motif chemokine receptor 1 | 1.08 | 0.036754041 |
| *OASL* | 2'-5'-oligoadenylate synthetase like | 1.07 | 0.029556704 |
| *ANOS1* | anosmin 1 | 1.07 | 0.028285618 |
| *STK33* | serine/threonine kinase 33 | 1.07 | 0.020727321 |
| *NRG1* | neuregulin 1 | 1.07 | 0.034815048 |
| *CH25H* | cholesterol 25-hydroxylase | 1.07 | 0.018722712 |
| *PALMD* | palmdelphin | 1.07 | 0.018125851 |
| *HCAR3* | hydroxycarboxylic acid receptor 3 | 1.06 | 0.026478959 |
| *SAMD10* | sterile alpha motif domain containing 10 | 1.06 | 1.37E-05 |
| *SLITRK6* | SLIT and NTRK like family member 6 | 1.05 | 0.025144925 |
| *RTN1* | reticulon 1 | 1.05 | 0.026215254 |
| *SLCO2A1* | solute carrier organic anion transporter family member 2A1 | 1.05 | 0.016809637 |
| *PDE1B* | phosphodiesterase 1B | 1.05 | 0.037056444 |
| *TGM1* | transglutaminase 1 | 1.05 | 0.026513345 |
| *HERC5* | HECT and RLD domain containing E3 ubiquitin protein ligase 5 | 1.05 | 0.04518502 |
| *SAMD12* | sterile alpha motif domain containing 12 | 1.04 | 0.032591111 |
| *PLPPR4* | phospholipid phosphatase related 4 | 1.04 | 0.033802231 |
| *EXTL1* | exostosin like glycosyltransferase 1 | 1.04 | 0.047854579 |
| *NA* | NA | 1.04 | 0.037056444 |

| | | | |
|---|---|---|---|
| *DAO* | D-amino acid oxidase | 1.04 | 0.049634866 |
| *REN* | renin | 1.03 | 0.042511987 |
| *ADAMTS16* | ADAM metallopeptidase with thrombospondin type 1 motif 16 | 1.03 | 0.028518081 |
| *ABCA3* | ATP binding cassette subfamily A member 3 | 1.03 | 0.005227157 |
| *RHPN1* | rhophilin Rho GTPase binding protein 1 | 1.03 | 0.018125851 |
| *FUCA1* | alpha-L-fucosidase 1 | 1.02 | 1.32E-05 |
| *ADGRG1* | adhesion G protein-coupled receptor G1 | 1.02 | 0.003171108 |
| *MGAT3* | beta-1,4-mannosyl-glycoprotein 4-beta-N-acetylglucosaminyltransferase | 1.01 | 0.019622647 |
| *TGFB3* | transforming growth factor beta 3 | 0.99 | 0.006348203 |
| *IGFBPL1* | insulin like growth factor binding protein like 1 | 0.98 | 0.010969325 |
| *DOK5* | docking protein 5 | 0.98 | 4.47E-11 |
| *INPP5J* | inositol polyphosphate-5-phosphatase J | 0.97 | 0.044715424 |
| *IL20RA* | interleukin 20 receptor subunit alpha | 0.96 | 0.000342765 |
| *LAMC3* | laminin subunit gamma 3 | 0.96 | 0.049941894 |
| *ASPHD2* | aspartate beta-hydroxylase domain containing 2 | 0.95 | 0.0005617 |
| *RUBCNL* | rubicon like autophagy enhancer | 0.95 | 0.030413861 |
| *GPC1* | glypican 1 | 0.95 | 4.18E-06 |
| *TTC9* | tetratricopeptide repeat domain 9 | 0.95 | 0.041059956 |
| *DACT3* | dishevelled binding antagonist of beta catenin 3 | 0.94 | 0.0005617 |
| *DUSP8* | dual specificity phosphatase 8 | 0.94 | 0.000874179 |
| *LRRC73* | leucine rich repeat containing 73 | 0.94 | 0.008356352 |
| *SAMD14* | sterile alpha motif domain containing 14 | 0.94 | 0.010151475 |
| *ATP1A3* | ATPase Na+/K+ transporting subunit alpha 3 | 0.94 | 0.047614233 |
| *CCDC148* | coiled-coil domain containing 148 | 0.93 | 0.027762374 |
| *PPM1H* | protein phosphatase, Mg2+/Mn2+ dependent 1H | 0.93 | 0.00493619 |
| *PLEKHA7* | pleckstrin homology domain containing A7 | 0.93 | 0.0253494 |
| *PDE1C* | phosphodiesterase 1C | 0.93 | 0.035283159 |
| *RPS6KA1* | ribosomal protein S6 kinase A1 | 0.93 | 0.038776363 |

| | | | |
|---|---|---|---|
| *PHLDA1* | pleckstrin homology like domain family A member 1 | 0.93 | 0.015573203 |
| *ITIH5* | inter-alpha-trypsin inhibitor heavy chain 5 | 0.92 | 0.009681211 |
| *EVA1A* | eva-1 homolog A, regulator of programmed cell death | 0.92 | 0.012983106 |
| *RCAN1* | regulator of calcineurin 1 | 0.92 | 0.002048462 |
| *PCSK1* | proprotein convertase subtilisin/kexin type 1 | 0.92 | 0.026822353 |
| *TGM5* | transglutaminase 5 | 0.92 | 0.003476304 |
| *LINC00856* | long intergenic non-protein coding RNA 856 | 0.92 | 0.010151475 |
| *NA* | NA | 0.92 | 0.026215254 |
| *NXPH4* | neurexophilin 4 | 0.91 | 0.045314329 |
| *CNKSR2* | connector enhancer of kinase suppressor of Ras 2 | 0.90 | 0.041059956 |
| *SMYD2* | SET and MYND domain containing 2 | 0.89 | 0.018791317 |
| *TMEM131L* | transmembrane 131 like | 0.88 | 0.00245372 |
| *EGFLAM* | EGF like, fibronectin type III and laminin G domains | 0.88 | 0.03073498 |
| *HOMER2* | homer scaffold protein 2 | 0.88 | 0.012940255 |
| *STAT4* | signal transducer and activator of transcription 4 | 0.88 | 0.00569635 |
| *ZP3* | zona pellucida glycoprotein 3 | 0.87 | 0.013398037 |
| *NA* | NA | 0.87 | 0.001168463 |
| *SORCS2* | sortilin related VPS10 domain containing receptor 2 | 0.86 | 0.028518081 |
| *NR3C2* | nuclear receptor subfamily 3 group C member 2 | 0.85 | 0.0104071 |
| *MIR1915HG* | MIR1915 host gene | 0.85 | 0.007580509 |
| *LACC1* | laccase domain containing 1 | 0.85 | 0.002033882 |
| *SEL1L3* | SEL1L family member 3 | 0.85 | 0.011317739 |
| *NCS1* | neuronal calcium sensor 1 | 0.84 | 2.08E-07 |
| *ASS1* | argininosuccinate synthase 1 | 0.84 | 0.000202089 |
| *LOXL1-AS1* | LOXL1 antisense RNA 1 | 0.84 | 0.004689278 |
| *GXYLT2* | glucoside xylosyltransferase 2 | 0.84 | 0.002061228 |
| *L3MBTL2-AS1* | L3MBTL2 antisense RNA 1 | 0.83 | 0.017538044 |
| *WARS1* | tryptophanyl-tRNA synthetase 1 | 0.82 | 0.011745386 |
| *TUFT1* | tuftelin 1 | 0.82 | 0.002749693 |
| *MTHFD1L* | methylenetetrahydrofolate dehydrogenase (NADP+ dependent) 1 like | 0.80 | 0.026513345 |

| | | | |
|---|---|---|---|
| *CHN2* | chimerin 2 | 0.78 | 0.027106408 |
| *KIAA1549L* | KIAA1549 like | 0.78 | 0.000936344 |
| *MELTF-AS1* | MELTF antisense RNA 1 | 0.77 | 0.01576322 |
| *DAB2* | DAB adaptor protein 2 | 0.77 | 0.010969325 |
| *ATP8B1* | ATPase phospholipid transporting 8B1 | 0.76 | 0.004560314 |
| *ARNTL2* | aryl hydrocarbon receptor nuclear translocator like 2 | 0.76 | 0.017567259 |
| *E2F5* | E2F transcription factor 5 | 0.75 | 0.005678885 |
| *RNF122* | ring finger protein 122 | 0.75 | 0.001358663 |
| *GLIS2* | GLIS family zinc finger 2 | 0.75 | 7.79E-05 |
| *MOCOS* | molybdenum cofactor sulfurase | 0.75 | 0.001215003 |
| *ARID5A* | AT-rich interaction domain 5A | 0.75 | 0.027106408 |
| *GPR153* | G protein-coupled receptor 153 | 0.74 | 0.001215003 |
| *FNIP2* | folliculin interacting protein 2 | 0.74 | 0.023101885 |
| *MYOM1* | myomesin 1 | 0.74 | 0.020063543 |
| *RGS3* | regulator of G protein signaling 3 | 0.74 | 1.66E-05 |
| *ZNF280B* | zinc finger protein 280B | 0.73 | 0.02553804 |
| *SHB* | SH2 domain containing adaptor protein B | 0.73 | 0.001343943 |
| *GUCY1A2* | guanylate cyclase 1 soluble subunit alpha 2 | 0.73 | 0.036754041 |
| *PARD6A* | par-6 family cell polarity regulator alpha | 0.72 | 0.047614233 |
| *GTPBP2* | GTP binding protein 2 | 0.72 | 0.001037711 |
| *AMACR* | alpha-methylacyl-CoA racemase | 0.72 | 1.64E-05 |
| *SDSL* | serine dehydratase like | 0.71 | 0.00316593 |
| *GSDME* | gasdermin E | 0.71 | 0.011482877 |
| *SERP2* | stress associated endoplasmic reticulum protein family member 2 | 0.71 | 0.00636045 |
| *TPST2* | tyrosylprotein sulfotransferase 2 | 0.71 | 0.000806423 |
| *BCAS4* | breast carcinoma amplified sequence 4 | 0.71 | 0.043472176 |
| *RALGPS1* | Ral GEF with PH domain and SH3 binding motif 1 | 0.70 | 0.023156482 |
| *MYO1B* | myosin IB | 0.70 | 0.004591736 |
| *ANKRD34A* | ankyrin repeat domain 34A | 0.70 | 0.005817268 |
| *ZBTB16* | zinc finger and BTB domain containing 16 | -4.81 | 1.34E-117 |
| *HIF3A* | hypoxia inducible factor 3 subunit alpha | -3.01 | 3.30E-17 |
| *FKBP5* | FKBP prolyl isomerase 5 | -2.83 | 2.05E-25 |
| *TIMP4* | TIMP metallopeptidase inhibitor 4 | -2.52 | 2.61E-11 |

| | | | |
|---|---|---|---|
| *PER1* | period circadian regulator 1 | -2.37 | 3.39E-45 |
| *SLC16A12* | solute carrier family 16 member 12 | -2.33 | 3.54E-10 |
| *ABCC2* | ATP binding cassette subfamily C member 2 | -2.29 | 6.34E-10 |
| *LRP1B* | LDL receptor related protein 1B | -2.24 | 1.05E-08 |
| *ANGPTL8* | angiopoietin like 8 | -2.22 | 1.38E-08 |
| *SAA1* | serum amyloid A1 | -2.20 | 1.89E-08 |
| *PILRA* | paired immunoglobin like type 2 receptor alpha | -2.15 | 3.53E-10 |
| *MAOA* | monoamine oxidase A | -2.11 | 1.23E-10 |
| *C6* | complement C6 | -2.11 | 3.43E-08 |
| *INHBB* | inhibin subunit beta B | -2.05 | 3.71E-11 |
| *CYP4B1* | cytochrome P450 family 4 subfamily B member 1 | -1.97 | 7.02E-07 |
| *GALNT15* | polypeptide N-acetylgalactosaminyltransferase 15 | -1.89 | 1.88E-06 |
| *TMPRSS5* | transmembrane serine protease 5 | -1.88 | 3.19E-06 |
| *RAPGEF5* | Rap guanine nucleotide exchange factor 5 | -1.86 | 5.02E-06 |
| *TMC2* | transmembrane channel like 2 | -1.83 | 2.75E-06 |
| *NEGR1* | neuronal growth regulator 1 | -1.82 | 9.04E-22 |
| *MMP28* | matrix metallopeptidase 28 | -1.81 | 1.17E-06 |
| *FMO2* | flavin containing dimethylaniline monoxygenase 2 | -1.81 | 1.37E-06 |
| *CRISPLD2* | cysteine rich secretory protein LCCL domain containing 2 | -1.79 | 1.23E-10 |
| *LMO3* | LIM domain only 3 | -1.78 | 2.45E-08 |
| *GRIA1* | glutamate ionotropic receptor AMPA type subunit 1 | -1.75 | 7.60E-07 |
| *STC1* | stanniocalcin 1 | -1.74 | 1.29E-05 |
| *AVPR1A* | arginine vasopressin receptor 1A | -1.72 | 3.69E-05 |
| *CPM* | carboxypeptidase M | -1.72 | 1.50E-05 |
| *NA* | NA | -1.71 | 3.25E-05 |
| *RASL11A* | RAS like family 11 member A | -1.71 | 1.97E-08 |
| *ATP1A2* | ATPase Na+/K+ transporting subunit alpha 2 | -1.71 | 3.72E-05 |
| *PKD2L1* | polycystin 2 like 1, transient receptor potential cation channel | -1.70 | 5.25E-05 |
| *GPX3* | glutathione peroxidase 3 | -1.68 | 1.29E-05 |
| *KIAA0040* | KIAA0040 | -1.68 | 6.53E-05 |
| *IP6K3* | inositol hexakisphosphate kinase 3 | -1.66 | 8.72E-05 |
| *METTL7A* | methyltransferase like 7A | -1.66 | 9.37E-06 |

| | | | |
|---|---|---|---|
| *GLYAT* | glycine-N-acyltransferase | -1.66 | 8.95E-05 |
| *F5* | coagulation factor V | -1.65 | 9.41E-05 |
| *SYN2* | synapsin II | -1.64 | 0.000114669 |
| *ALS2CL* | ALS2 C-terminal like | -1.63 | 0.000100674 |
| *GGT5* | gamma-glutamyltransferase 5 | -1.61 | 5.30E-05 |
| *PRODH* | proline dehydrogenase 1 | -1.60 | 9.41E-05 |
| *ACSL6* | acyl-CoA synthetase long chain family member 6 | -1.58 | 0.00023662 |
| *PLCH1* | phospholipase C eta 1 | -1.57 | 0.000270556 |
| *GLUL* | glutamate-ammonia ligase | -1.56 | 2.74E-07 |
| *ARHGEF16* | Rho guanine nucleotide exchange factor 16 | -1.56 | 0.000301928 |
| *APCDD1* | APC down-regulated 1 | -1.54 | 5.25E-05 |
| *DPT* | dermatopontin | -1.54 | 4.52E-05 |
| *CLCN1* | chloride voltage-gated channel 1 | -1.53 | 0.000292718 |
| *SERPINA3* | serpin family A member 3 | -1.53 | 4.91E-05 |
| *MAP2K6* | mitogen-activated protein kinase kinase 6 | -1.52 | 0.000352204 |
| *RXFP1* | relaxin family peptide receptor 1 | -1.51 | 0.000229271 |
| *APOD* | apolipoprotein D | -1.49 | 0.000656841 |
| *FAM166B* | family with sequence similarity 166 member B | -1.49 | 0.000584654 |
| *ACKR2* | atypical chemokine receptor 2 | -1.48 | 5.91E-06 |
| *KCNK5* | potassium two pore domain channel subfamily K member 5 | -1.48 | 0.0006409 |
| *ANGPTL1* | angiopoietin like 1 | -1.47 | 3.46E-06 |
| *FGD4* | FYVE, RhoGEF and PH domain containing 4 | -1.47 | 4.52E-05 |
| *HS3ST2* | heparan sulfate-glucosamine 3-sulfotransferase 2 | -1.47 | 0.000806423 |
| *MGP* | matrix Gla protein | -1.46 | 0.000202089 |
| *TSC22D3* | TSC22 domain family member 3 | -1.45 | 6.17E-22 |
| *ADRA1B* | adrenoceptor alpha 1B | -1.45 | 5.88E-07 |
| *KDR* | kinase insert domain receptor | -1.45 | 0.001125985 |
| *LINC01088* | long intergenic non-protein coding RNA 1088 | -1.44 | 0.000306569 |
| *NA* | NA | -1.43 | 0.001332712 |
| *ALOX5AP* | arachidonate 5-lipoxygenase activating protein | -1.43 | 0.00083161 |
| *NKD2* | NKD inhibitor of WNT signaling pathway 2 | -1.42 | 0.001498731 |
| *DPEP1* | dipeptidase 1 | -1.42 | 0.001567438 |

| | | | |
|---|---|---|---|
| *CYP8B1* | cytochrome P450 family 8 subfamily B member 1 | -1.41 | 0.000215184 |
| *MRO* | maestro | -1.41 | 0.000821422 |
| *GSTA1* | glutathione S-transferase alpha 1 | -1.40 | 0.001800055 |
| *EPHB6* | EPH receptor B6 | -1.39 | 0.000593528 |
| *ZNF385B* | zinc finger protein 385B | -1.38 | 0.002168056 |
| *PTGS1* | prostaglandin-endoperoxide synthase 1 | -1.37 | 1.01E-06 |
| *ADH1A* | alcohol dehydrogenase 1A (class I), alpha polypeptide | -1.37 | 0.001111716 |
| *VIT* | vitrin | -1.37 | 0.00033432 |
| *CRLF1* | cytokine receptor like factor 1 | -1.36 | 8.29E-05 |
| *TENT5B* | terminal nucleotidyltransferase 5B | -1.36 | 4.18E-06 |
| *ACSM2A* | acyl-CoA synthetase medium chain family member 2A | -1.36 | 0.002809072 |
| *LOC285847* | uncharacterized LOC285847 | -1.36 | 0.001858037 |
| *GPM6B* | glycoprotein M6B | -1.36 | 3.25E-05 |
| *PLXNA4* | plexin A4 | -1.35 | 1.02E-06 |
| *LINC00482* | long intergenic non-protein coding RNA 482 | -1.35 | 0.000752396 |
| *GPC3* | glypican 3 | -1.35 | 0.003255676 |
| *GABRA5* | gamma-aminobutyric acid type A receptor subunit alpha5 | -1.35 | 0.003333887 |
| *DAAM2-AS1* | DAAM2 antisense RNA 1 | -1.34 | 0.000236189 |
| *NGFR* | nerve growth factor receptor | -1.32 | 0.002939022 |
| *RGCC* | regulator of cell cycle | -1.32 | 0.003036311 |
| *HPD* | 4-hydroxyphenylpyruvate dioxygenase | -1.31 | 0.002049237 |
| *ACSM2B* | acyl-CoA synthetase medium chain family member 2B | -1.31 | 0.004536672 |
| *MYEOV* | myeloma overexpressed | -1.30 | 0.004458824 |
| *MT1X* | metallothionein 1X | -1.30 | 7.93E-06 |
| *PTPN22* | protein tyrosine phosphatase non-receptor type 22 | -1.29 | 0.004907835 |
| *SLC34A2* | solute carrier family 34 member 2 | -1.28 | 0.006593828 |
| *ISM1* | isthmin 1 | -1.28 | 0.006226472 |
| *C2orf88* | chromosome 2 open reading frame 88 | -1.28 | 0.004347359 |
| *LEP* | leptin | -1.27 | 0.005234252 |
| *TTYH1* | tweety family member 1 | -1.27 | 0.005418178 |
| *SLA* | Src like adaptor | -1.27 | 0.007084364 |
| *CCN4* | cellular communication network factor 4 | -1.26 | 0.00560254 |

| | | | |
|---|---|---|---|
| *ADH1B* | alcohol dehydrogenase 1B (class I), beta polypeptide | -1.26 | 0.004045742 |
| *GPR88* | G protein-coupled receptor 88 | -1.26 | 0.006812536 |
| *RIPOR3* | RIPOR family member 3 | -1.25 | 0.007150537 |
| *HSPA6* | heat shock protein family A (Hsp70) member 6 | -1.25 | 0.008124626 |
| *AACS* | acetoacetyl-CoA synthetase | -1.24 | 0.001330104 |
| *ALKAL2* | ALK and LTK ligand 2 | -1.23 | 0.009692994 |
| *CST3* | cystatin C | -1.23 | 1.98E-05 |
| *SLPI* | secretory leukocyte peptidase inhibitor | -1.21 | 0.001879762 |
| *LRRN3* | leucine rich repeat neuronal 3 | -1.21 | 0.008058827 |
| *C2orf72* | chromosome 2 open reading frame 72 | -1.21 | 0.012130134 |
| *HSPA7* | heat shock protein family A (Hsp70) member 7 (pseudogene) | -1.21 | 0.007617462 |
| *OLFM2* | olfactomedin 2 | -1.20 | 0.000806423 |
| *ABLIM3* | actin binding LIM protein family member 3 | -1.20 | 0.001498731 |
| *ALDOC* | aldolase, fructose-bisphosphate C | -1.20 | 0.010233425 |
| *RASSF4* | Ras association domain family member 4 | -1.19 | 0.00020977 |
| *CTHRC1* | collagen triple helix repeat containing 1 | -1.19 | 0.001266085 |
| *POU3F3* | POU class 3 homeobox 3 | -1.18 | 0.011317739 |
| *DNASE1L3* | deoxyribonuclease 1 like 3 | -1.18 | 0.008730918 |
| *TRPV6* | transient receptor potential cation channel subfamily V member 6 | -1.18 | 0.015573203 |
| *SLC4A11* | solute carrier family 4 member 11 | -1.18 | 0.006033697 |
| *PIK3R1* | phosphoinositide-3-kinase regulatory subunit 1 | -1.17 | 0.002344534 |
| *TMEM150C* | transmembrane protein 150C | -1.17 | 0.002738517 |
| *AGTR1* | angiotensin II receptor type 1 | -1.17 | 0.011985927 |
| *PRRT4* | proline rich transmembrane protein 4 | -1.16 | 0.017744288 |
| *DUSP4* | dual specificity phosphatase 4 | -1.16 | 0.006876331 |
| *TRIM16L* | tripartite motif containing 16 like | -1.16 | 0.005161367 |
| *SLC2A13* | solute carrier family 2 member 13 | -1.15 | 0.003690124 |
| *MAPK4* | mitogen-activated protein kinase 4 | -1.15 | 0.020485089 |
| *LINC01554* | long intergenic non-protein coding RNA 1554 | -1.15 | 0.012289729 |
| *TRARG1* | trafficking regulator of GLUT4 (SLC2A4) 1 | -1.14 | 0.020340131 |

| | | | |
|---|---|---|---|
| *KLF9* | Kruppel like factor 9 | -1.14 | 5.37E-10 |
| *RGMA* | repulsive guidance molecule BMP co-receptor a | -1.14 | 1.02E-06 |
| *BOC* | BOC cell adhesion associated, oncogene regulated | -1.13 | 0.000208859 |
| *ZBED3-AS1* | ZBED3 antisense RNA 1 | -1.12 | 0.018125851 |
| *EYA2* | EYA transcriptional coactivator and phosphatase 2 | -1.12 | 0.013696386 |
| *AFAP1L1* | actin filament associated protein 1 like 1 | -1.12 | 0.018334162 |
| *FAM107A* | family with sequence similarity 107 member A | -1.12 | 0.026513345 |
| *TWF2* | twinfilin actin binding protein 2 | -1.11 | 0.003377255 |
| *OMD* | osteomodulin | -1.11 | 0.019915143 |
| *VAV3* | vav guanine nucleotide exchange factor 3 | -1.11 | 0.026184562 |
| *GDF7* | growth differentiation factor 7 | -1.11 | 0.0253494 |
| *TC2N* | tandem C2 domains, nuclear | -1.11 | 0.005418178 |
| *CUTC* | cutC copper transporter | -1.11 | 4.76E-14 |
| *TPRG1* | tumor protein p63 regulated 1 | -1.11 | 0.010821996 |
| *P2RY14* | purinergic receptor P2Y14 | -1.11 | 0.028057773 |
| *ANGPTL5* | angiopoietin like 5 | -1.11 | 0.020881077 |
| *NPTX1* | neuronal pentraxin 1 | -1.11 | 0.028586024 |
| *NA* | NA | -1.10 | 0.009499846 |
| *POM121L9P* | POM121 transmembrane nucleoporin like 9, pseudogene | -1.10 | 0.018281172 |
| *WNT5A* | Wnt family member 5A | -1.09 | 0.005683595 |
| *GNG2* | G protein subunit gamma 2 | -1.08 | 0.023417525 |
| *CLDN7* | claudin 7 | -1.08 | 0.014401277 |
| *CFD* | complement factor D | -1.08 | 0.000359467 |
| *TNFSF10* | TNF superfamily member 10 | -1.07 | 0.020881077 |
| *ERRFI1* | ERBB receptor feedback inhibitor 1 | -1.07 | 3.49E-06 |
| *SMCO3* | single-pass membrane protein with coiled-coil domains 3 | -1.07 | 0.021325691 |
| *NA* | NA | -1.06 | 0.039010781 |
| *KCTD14* | potassium channel tetramerization domain containing 14 | -1.05 | 0.025245361 |
| *P2RY12* | purinergic receptor P2Y12 | -1.05 | 0.041059956 |
| *GDF10* | growth differentiation factor 10 | -1.05 | 0.029701377 |
| *CSRNP3* | cysteine and serine rich nuclear protein 3 | -1.05 | 0.005351338 |
| *MEOX1* | mesenchyme homeobox 1 | -1.04 | 0.047614233 |

| | | | |
|---|---|---|---|
| *DHRS3* | dehydrogenase/reductase 3 | -1.04 | 0.000107485 |
| *AZGP1* | alpha-2-glycoprotein 1, zinc-binding | -1.04 | 0.046154705 |
| *SCUBE1* | signal peptide, CUB domain and EGF like domain containing 1 | -1.04 | 0.048268887 |
| *TF* | transferrin | -1.04 | 0.040204145 |
| *C7* | complement C7 | -1.04 | 0.041428808 |
| *C1QTNF7* | C1q and TNF related 7 | -1.03 | 0.003415192 |
| *CACNB2* | calcium voltage-gated channel auxiliary subunit beta 2 | -1.03 | 0.002505476 |
| *SSH2* | slingshot protein phosphatase 2 | -1.02 | 0.000806423 |
| *ZNF395* | zinc finger protein 395 | -1.02 | 9.78E-09 |
| *EDN1* | endothelin 1 | -1.02 | 0.038653991 |
| *PLEKHH1* | pleckstrin homology, MyTH4 and FERM domain containing H1 | -1.02 | 0.044715424 |
| *NA* | NA | -1.02 | 0.044715424 |
| *GPR158* | G protein-coupled receptor 158 | -1.02 | 0.047205566 |
| *RETREG1* | reticulophagy regulator 1 | -1.01 | 0.030359313 |
| *ANKEF1* | ankyrin repeat and EF-hand domain containing 1 | -1.01 | 0.049854652 |
| *TRNP1* | TMF1 regulated nuclear protein 1 | -1.01 | 0.00255633 |
| *CYP4F22* | cytochrome P450 family 4 subfamily F member 22 | -1.00 | 0.037007094 |
| *HSD17B13* | hydroxysteroid 17-beta dehydrogenase 13 | -1.00 | 0.044705798 |
| *SYNE2* | spectrin repeat containing nuclear envelope protein 2 | -1.00 | 0.023156482 |
| *OLAH* | oleoyl-ACP hydrolase | -1.00 | 0.029149252 |
| *TG* | thyroglobulin | -1.00 | 0.008722934 |
| *WASF3* | WASP family member 3 | -0.99 | 2.89E-07 |
| *NES* | nestin | -0.99 | 0.043040684 |
| *CORO6* | coronin 6 | -0.99 | 0.000191454 |
| *SAMHD1* | SAM and HD domain containing deoxynucleoside triphosphate triphosphohydrolase 1 | -0.99 | 0.027410528 |
| *WNT11* | Wnt family member 11 | -0.99 | 0.029701377 |
| *PPARG* | peroxisome proliferator activated receptor gamma | -0.98 | 0.040510561 |
| *SLC25A10* | solute carrier family 25 member 10 | -0.98 | 0.033690709 |
| *ICOSLG* | inducible T cell costimulator ligand | -0.98 | 0.002446536 |
| *SMARCD2* | SWI/SNF related, matrix associated, actin dependent | -0.97 | 2.37E-11 |

| | | | |
|---|---|---|---|
| | regulator of chromatin, subfamily d, member 2 | | |
| *CTSC* | cathepsin C | -0.96 | 0.012045884 |
| *TUBA8* | tubulin alpha 8 | -0.96 | 0.000438903 |
| *ABCA6* | ATP binding cassette subfamily A member 6 | -0.96 | 0.022413814 |
| *MYCL* | MYCL proto-oncogene, bHLH transcription factor | -0.95 | 0.039950059 |
| *NRCAM* | neuronal cell adhesion molecule | -0.95 | 0.023054091 |
| *STK17B* | serine/threonine kinase 17b | -0.95 | 0.011728543 |
| *GPSM2* | G protein signaling modulator 2 | -0.94 | 0.01272634 |
| *RNASE4* | ribonuclease A family member 4 | -0.94 | 0.000344928 |
| *AOX1* | aldehyde oxidase 1 | -0.94 | 0.009434382 |
| *SPARCL1* | SPARC like 1 | -0.94 | 0.026208715 |
| *LAMA2* | laminin subunit alpha 2 | -0.93 | 0.000662436 |
| *RGS22* | regulator of G protein signaling 22 | -0.93 | 0.015666196 |
| *PHKG1* | phosphorylase kinase catalytic subunit gamma 1 | -0.93 | 0.030359313 |
| *TMEM64* | transmembrane protein 64 | -0.93 | 0.007339432 |
| *ANPEP* | alanyl aminopeptidase, membrane | -0.92 | 0.006783376 |
| *GPATCH11* | G-patch domain containing 11 | -0.92 | 0.036209226 |
| *CDKN1C* | cyclin dependent kinase inhibitor 1C | -0.91 | 0.002000403 |
| *HLX* | H2.0 like homeobox | -0.91 | 4.18E-06 |
| *NEXN* | nexilin F-actin binding protein | -0.89 | 0.011744388 |
| *TGFBR2* | transforming growth factor beta receptor 2 | -0.89 | 2.06E-07 |
| *FMN1* | formin 1 | -0.87 | 0.00839677 |
| *SOX13* | SRY-box transcription factor 13 | -0.87 | 6.66E-06 |
| *IRS1* | insulin receptor substrate 1 | -0.86 | 0.006812536 |
| *ELANE* | elastase, neutrophil expressed | -0.85 | 0.022442794 |
| *SMOC2* | SPARC related modular calcium binding 2 | -0.84 | 0.023365789 |
| *NID1* | nidogen 1 | -0.84 | 0.002123405 |
| *GREB1L* | GREB1 like retinoic acid receptor coactivator | -0.83 | 0.028586024 |
| *PLPP3* | phospholipid phosphatase 3 | -0.83 | 0.024461597 |
| *JADE2* | jade family PHD finger 2 | -0.82 | 0.002505476 |
| *HNMT* | histamine N-methyltransferase | -0.82 | 0.012360534 |
| *NA* | NA | -0.82 | 0.025698089 |

| | | | |
|---|---|---|---|
| *ALDH16A1* | aldehyde dehydrogenase 16 family member A1 | -0.81 | 0.00164913 |
| *IQCH-AS1* | IQCH antisense RNA 1 | -0.81 | 0.006859063 |
| *TNS2* | tensin 2 | -0.81 | 0.003834459 |
| *SLC4A4* | solute carrier family 4 member 4 | -0.81 | 0.047220525 |
| *ANG* | angiogenin | -0.80 | 0.000760663 |
| *NAV2* | neuron navigator 2 | -0.80 | 0.00759564 |
| *TACC1* | transforming acidic coiled-coil containing protein 1 | -0.79 | 5.52E-05 |
| *ADARB1* | adenosine deaminase RNA specific B1 | -0.79 | 0.037699346 |
| *ZNF438* | zinc finger protein 438 | -0.78 | 0.008116872 |
| *LRP4* | LDL receptor related protein 4 | -0.78 | 0.005584825 |
| *TTPAL* | alpha tocopherol transfer protein like | -0.77 | 0.000203943 |
| *LINC00472* | long intergenic non-protein coding RNA 472 | -0.77 | 0.049634866 |
| *LINC01960* | long intergenic non-protein coding RNA 1960 | -0.76 | 0.047614233 |
| *EVA1C* | eva-1 homolog C | -0.76 | 0.041059956 |
| *APOL3* | apolipoprotein L3 | -0.75 | 0.0104071 |
| *TNFAIP8L3* | TNF alpha induced protein 8 like 3 | -0.75 | 0.029107382 |
| *GAS6* | growth arrest specific 6 | -0.75 | 0.005021214 |
| *IL17RE* | interleukin 17 receptor E | -0.75 | 0.043273779 |
| *CHST2* | carbohydrate sulfotransferase 2 | -0.74 | 0.003690124 |
| *PDGFRA* | platelet derived growth factor receptor alpha | -0.74 | 0.002219599 |
| *DIAPH2* | diaphanous related formin 2 | -0.73 | 0.026215254 |
| *BMPR1B* | bone morphogenetic protein receptor type 1B | -0.73 | 0.018281172 |
| *TCEAL4* | transcription elongation factor A like 4 | -0.71 | 0.009681211 |
| *ZHX3* | zinc fingers and homeoboxes 3 | -0.71 | 3.11E-05 |
| *ZBED3* | zinc finger BED-type containing 3 | -0.71 | 0.041059956 |

Supplementary Table 3. DEGs from comparison of active beige and inactive beige adipocytes; n=8

| Symbol | Gene Name | Log2 Fold Change | Adj. P value |
|---|---|---|---|
| *SLC22A12* | solute carrier family 22 member 12 | 1.75 | 5.79E-07 |
| *KRT79* | keratin 79 | 1.69 | 1.40E-06 |
| *PM20D1* | peptidase M20 domain containing 1 | 1.67 | 2.16E-06 |
| *CPA4* | carboxypeptidase A4 | 1.63 | 3.35E-06 |
| *FNDC1* | fibronectin type III domain containing 1 | 1.56 | 3.95E-06 |
| *SCN4A* | sodium voltage-gated channel alpha subunit 4 | 1.56 | 1.70E-05 |
| *AQP3* | aquaporin 3 (Gill blood group) | 1.53 | 5.03E-06 |
| *TENT5C* | terminal nucleotidyltransferase 5C | 1.52 | 1.04E-06 |
| *PCK1* | phosphoenolpyruvate carboxykinase 1 | 1.50 | 3.20E-05 |
| *C1QTNF3* | C1q and TNF related 3 | 1.49 | 1.90E-05 |
| *CD96* | CD96 molecule | 1.47 | 5.81363E-05 |
| *LINC02554* | long intergenic non-protein coding RNA 2554 | 1.47 | 6.38E-05 |
| *C3orf80* | chromosome 3 open reading frame 80 | 1.47 | 1.13E-05 |
| *NA* | NA | 1.46 | 5.29225E-05 |
| *MAP3K7CL* | MAP3K7 C-terminal like | 1.46 | 1.08E-10 |
| *CD52* | CD52 molecule | 1.44 | 9.25E-05 |
| *CLDN2* | claudin 2 | 1.43 | 1.23E-04 |
| *IL1B* | interleukin 1 beta | 1.43 | 4.29E-05 |
| *ADAMTS18* | ADAM metallopeptidase with thrombospondin type 1 motif 18 | 1.42 | 0.000153681 |
| *FCN1* | ficolin 1 | 1.41 | 1.68E-04 |
| *KCNK12* | potassium two pore domain channel subfamily K member 12 | 1.37 | 2.63E-04 |
| *LRRC15* | leucine rich repeat containing 15 | 1.37 | 3.00E-04 |
| *KCNA1* | potassium voltage-gated channel subfamily A member 1 | 1.37 | 7.44808E-05 |
| *SORL1* | sortilin related receptor 1 | 1.36 | 0.000261364 |
| *MYL4* | myosin light chain 4 | 1.34 | 0.000337071 |
| *GK* | glycerol kinase | 1.32 | 1.41E-06 |
| *PPP1R14C* | protein phosphatase 1 regulatory inhibitor subunit 14C | 1.31 | 6.28E-04 |
| *CITED1* | Cbp/p300 interacting transactivator with Glu/Asp rich carboxy-terminal domain 1 | 1.31 | 0.000200952 |
| *BBOX1* | gamma-butyrobetaine hydroxylase 1 | 1.31 | 0.000662964 |

| | | | |
|---|---|---|---|
| *THBD* | thrombomodulin | 1.30 | 1.24E-04 |
| *CTXN1* | cortexin 1 | 1.29 | 1.31E-05 |
| *ADGRG1* | adhesion G protein-coupled receptor G1 | 1.26 | 5.29E-05 |
| *ADAMTS14* | ADAM metallopeptidase with thrombospondin type 1 motif 14 | 1.21 | 5.66E-04 |
| *SLC22A3* | solute carrier family 22 member 3 | 1.21 | 6.26E-05 |
| *PLIN5* | perilipin 5 | 1.20 | 0.00124726 |
| *LOC100506253* | uncharacterized LOC100506253 | 1.18 | 3.39E-03 |
| *RTKN2* | rhotekin 2 | 1.17 | 5.71453E-05 |
| *ACSM3* | acyl-CoA synthetase medium chain family member 3 | 1.17 | 0.00066074 |
| *KCNK15* | potassium two pore domain channel subfamily K member 15 | 1.17 | 1.24E-03 |
| *NA* | NA | 1.17 | 0.003957437 |
| *ECSCR* | endothelial cell surface expressed chemotaxis and apoptosis regulator | 1.16 | 0.003939119 |
| *SLCO2A1* | solute carrier organic anion transporter family member 2A1 | 1.16 | 0.002584458 |
| *CKMT1B* | creatine kinase, mitochondrial 1B | 1.14 | 0.003461073 |
| *LUCAT1* | lung cancer associated transcript 1 | 1.13 | 0.004170862 |
| *GAP43* | growth associated protein 43 | 1.13 | 0.005590778 |
| *RUBCNL* | rubicon like autophagy enhancer | 1.12 | 0.000928559 |
| *RFPL4B* | ret finger protein like 4B | 1.12 | 0.00566713 |
| *MEST* | mesoderm specific transcript | 1.11 | 6.38327E-05 |
| *E2F7* | E2F transcription factor 7 | 1.11 | 3.81167E-05 |
| *CPA2* | carboxypeptidase A2 | 1.11 | 0.007646033 |
| *SCAMP5* | secretory carrier membrane protein 5 | 1.10 | 0.000445285 |
| *RSPO3* | R-spondin 3 | 1.10 | 0.006858304 |
| *CIDEA* | cell death inducing DFFA like effector a | 1.09 | 0.008738219 |
| *ADAM12* | ADAM metallopeptidase domain 12 | 1.09 | 0.001696744 |
| *SELPLG* | selectin P ligand | 1.08 | 7.39E-05 |
| *SLC7A10* | solute carrier family 7 member 10 | 1.08 | 0.009565079 |
| *CIART* | circadian associated repressor of transcription | 1.08 | 2.43E-08 |
| *EPB41L4B* | erythrocyte membrane protein band 4.1 like 4B | 1.08 | 0.000915872 |
| *ADAMTS16* | ADAM metallopeptidase with thrombospondin type 1 motif 16 | 1.08 | 0.007957327 |

| | | | |
|---|---|---|---|
| *AATBC* | apoptosis associated transcript in bladder cancer | 1.07 | 0.010616214 |
| *LAMP3* | lysosomal associated membrane protein 3 | 1.06 | 0.012656697 |
| *PPP1R1A* | protein phosphatase 1 regulatory inhibitor subunit 1A | 1.06 | 0.011764285 |
| *FABP4* | fatty acid binding protein 4 | 1.05 | 0.010889907 |
| *TSHR* | thyroid stimulating hormone receptor | 1.05 | 0.012640386 |
| *RPS6KA1* | ribosomal protein S6 kinase A1 | 1.04 | 0.00720311 |
| *CKMT1A* | creatine kinase, mitochondrial 1A | 1.04 | 0.01438844 |
| *LAMC3* | laminin subunit gamma 3 | 1.04 | 0.010130293 |
| *PHLDA1* | pleckstrin homology like domain family A member 1 | 1.02 | 0.004202125 |
| *TRPM8* | transient receptor potential cation channel subfamily M member 8 | 1.01 | 0.019770198 |
| *RPLP0P2* | ribosomal protein lateral stalk subunit P0 pseudogene 2 | 1.01 | 0.003176049 |
| *KIRREL3* | kirre like nephrin family adhesion molecule 3 | 1.01 | 0.018804338 |
| *DPYSL3* | dihydropyrimidinase like 3 | 1.00 | 0.002259103 |
| *OPRK1* | opioid receptor kappa 1 | 0.99 | 0.020054477 |
| *PRG4* | proteoglycan 4 | 0.99 | 0.011534645 |
| *DLX3* | distal-less homeobox 3 | 0.99 | 0.022254759 |
| *FER1L6* | fer-1 like family member 6 | 0.99 | 0.025850071 |
| *NUDT14* | nudix hydrolase 14 | 0.99 | 1.14E-07 |
| *ICA1* | islet cell autoantigen 1 | 0.99 | 0.009355549 |
| *SH3GL3* | SH3 domain containing GRB2 like 3, endophilin A3 | 0.99 | 0.025960556 |
| *CEMIP* | cell migration inducing hyaluronidase 1 | 0.99 | 0.008061658 |
| *RIMS4* | regulating synaptic membrane exocytosis 4 | 0.96 | 2.31E-02 |
| *ZNF365* | zinc finger protein 365 | 0.96 | 0.012640386 |
| *DOK5* | docking protein 5 | 0.96 | 6.72E-12 |
| *IL13RA2* | interleukin 13 receptor subunit alpha 2 | 0.96 | 0.03383346 |
| *CYP4F12* | cytochrome P450 family 4 subfamily F member 12 | 0.96 | 0.030761867 |
| *PDE1B* | phosphodiesterase 1B | 0.95 | 0.032617469 |
| *PALD1* | phosphatase domain containing paladin 1 | 0.95 | 0.028348773 |
| *CH25H* | cholesterol 25-hydroxylase | 0.95 | 0.030970829 |

| | | | |
|---|---|---|---|
| *LMO2* | LIM domain only 2 | 0.95 | 0.000818633 |
| *MYOM3* | myomesin 3 | 0.95 | 0.03593656 |
| *INA* | internexin neuronal intermediate filament protein alpha | 0.95 | 0.027731101 |
| *TMEM63C* | transmembrane protein 63C | 0.94 | 0.037583312 |
| *VAT1L* | vesicle amine transport 1 like | 0.94 | 3.89E-02 |
| *TMEM132C* | transmembrane protein 132C | 0.94 | 0.035016534 |
| *CSRP2* | cysteine and glycine rich protein 2 | 0.94 | 0.018804338 |
| *LHCGR* | luteinizing hormone/choriogonadotropin receptor | 0.94 | 0.040842846 |
| *INPP5J* | inositol polyphosphate-5-phosphatase J | 0.93 | 3.29E-02 |
| *COBL* | cordon-bleu WH2 repeat protein | 0.93 | 0.044827867 |
| *EVA1A* | eva-1 homolog A, regulator of programmed cell death | 0.93 | 0.019781645 |
| *FAM151A* | family with sequence similarity 151 member A | 0.92 | 0.046370483 |
| *SSC5D* | scavenger receptor cysteine rich family member with 5 domains | 0.92 | 0.002407721 |
| *SLC24A2* | solute carrier family 24 member 2 | 0.92 | 4.13E-02 |
| *STX11* | syntaxin 11 | 0.91 | 9.28E-04 |
| *EPHB2* | EPH receptor B2 | 0.91 | 0.002996158 |
| *CYS1* | cystin 1 | 0.91 | 1.25E-03 |
| *DIRAS1* | DIRAS family GTPase 1 | 0.90 | 0.006008681 |
| *DIRC3* | disrupted in renal carcinoma 3 | 0.90 | 0.041835677 |
| *PPP2R2C* | protein phosphatase 2 regulatory subunit Bgamma | 0.90 | 0.028348773 |
| *LGI2* | leucine rich repeat LGI family member 2 | 0.89 | 0.042654121 |
| *ARC* | activity regulated cytoskeleton associated protein | 0.89 | 0.032056619 |
| *AP3B2* | adaptor related protein complex 3 subunit beta 2 | 0.89 | 0.047838824 |
| *LOC100507006* | uncharacterized LOC100507006 | 0.88 | 0.040407218 |
| *RHPN1* | rhophilin Rho GTPase binding protein 1 | 0.88 | 0.046271955 |
| *ABCA3* | ATP binding cassette subfamily A member 3 | 0.86 | 0.013166092 |
| *PLXDC1* | plexin domain containing 1 | 0.86 | 0.011636014 |
| *GXYLT2* | glucoside xylosyltransferase 2 | 0.85 | 0.002284371 |
| *SALL4* | spalt like transcription factor 4 | 0.85 | 0.032617469 |
| *TUBB3* | tubulin beta 3 class III | 0.84 | 0.008271003 |

| | | | |
|---|---|---|---|
| *ABCG1* | ATP binding cassette subfamily G member 1 | 0.84 | 0.017863114 |
| *LIF* | LIF interleukin 6 family cytokine | 0.82 | 0.012448106 |
| *SEL1L3* | SEL1L family member 3 | 0.82 | 0.026742566 |
| *PER3* | period circadian regulator 3 | 0.81 | 0.006539207 |
| *EGFLAM* | EGF like, fibronectin type III and laminin G domains | 0.81 | 0.027873852 |
| *ADGRB2* | adhesion G protein-coupled receptor B2 | 0.81 | 0.016609283 |
| *ASPHD2* | aspartate beta-hydroxylase domain containing 2 | 0.81 | 0.003141574 |
| *EPHA2* | EPH receptor A2 | 0.80 | 0.032617469 |
| *LTC4S* | leukotriene C4 synthase | 0.80 | 0.040842846 |
| *LACC1* | laccase domain containing 1 | 0.80 | 0.00175865 |
| *RBM24* | RNA binding motif protein 24 | 0.80 | 0.041264278 |
| *PLEKHA7* | pleckstrin homology domain containing A7 | 0.80 | 0.041264278 |
| *RGL3* | ral guanine nucleotide dissociation stimulator like 3 | 0.79 | 0.048585751 |
| *DUSP8* | dual specificity phosphatase 8 | 0.79 | 0.007515436 |
| *TDRKH* | tudor and KH domain containing | 0.78 | 0.003107981 |
| *EPB41* | erythrocyte membrane protein band 4.1 | 0.78 | 0.004170862 |
| *GPC1* | glypican 1 | 0.78 | 0.002463807 |
| *FUCA1* | alpha-L-fucosidase 1 | 0.77 | 0.000696625 |
| *CAMKK1* | calcium/calmodulin dependent protein kinase kinase 1 | 0.77 | 0.006354175 |
| *HOMER2* | homer scaffold protein 2 | 0.76 | 0.019363023 |
| *LUM* | lumican | 0.76 | 0.004412096 |
| *ASS1* | argininosuccinate synthase 1 | 0.75 | 0.001070742 |
| *TGM5* | transglutaminase 5 | 0.74 | 0.028464297 |
| *KIAA1549L* | KIAA1549 like | 0.74 | 0.000270804 |
| *PLAU* | plasminogen activator, urokinase | 0.73 | 0.001151066 |
| *AKR1B1* | aldo-keto reductase family 1 member B | 0.71 | 0.018470855 |
| *ARNTL2* | aryl hydrocarbon receptor nuclear translocator like 2 | 0.71 | 0.026616037 |
| *STARD10* | StAR related lipid transfer domain containing 10 | 0.70 | 0.013173834 |
| *ZBTB16* | zinc finger and BTB domain containing 16 | -4.55 | 7.5279E-135 |
| *HIF3A* | hypoxia inducible factor 3 subunit alpha | -2.80 | 6.48078E-20 |

| | | | |
|---|---|---|---|
| *FKBP5* | FKBP prolyl isomerase 5 | -2.63 | 7.43547E-36 |
| *PER1* | period circadian regulator 1 | -2.48 | 1.05052E-66 |
| *PILRA* | paired immunoglobin like type 2 receptor alpha | -2.41 | 7.9946E-20 |
| *INHBB* | inhibin subunit beta B | -2.04 | 1.0022E-16 |
| *MAOA* | monoamine oxidase A | -1.96 | 6.13575E-15 |
| *TIMP4* | TIMP metallopeptidase inhibitor 4 | -1.94 | 6.0412E-09 |
| *CRISPLD2* | cysteine rich secretory protein LCCL domain containing 2 | -1.90 | 3.8403E-17 |
| *SLC16A12* | solute carrier family 16 member 12 | -1.90 | 2.56274E-08 |
| *NEGR1* | neuronal growth regulator 1 | -1.86 | 3.01063E-22 |
| *GGT5* | gamma-glutamyltransferase 5 | -1.85 | 4.76338E-09 |
| *RAPGEF5* | Rap guanine nucleotide exchange factor 5 | -1.82 | 1.43402E-07 |
| *ACKR2* | atypical chemokine receptor 2 | -1.77 | 2.67498E-12 |
| *ABCC2* | ATP binding cassette subfamily C member 2 | -1.76 | 2.06239E-07 |
| *LMO3* | LIM domain only 3 | -1.76 | 5.50293E-12 |
| *RASL11A* | RAS like family 11 member A | -1.76 | 5.53268E-13 |
| *NA* | NA | -1.74 | 5.92002E-07 |
| *PRODH* | proline dehydrogenase 1 | -1.72 | 4.50E-07 |
| *TMC2* | transmembrane channel like 2 | -1.68 | 2.41386E-07 |
| *GALNT15* | polypeptide N-acetylgalactosaminyltransferase 15 | -1.67 | 1.91091E-06 |
| *GRIA1* | glutamate ionotropic receptor AMPA type subunit 1 | -1.65 | 7.5525E-07 |
| *CYP4B1* | cytochrome P450 family 4 subfamily B member 1 | -1.65 | 1.41023E-06 |
| *C6* | complement C6 | -1.64 | 3.84958E-06 |
| *DPEP1* | dipeptidase 1 | -1.64 | 3.95E-06 |
| *LRP1B* | LDL receptor related protein 1B | -1.63 | 3.84958E-06 |
| *CYP8B1* | cytochrome P450 family 8 subfamily B member 1 | -1.63 | 3.16197E-08 |
| *FMO2* | flavin containing dimethylaniline monoxygenase 2 | -1.59 | 1.25734E-06 |
| *PLXNA4* | plexin A4 | -1.54 | 4.73626E-11 |
| *ADRA1B* | adrenoceptor alpha 1B | -1.51 | 2.70604E-07 |
| *ANGPTL1* | angiopoietin like 1 | -1.50 | 1.31348E-11 |
| *DPT* | dermatopontin | -1.50 | 2.97073E-06 |
| *STC1* | stanniocalcin 1 | -1.47 | 5.10E-05 |
| *MGP* | matrix Gla protein | -1.46 | 2.07477E-05 |
| *CRLF1* | cytokine receptor like factor 1 | -1.44 | 1.50582E-08 |

| | | | |
|---|---|---|---|
| *TENT5B* | terminal nucleotidyltransferase 5B | -1.43 | 1.02619E-07 |
| *ANGPTL8* | angiopoietin like 8 | -1.42 | 1.15E-04 |
| *F5* | coagulation factor V | -1.41 | 1.63E-04 |
| *TSC22D3* | TSC22 domain family member 3 | -1.40 | 4.10922E-25 |
| *GABRA5* | gamma-aminobutyric acid type A receptor subunit alpha5 | -1.40 | 1.68E-04 |
| *LOC285847* | uncharacterized LOC285847 | -1.40 | 1.23E-04 |
| *GPX3* | glutathione peroxidase 3 | -1.40 | 1.75E-04 |
| *LEP* | leptin | -1.40 | 1.75E-04 |
| *AVPR1A* | arginine vasopressin receptor 1A | -1.36 | 3.37E-04 |
| *SAA1* | serum amyloid A1 | -1.35 | 1.57E-04 |
| *CPM* | carboxypeptidase M | -1.35 | 2.60E-04 |
| *MMP28* | matrix metallopeptidase 28 | -1.34 | 1.23E-04 |
| *KIAA0040* | KIAA0040 | -1.34 | 3.50E-04 |
| *POM121L9P* | POM121 transmembrane nucleoporin like 9, pseudogene | -1.34 | 1.49E-04 |
| *FGD4* | FYVE, RhoGEF and PH domain containing 4 | -1.33 | 1.07333E-05 |
| *ALOX5AP* | arachidonate 5-lipoxygenase activating protein | -1.31 | 6.15E-04 |
| *GLUL* | glutamate-ammonia ligase | -1.30 | 4.42008E-07 |
| *RGCC* | regulator of cell cycle | -1.27 | 0.000915872 |
| *WNT5A* | Wnt family member 5A | -1.27 | 5.74E-05 |
| *EPHB6* | EPH receptor B6 | -1.27 | 4.32E-04 |
| *SLA* | Src like adaptor | -1.26 | 1.12E-03 |
| *CCN4* | cellular communication network factor 4 | -1.26 | 9.98E-04 |
| *LRRN3* | leucine rich repeat neuronal 3 | -1.26 | 8.26E-04 |
| *HPD* | 4-hydroxyphenylpyruvate dioxygenase | -1.24 | 0.000893404 |
| *SYN2* | synapsin II | -1.23 | 1.58E-03 |
| *CLCN1* | chloride voltage-gated channel 1 | -1.23 | 1.51E-03 |
| *SLPI* | secretory leukocyte peptidase inhibitor | -1.23 | 0.001085974 |
| *VIT* | vitrin | -1.23 | 3.01E-04 |
| *METTL7A* | methyltransferase like 7A | -1.21 | 0.000737103 |
| *PTGS1* | prostaglandin-endoperoxide synthase 1 | -1.21 | 4.26491E-07 |
| *APCDD1* | APC down-regulated 1 | -1.21 | 0.00124726 |
| *KLF9* | Kruppel like factor 9 | -1.20 | 4.70306E-23 |
| *TMPRSS5* | transmembrane serine protease 5 | -1.19 | 0.00308391 |
| *ART4* | ADP-ribosyltransferase 4 (inactive) (Dombrock blood group) | -1.17 | 0.003269535 |

| | | | |
|---|---|---|---|
| *GPM6B* | glycoprotein M6B | -1.17 | 8.75E-04 |
| *ERRFI1* | ERBB receptor feedback inhibitor 1 | -1.16 | 8.19E-09 |
| *SMCO3* | single-pass membrane protein with coiled-coil domains 3 | -1.15 | 0.001962714 |
| *PKD2L1* | polycystin 2 like 1, transient receptor potential cation channel | -1.15 | 0.004170862 |
| *ALKAL2* | ALK and LTK ligand 2 | -1.14 | 0.005150176 |
| *PIK3R1* | phosphoinositide-3-kinase regulatory subunit 1 | -1.13 | 5.24E-04 |
| *LINC01088* | long intergenic non-protein coding RNA 1088 | -1.12 | 0.006055842 |
| *APOD* | apolipoprotein D | -1.12 | 0.006846304 |
| *TG* | thyroglobulin | -1.12 | 0.000915872 |
| *TC2N* | tandem C2 domains, nuclear | -1.12 | 1.16E-03 |
| *SPARCL1* | SPARC like 1 | -1.11 | 0.007330778 |
| *RGMA* | repulsive guidance molecule BMP co-receptor a | -1.11 | 1.15509E-12 |
| *OMD* | osteomodulin | -1.11 | 0.004922461 |
| *C1QTNF7* | C1q and TNF related 7 | -1.10 | 0.000915872 |
| *TRNP1* | TMF1 regulated nuclear protein 1 | -1.10 | 6.58E-05 |
| *ATP1A2* | ATPase Na+/K+ transporting subunit alpha 2 | -1.10 | 0.005072343 |
| *CDKN1C* | cyclin dependent kinase inhibitor 1C | -1.10 | 2.40E-04 |
| *MT1X* | metallothionein 1X | -1.10 | 8.76771E-08 |
| *CST3* | cystatin C | -1.09 | 3.30E-05 |
| *OLAH* | oleoyl-ACP hydrolase | -1.09 | 6.50E-03 |
| *ISM1* | isthmin 1 | -1.08 | 0.009823468 |
| *ZNF385B* | zinc finger protein 385B | -1.08 | 0.008080662 |
| *EDN1* | endothelin 1 | -1.07 | 0.005995502 |
| *FAM166B* | family with sequence similarity 166 member B | -1.07 | 0.00842496 |
| *NOSTRIN* | nitric oxide synthase trafficking | -1.07 | 0.005427007 |
| *CACNB2* | calcium voltage-gated channel auxiliary subunit beta 2 | -1.06 | 5.31E-05 |
| *ADH1A* | alcohol dehydrogenase 1A (class I), alpha polypeptide | -1.06 | 0.008080662 |
| *NA* | NA | -1.06 | 2.92E-04 |
| *STK17B* | serine/threonine kinase 17b | -1.06 | 0.00073653 |
| *WASF3* | WASP family member 3 | -1.06 | 1.8619E-18 |
| *POU3F3* | POU class 3 homeobox 3 | -1.05 | 0.011501075 |

| | | | |
|---|---|---|---|
| *SYNE2* | spectrin repeat containing nuclear envelope protein 2 | -1.05 | 0.005255904 |
| *HLX* | H2.0 like homeobox | -1.04 | 5.50293E-12 |
| *SOX13* | SRY-box transcription factor 13 | -1.04 | 1.31011E-08 |
| *ADARB1* | adenosine deaminase RNA specific B1 | -1.03 | 0.001917124 |
| *GREB1L* | GREB1 like retinoic acid receptor coactivator | -1.03 | 5.35584E-06 |
| *SERPINA3* | serpin family A member 3 | -1.03 | 0.013858835 |
| *SMOC2* | SPARC related modular calcium binding 2 | -1.03 | 0.002584458 |
| *GLYAT* | glycine-N-acyltransferase | -1.02 | 0.014558379 |
| *MAP2K6* | mitogen-activated protein kinase kinase 6 | -1.01 | 0.015888699 |
| *ANGPTL5* | angiopoietin like 5 | -1.01 | 0.019781645 |
| *NA* | NA | -1.01 | 0.004317435 |
| *AOX1* | aldehyde oxidase 1 | -1.00 | 0.00062806 |
| *EYA2* | EYA transcriptional coactivator and phosphatase 2 | -1.00 | 0.012916704 |
| *CCDC54* | coiled-coil domain containing 54 | -1.00 | 2.25E-02 |
| *NRCAM* | neuronal cell adhesion molecule | -1.00 | 0.008061658 |
| *TRABD2B* | TraB domain containing 2B | -1.00 | 0.013858835 |
| *NEBL* | nebulette | -1.00 | 0.021379713 |
| *RASSF4* | Ras association domain family member 4 | -1.00 | 3.84958E-06 |
| *RIPOR3* | RIPOR family member 3 | -1.00 | 0.024466138 |
| *CYP4Z2P* | cytochrome P450 family 4 subfamily Z member 2, pseudogene | -0.99 | 0.019980284 |
| *ADH1B* | alcohol dehydrogenase 1B (class I), beta polypeptide | -0.99 | 0.019748554 |
| *ROBO2* | roundabout guidance receptor 2 | -0.99 | 2.24E-02 |
| *ANPEP* | alanyl aminopeptidase, membrane | -0.98 | 1.93E-04 |
| *NGFR* | nerve growth factor receptor | -0.97 | 0.012356687 |
| *DHRS3* | dehydrogenase/reductase 3 | -0.97 | 1.67508E-05 |
| *PPP4R4* | protein phosphatase 4 regulatory subunit 4 | -0.97 | 0.028182462 |
| *NID1* | nidogen 1 | -0.97 | 3.67673E-11 |
| *TACC1* | transforming acidic coiled-coil containing protein 1 | -0.97 | 1.08525E-08 |
| *LAMA2* | laminin subunit alpha 2 | -0.97 | 4.88473E-06 |
| *ZNF395* | zinc finger protein 395 | -0.97 | 7.15001E-13 |
| *TNFSF10* | TNF superfamily member 10 | -0.97 | 0.016609283 |

| | | | |
|---|---|---|---|
| *NPY5R* | neuropeptide Y receptor Y5 | -0.97 | 0.027192915 |
| *SPATA41* | spermatogenesis associated 41 | -0.96 | 0.022419483 |
| *CFD* | complement factor D | -0.96 | 0.00285215 |
| *CORO6* | coronin 6 | -0.95 | 5.06E-04 |
| *RNF144B* | ring finger protein 144B | -0.95 | 0.027643637 |
| *NPR3* | natriuretic peptide receptor 3 | -0.95 | 0.009253616 |
| *FRZB* | frizzled related protein | -0.94 | 0.038107473 |
| *IP6K3* | inositol hexakisphosphate kinase 3 | -0.94 | 0.034988853 |
| *TRPV6* | transient receptor potential cation channel subfamily V member 6 | -0.94 | 0.037485797 |
| *NEXN* | nexilin F-actin binding protein | -0.94 | 0.002125546 |
| *HPSE* | heparanase | -0.94 | 0.037619397 |
| *FAM107A* | family with sequence similarity 107 member A | -0.94 | 0.041835677 |
| *RXFP1* | relaxin family peptide receptor 1 | -0.93 | 0.042064995 |
| *SMARCD2* | SWI/SNF related, matrix associated, actin dependent regulator of chromatin, subfamily d, member 2 | -0.93 | 4.03555E-17 |
| *ARMC12* | armadillo repeat containing 12 | -0.93 | 0.009686755 |
| *DAAM2-AS1* | DAAM2 antisense RNA 1 | -0.93 | 0.01375562 |
| *BOC* | BOC cell adhesion associated, oncogene regulated | -0.93 | 4.65E-03 |
| *NA* | NA | -0.93 | 0.040280158 |
| *NA* | NA | -0.93 | 0.037007018 |
| *SSH2* | slingshot protein phosphatase 2 | -0.93 | 6.58E-05 |
| *CTHRC1* | collagen triple helix repeat containing 1 | -0.93 | 0.007646033 |
| *SLC2A13* | solute carrier family 2 member 13 | -0.92 | 0.005695219 |
| *NPY1R* | neuropeptide Y receptor Y1 | -0.91 | 0.028464297 |
| *AFAP1L1* | actin filament associated protein 1 like 1 | -0.91 | 0.041186015 |
| *SAMHD1* | SAM and HD domain containing deoxynucleoside triphosphate triphosphohydrolase 1 | -0.91 | 0.024527896 |
| *IQCH-AS1* | IQCH antisense RNA 1 | -0.91 | 2.56892E-05 |
| *OLFM2* | olfactomedin 2 | -0.90 | 4.10E-05 |
| *NAV2* | neuron navigator 2 | -0.90 | 6.20E-05 |
| *TMEM150C* | transmembrane protein 150C | -0.89 | 0.032617469 |
| *GRK5* | G protein-coupled receptor kinase 5 | -0.89 | 0.00166459 |
| *TGFBR2* | transforming growth factor beta receptor 2 | -0.89 | 3.94342E-32 |
| *TNS2* | tensin 2 | -0.89 | 1.27664E-07 |

| | | | |
|---|---|---|---|
| *CAB39L* | calcium binding protein 39 like | -0.89 | 1.23606E-05 |
| *TMEM64* | transmembrane protein 64 | -0.89 | 0.001473297 |
| *ICOSLG* | inducible T cell costimulator ligand | -0.88 | 4.09E-03 |
| *DNASE1L3* | deoxyribonuclease 1 like 3 | -0.88 | 0.023948217 |
| *CUTC* | cutC copper transporter | -0.88 | 4.42008E-07 |
| *TNNT3* | troponin T3, fast skeletal type | -0.88 | 3.21E-02 |
| *TUBA8* | tubulin alpha 8 | -0.88 | 6.78E-04 |
| *RAB40A* | RAB40A, member RAS oncogene family | -0.88 | 0.011749104 |
| *HNMT* | histamine N-methyltransferase | -0.87 | 0.006531785 |
| *GSN* | gelsolin | -0.87 | 0.034022986 |
| *LINC00921* | long intergenic non-protein coding RNA 921 | -0.86 | 6.21E-04 |
| *TCEAL4* | transcription elongation factor A like 4 | -0.85 | 9.01E-04 |
| *DNAJC6* | DnaJ heat shock protein family (Hsp40) member C6 | -0.85 | 0.007612416 |
| *RNASE4* | ribonuclease A family member 4 | -0.85 | 6.26E-05 |
| *IRS2* | insulin receptor substrate 2 | -0.85 | 0.002256864 |
| *CSRNP3* | cysteine and serine rich nuclear protein 3 | -0.85 | 0.037619397 |
| *GUCY1B1* | guanylate cyclase 1 soluble subunit beta 1 | -0.84 | 0.019781645 |
| *LAMA3* | laminin subunit alpha 3 | -0.84 | 4.55E-03 |
| *CTSC* | cathepsin C | -0.84 | 0.013166092 |
| *TMEM119* | transmembrane protein 119 | -0.82 | 0.043667191 |
| *BMPR1B* | bone morphogenetic protein receptor type 1B | -0.82 | 0.002799133 |
| *EVA1C* | eva-1 homolog C | -0.82 | 2.98E-03 |
| *FOXO1* | forkhead box O1 | -0.81 | 7.67E-04 |
| *LINC00968* | long intergenic non-protein coding RNA 968 | -0.80 | 0.037925198 |
| *ABCA6* | ATP binding cassette subfamily A member 6 | -0.80 | 0.047838824 |
| *FBLN2* | fibulin 2 | -0.79 | 1.81E-04 |
| *RHOBTB3* | Rho related BTB domain containing 3 | -0.79 | 0.013385188 |
| *ZHX3* | zinc fingers and homeoboxes 3 | -0.79 | 7.56081E-06 |
| *ELANE* | elastase, neutrophil expressed | -0.79 | 0.019363023 |
| *TLR4* | toll like receptor 4 | -0.79 | 0.000798292 |
| *FBLN5* | fibulin 5 | -0.79 | 0.047104552 |
| *JAK2* | Janus kinase 2 | -0.79 | 1.56E-04 |
| *SERPING1* | serpin family G member 1 | -0.79 | 4.38E-04 |

| | | | |
|---|---|---|---|
| *PLPP3* | phospholipid phosphatase 3 | -0.78 | 0.00240489 |
| *PDGFRA* | platelet derived growth factor receptor alpha | -0.78 | 1.28E-04 |
| *FUT8-AS1* | FUT8 antisense RNA 1 | -0.77 | 0.030646591 |
| *ANG* | angiogenin | -0.77 | 2.18E-06 |
| *PROSER2* | proline and serine rich 2 | -0.77 | 0.03585722 |
| *TMEM158* | transmembrane protein 158 | -0.76 | 0.012916704 |
| *NABP1* | nucleic acid binding protein 1 | -0.76 | 0.00185271 |
| *MBNL2* | muscleblind like splicing regulator 2 | -0.76 | 0.002457234 |
| *CALCOCO2* | calcium binding and coiled-coil domain 2 | -0.76 | 2.53E-04 |
| *DSEL* | dermatan sulfate epimerase like | -0.75 | 0.027172422 |
| *APOL3* | apolipoprotein L3 | -0.75 | 0.005398914 |
| *SH3D19* | SH3 domain containing 19 | -0.75 | 0.00842496 |
| *SYP* | synaptophysin | -0.75 | 0.043274778 |
| *PLCL1* | phospholipase C like 1 (inactive) | -0.74 | 0.018046668 |
| *ZFP36* | ZFP36 ring finger protein | -0.74 | 2.36207E-06 |
| *SLC4A11* | solute carrier family 4 member 11 | -0.74 | 3.87E-02 |
| *PID1* | phosphotyrosine interaction domain containing 1 | -0.74 | 0.037987105 |
| *GAS6* | growth arrest specific 6 | -0.73 | 0.000983242 |
| *NA* | NA | -0.73 | 0.008061658 |
| *EPN2* | epsin 2 | -0.72 | 6.38E-04 |
| *IRS1* | insulin receptor substrate 1 | -0.72 | 3.37E-04 |
| *PDE8B* | phosphodiesterase 8B | -0.72 | 0.022575341 |
| *KLF13* | Kruppel like factor 13 | -0.72 | 4.11E-05 |
| *FOXO3* | forkhead box O3 | -0.71 | 1.92007E-09 |
| *ARRDC2* | arrestin domain containing 2 | -0.71 | 0.00062806 |
| *MFAP5* | microfibril associated protein 5 | -0.71 | 0.018804338 |
| *ZEB1* | zinc finger E-box binding homeobox 1 | -0.70 | 0.021504341 |
| *MPP3* | membrane palmitoylated protein 3 | -0.70 | 1.12E-02 |
| *TRAF1* | TNF receptor associated factor 1 | -0.70 | 2.52E-04 |
| *LRP4* | LDL receptor related protein 4 | -0.70 | 3.71E-03 |

Supplementary Table 4 DEGs from comparison of active beige adipocytes carrying *FTO* obesity-risk vs risk-free genotypes; n=4 of each genotype

| Symbol | Gene Name | Log2 Fold Change | Adj. P value |
|---|---|---|---|
| *MAOB* | monoamine oxidase B | -2.03 | 2.99E-16 |
| *NKD2* | NKD inhibitor of WNT signaling pathway 2 | -1.86 | 8.11E-06 |
| *C16orf89* | chromosome 16 open reading frame 89 | -1.84 | 1.35E-05 |
| *PLXNC1* | plexin C1 | -1.82 | 1.18E-04 |
| *MAPK8IP2* | mitogen-activated protein kinase 8 interacting protein 2 | -1.74 | 8.14E-11 |
| *SYT17* | synaptotagmin 17 | -1.74 | 1.18E-04 |
| *CDHR1* | cadherin related family member 1 | -1.74 | 2.38E-04 |
| *ABHD1* | abhydrolase domain containing 1 | -1.68 | 2.52E-04 |
| *CKMT2* | creatine kinase, mitochondrial 2 | -1.62 | 5.75E-04 |
| *SIM1* | SIM bHLH transcription factor 1 | -1.61 | 4.11E-04 |
| *PPP1R1B* | protein phosphatase 1 regulatory inhibitor subunit 1B | -1.61 | 0.000575113 |
| *UCP2* | uncoupling protein 2 | -1.60 | 2.38E-04 |
| *ZNF385A* | zinc finger protein 385A | -1.58 | 1.29E-08 |
| *LDHD* | lactate dehydrogenase D | -1.57 | 0.000118498 |
| *ANO5* | anoctamin 5 | -1.56 | 2.01E-03 |
| *RBM47* | RNA binding motif protein 47 | -1.51 | 2.38E-04 |
| *MYH14* | myosin heavy chain 14 | -1.50 | 1.18E-04 |
| *ALDH3B2* | aldehyde dehydrogenase 3 family member B2 | -1.50 | 4.44E-03 |
| *KCNB1* | potassium voltage-gated channel subfamily B member 1 | -1.49 | 0.002342861 |
| *RAMP2-AS1* | RAMP2 antisense RNA 1 | -1.49 | 2.76E-03 |
| *TRDC* | T cell receptor delta constant | -1.47 | 5.17E-03 |
| *ERV3-1* | endogenous retrovirus group 3 member 1, envelope | -1.47 | 1.40E-04 |
| *MAPK10* | mitogen-activated protein kinase 10 | -1.45 | 0.001513359 |
| *SCD* | stearoyl-CoA desaturase | -1.42 | 0.002342861 |
| *ACACB* | acetyl-CoA carboxylase beta | -1.42 | 0.002763087 |
| *NR1H3* | nuclear receptor subfamily 1 group H member 3 | -1.42 | 5.57E-03 |
| *CLIC6* | chloride intracellular channel 6 | -1.41 | 5.17E-03 |
| *STON2* | stonin 2 | -1.41 | 0.003146207 |
| *KIF1A* | kinesin family member 1A | -1.41 | 0.007671087 |
| *GPAM* | glycerol-3-phosphate acyltransferase, mitochondrial | -1.41 | 6.74E-03 |
| *PRKG2* | protein kinase cGMP-dependent 2 | -1.40 | 8.34E-03 |

| | | | |
|---|---|---|---|
| *C1orf115* | chromosome 1 open reading frame 115 | -1.39 | 8.23E-03 |
| *CES1* | carboxylesterase 1 | -1.39 | 3.98E-03 |
| *GPD1* | glycerol-3-phosphate dehydrogenase 1 | -1.38 | 7.97E-03 |
| *APCDD1* | APC down-regulated 1 | -1.38 | 9.7593E-05 |
| *ELOVL6* | ELOVL fatty acid elongase 6 | -1.38 | 1.05E-02 |
| *FCN1* | ficolin 1 | -1.38 | 0.010962781 |
| *AQP7P1* | aquaporin 7 pseudogene 1 | -1.37 | 0.007384258 |
| *PLAAT5* | phospholipase A and acyltransferase 5 | -1.36 | 9.42E-03 |
| *PPP1R1A* | protein phosphatase 1 regulatory inhibitor subunit 1A | -1.36 | 0.007918342 |
| *S100B* | S100 calcium binding protein B | -1.35 | 0.015108531 |
| *CDH23* | cadherin related 23 | -1.34 | 0.016546538 |
| *RET* | ret proto-oncogene | -1.34 | 0.013288966 |
| *LHCGR* | luteinizing hormone/choriogonadotropin receptor | -1.34 | 0.014115139 |
| *ACSF2* | acyl-CoA synthetase family member 2 | -1.32 | 0.003227973 |
| *GRK3* | G protein-coupled receptor kinase 3 | -1.30 | 0.011607392 |
| *MOGAT1* | monoacylglycerol O-acyltransferase 1 | -1.30 | 0.021836614 |
| *ME1* | malic enzyme 1 | -1.30 | 0.00477724 |
| *CIDEC* | cell death inducing DFFA like effector c | -1.29 | 0.013532874 |
| *SHMT1* | serine hydroxymethyltransferase 1 | -1.28 | 0.010334498 |
| *PRKAR2B* | protein kinase cAMP-dependent type II regulatory subunit beta | -1.28 | 0.01008547 |
| *SORL1* | sortilin related receptor 1 | -1.28 | 0.017034961 |
| *CDK15* | cyclin dependent kinase 15 | -1.27 | 0.021836614 |
| *LGALS12* | galectin 12 | -1.27 | 0.018978932 |
| *PXDNL* | peroxidasin like | -1.27 | 2.71E-02 |
| *SLC29A4* | solute carrier family 29 member 4 | -1.27 | 1.32603E-06 |
| *ADIPOQ* | adiponectin, C1Q and collagen domain containing | -1.26 | 2.54E-02 |
| *GYG2* | glycogenin 2 | -1.26 | 0.003089649 |
| *PLEKHG1* | pleckstrin homology and RhoGEF domain containing G1 | -1.26 | 0.022551038 |
| *USP51* | ubiquitin specific peptidase 51 | -1.26 | 0.003089649 |
| *CLMN* | calmin | -1.25 | 0.02264628 |
| *PIWIL2* | piwi like RNA-mediated gene silencing 2 | -1.24 | 0.032058082 |
| *FGR* | FGR proto-oncogene, Src family tyrosine kinase | -1.24 | 0.030852179 |
| *PDE3B* | phosphodiesterase 3B | -1.24 | 0.028787624 |
| *APOE* | apolipoprotein E | -1.23 | 0.01729454 |

| | | | |
|---|---|---|---|
| *NNAT* | neuronatin | -1.23 | 0.008226534 |
| *ACSS2* | acyl-CoA synthetase short chain family member 2 | -1.23 | 0.022483074 |
| *SV2A* | synaptic vesicle glycoprotein 2A | -1.22 | 0.031596998 |
| *C14orf180* | chromosome 14 open reading frame 180 | -1.22 | 0.027149535 |
| *PCSK1N* | proprotein convertase subtilisin/kexin type 1 inhibitor | -1.22 | 0.035904464 |
| *SLC47A1* | solute carrier family 47 member 1 | -1.22 | 0.010775028 |
| *FNDC4* | fibronectin type III domain containing 4 | -1.22 | 0.000891511 |
| *FCRLB* | Fc receptor like B | -1.22 | 0.001587728 |
| *ALDH1L1* | aldehyde dehydrogenase 1 family member L1 | -1.21 | 0.034105208 |
| *LRRC2* | leucine rich repeat containing 2 | -1.21 | 0.014644374 |
| *TMEM132C* | transmembrane protein 132C | -1.21 | 0.022551038 |
| *MVD* | mevalonate diphosphate decarboxylase | -1.21 | 2.49E-02 |
| *HAVCR2* | hepatitis A virus cellular receptor 2 | -1.21 | 0.01008547 |
| *ACHE* | acetylcholinesterase (Cartwright blood group) | -1.21 | 0.038909524 |
| *SNX10* | sorting nexin 10 | -1.20 | 0.008209466 |
| *C3* | complement C3 | -1.20 | 3.88E-02 |
| *CHCHD10* | coiled-coil-helix-coiled-coil-helix domain containing 10 | -1.20 | 2.4113E-06 |
| *CNTFR* | ciliary neurotrophic factor receptor | -1.19 | 2.71E-02 |
| *PFKFB1* | 6-phosphofructo-2-kinase/fructose-2,6-biphosphatase 1 | -1.19 | 0.045723788 |
| *RAMP2* | receptor activity modifying protein 2 | -1.19 | 0.015108531 |
| *ENPP3* | ectonucleotide pyrophosphatase/phosphodiesterase 3 | -1.18 | 0.047397997 |
| *CITED1* | Cbp/p300 interacting transactivator with Glu/Asp rich carboxy-terminal domain 1 | -1.18 | 0.015187005 |
| *NDRG2* | NDRG family member 2 | -1.17 | 0.014644374 |
| *DMRT2* | doublesex and mab-3 related transcription factor 2 | -1.17 | 0.043224137 |
| *FCHO1* | FCH and mu domain containing endocytic adaptor 1 | -1.16 | 0.040220188 |
| *PTPN3* | protein tyrosine phosphatase non-receptor type 3 | -1.16 | 0.018033508 |
| *SORBS1* | sorbin and SH3 domain containing 1 | -1.16 | 0.027149535 |
| *PLIN1* | perilipin 1 | -1.14 | 2.73E-02 |
| *EPHX2* | epoxide hydrolase 2 | -1.13 | 0.029536942 |

| | | | |
|---|---|---|---|
| *DTX1* | deltex E3 ubiquitin ligase 1 | -1.12 | 0.048963604 |
| *PC* | pyruvate carboxylase | -1.12 | 0.010663838 |
| *DHRS9* | dehydrogenase/reductase 9 | -1.12 | 3.62E-02 |
| *LIPE* | lipase E, hormone sensitive type | -1.12 | 0.045411139 |
| *RASD1* | ras related dexamethasone induced 1 | -1.12 | 0.028343244 |
| *ACSL1* | acyl-CoA synthetase long chain family member 1 | -1.12 | 0.040220188 |
| *F11R* | F11 receptor | -1.11 | 0.028343244 |
| *AOC3* | amine oxidase copper containing 3 | -1.11 | 4.13E-02 |
| *BMF* | Bcl2 modifying factor | -1.11 | 1.08E-02 |
| *PALMD* | palmdelphin | -1.11 | 0.018807118 |
| *TM7SF2* | transmembrane 7 superfamily member 2 | -1.10 | 3.90E-02 |
| *PPM1L* | protein phosphatase, Mg2+/Mn2+ dependent 1L | -1.09 | 0.021242961 |
| *PPARGC1B* | PPARG coactivator 1 beta | -1.09 | 0.028787624 |
| *LPCAT3* | lysophosphatidylcholine acyltransferase 3 | -1.09 | 0.035725208 |
| *REEP6* | receptor accessory protein 6 | -1.09 | 0.019949871 |
| *PDZD2* | PDZ domain containing 2 | -1.09 | 0.004263226 |
| *PKP2* | plakophilin 2 | -1.08 | 0.014644374 |
| *RETSAT* | retinol saturase | -1.08 | 0.004014571 |
| *METTL7A* | methyltransferase like 7A | -1.08 | 0.026331007 |
| *SLC25A10* | solute carrier family 25 member 10 | -1.07 | 0.000579646 |
| *AGPAT2* | 1-acylglycerol-3-phosphate O-acyltransferase 2 | -1.07 | 0.003244399 |
| *HADH* | hydroxyacyl-CoA dehydrogenase | -1.07 | 0.00929179 |
| *AR* | androgen receptor | -1.07 | 0.038909524 |
| *CLU* | clusterin | -1.03 | 0.038780413 |
| *CA5B* | carbonic anhydrase 5B | -1.03 | 0.029536942 |
| *PECR* | peroxisomal trans-2-enoyl-CoA reductase | -1.01 | 0.047397997 |
| *PLAAT3* | phospholipase A and acyltransferase 3 | -1.01 | 0.001587728 |
| *STBD1* | starch binding domain 1 | -1.00 | 0.027149535 |
| *TMEM37* | transmembrane protein 37 | -1.00 | 0.028787624 |
| *MTARC2* | mitochondrial amidoxime reducing component 2 | -1.00 | 0.000610427 |
| *SLC25A1* | solute carrier family 25 member 1 | -0.99 | 0.016546538 |
| *PNPLA2* | patatin like phospholipase domain containing 2 | -0.99 | 0.028787624 |
| *VKORC1L1* | vitamin K epoxide reductase complex subunit 1 like 1 | -0.98 | 0.007907268 |

| | | | |
|---|---|---|---|
| *SOBP* | sine oculis binding protein homolog | -0.98 | 0.005169488 |
| *CLYBL* | citramalyl-CoA lyase | -0.97 | 0.023792175 |
| *ACACA* | acetyl-CoA carboxylase alpha | -0.96 | 0.010455128 |
| *ZNF608* | zinc finger protein 608 | -0.96 | 0.038965181 |
| *WDR31* | WD repeat domain 31 | -0.96 | 0.034798199 |
| *ACLY* | ATP citrate lyase | -0.96 | 0.034105208 |
| *PDP2* | pyruvate dehyrogenase phosphatase catalytic subunit 2 | -0.96 | 0.011847894 |
| *C2CD2* | C2 calcium dependent domain containing 2 | -0.95 | 0.047397997 |
| *CS* | citrate synthase | -0.94 | 0.000827079 |
| *EPB41* | erythrocyte membrane protein band 4.1 | -0.94 | 0.000575113 |
| *LETM1* | leucine zipper and EF-hand containing transmembrane protein 1 | -0.93 | 0.028787624 |
| *RRAGD* | Ras related GTP binding D | -0.93 | 0.018978932 |
| *PLCD4* | phospholipase C delta 4 | -0.93 | 0.04761052 |
| *CPT2* | carnitine palmitoyltransferase 2 | -0.92 | 0.002342861 |
| *CPEB1* | cytoplasmic polyadenylation element binding protein 1 | -0.91 | 0.042067429 |
| *EPHX1* | epoxide hydrolase 1 | -0.90 | 0.005567697 |
| *RALGAPA2* | Ral GTPase activating protein catalytic subunit alpha 2 | -0.90 | 0.007384258 |
| *DECR1* | 2,4-dienoyl-CoA reductase 1 | -0.90 | 0.009411958 |
| *PCCA* | propionyl-CoA carboxylase subunit alpha | -0.89 | 0.028343244 |
| *SIK2* | salt inducible kinase 2 | -0.88 | 0.012177877 |
| *GPT2* | glutamic--pyruvic transaminase 2 | -0.88 | 0.004014571 |
| *ADIPOR2* | adiponectin receptor 2 | -0.87 | 0.025051449 |
| *TSHZ2* | teashirt zinc finger homeobox 2 | -0.86 | 0.037803355 |
| *ACAA2* | acetyl-CoA acyltransferase 2 | -0.86 | 0.010663838 |
| *ARHGAP42* | Rho GTPase activating protein 42 | -0.86 | 0.047645591 |
| *COX3* | cytochrome c oxidase subunit III | -0.84 | 0.001587728 |
| *FAH* | fumarylacetoacetate hydrolase | -0.84 | 0.025409004 |
| *MLYCD* | malonyl-CoA decarboxylase | -0.84 | 0.036999293 |
| *THEM6* | thioesterase superfamily member 6 | -0.84 | 0.040907997 |
| *ALDH5A1* | aldehyde dehydrogenase 5 family member A1 | -0.84 | 0.042878515 |
| *ETFDH* | electron transfer flavoprotein dehydrogenase | -0.82 | 0.043061694 |
| *ACY1* | aminoacylase 1 | -0.82 | 0.017682411 |

| | | | |
|---|---|---|---|
| *GAREM1* | GRB2 associated regulator of MAPK1 subtype 1 | -0.82 | 0.034105208 |
| *MKNK2* | MAPK interacting serine/threonine kinase 2 | -0.81 | 0.023265475 |
| *ELOVL5* | ELOVL fatty acid elongase 5 | -0.80 | 0.010455128 |
| *ISOC1* | isochorismatase domain containing 1 | -0.79 | 0.0468927 |
| *DGAT1* | diacylglycerol O-acyltransferase 1 | -0.79 | 0.017034961 |
| *HOOK2* | hook microtubule tethering protein 2 | -0.78 | 5.57E-03 |
| *DHTKD1* | dehydrogenase E1 and transketolase domain containing 1 | -0.75 | 0.028787624 |
| *ACAD10* | acyl-CoA dehydrogenase family member 10 | -0.74 | 0.026549713 |
| *APBB1IP* | amyloid beta precursor protein binding family B member 1 interacting protein | -0.74 | 0.007442582 |
| *PCDHGB7* | protocadherin gamma subfamily B, 7 | -0.72 | 0.004519361 |
| *ACO2* | aconitase 2 | -0.71 | 0.016085516 |
| *SLC25A51* | solute carrier family 25 member 51 | -0.71 | 4.31E-02 |
| *CLUH* | clustered mitochondria homolog | -0.71 | 0.04467553 |
| *TECR* | trans-2,3-enoyl-CoA reductase | -0.70 | 0.028787624 |
| *COX1* | cytochrome c oxidase subunit I | -0.70 | 0.024248613 |
| *ARHGAP32* | Rho GTPase activating protein 32 | -0.70 | 0.021836614 |
| *ERAP2* | endoplasmic reticulum aminopeptidase 2 | 2.37 | 8.08937E-11 |
| *CLEC14A* | C-type lectin domain containing 14A | 1.45 | 0.000695096 |
| *OLR1* | oxidized low density lipoprotein receptor 1 | 1.33 | 0.014644374 |
| *NOG* | noggin | 1.30 | 7.85E-03 |
| *PTGS2* | prostaglandin-endoperoxide synthase 2 | 1.29 | 0.003089649 |
| *PEAR1* | platelet endothelial aggregation receptor 1 | 1.25 | 0.000238258 |
| *CARD9* | caspase recruitment domain family member 9 | 1.24 | 0.024780772 |
| *KRT34* | keratin 34 | 1.24 | 2.83E-02 |
| *TBX1* | T-box transcription factor 1 | 1.23 | 3.56E-02 |
| *FLT1* | fms related receptor tyrosine kinase 1 | 1.20 | 0.029536942 |
| *CADPS* | calcium dependent secretion activator | 1.18 | 1.65E-02 |
| *SCUBE3* | signal peptide, CUB domain and EGF like domain containing 3 | 1.18 | 4.92E-02 |
| *ISYNA1* | inositol-3-phosphate synthase 1 | 1.16 | 1.03E-02 |
| *PCSK9* | proprotein convertase subtilisin/kexin type 9 | 1.09 | 2.67E-03 |

Supplementary Table 5 DEGs from comparison of inactive beige adipocytes carrying *FTO* obesity-risk vs risk-free genotypes; n=4 of each genotype

| Symbol | Gene Name | Log2 Fold Change | Adj. P value |
|---|---|---|---|
| *CLIC6* | chloride intracellular channel 6 | -1.50 | 8.59E-03 |
| *PLXNC1* | plexin C1 | -1.44 | 1.92E-02 |
| *UCP2* | uncoupling protein 2 | -1.41 | 5.30E-03 |
| *CNGA3* | cyclic nucleotide gated channel subunit alpha 3 | -1.39 | 3.11E-02 |
| *STON2* | stonin 2 | -1.36 | 2.74E-02 |
| *ZNF385A* | zinc finger protein 385A | -1.36 | 1.43E-04 |
| *SIM1* | SIM bHLH transcription factor 1 | -1.33 | 9.37E-03 |
| *TXLNB* | taxilin beta | -1.32 | 4.36E-02 |
| *PNLIPRP3* | pancreatic lipase related protein 3 | -1.31 | 4.51E-02 |
| *NA* | NA | -1.02 | 4.63E-02 |
| *BMF* | Bcl2 modifying factor | -0.89 | 0.031123633 |
| *ERAP2* | endoplasmic reticulum aminopeptidase 2 | 3.22 | 2.22E-42 |
| *NOG* | noggin | 1.87 | 2.73192E-05 |
| *SEMA5B* | semaphorin 5B | 1.81 | 1.43E-04 |
| *FLT1* | fms related receptor tyrosine kinase 1 | 1.78 | 1.87E-04 |
| *GDF7* | growth differentiation factor 7 | 1.59 | 4.33E-03 |
| *CLEC14A* | C-type lectin domain containing 14A | 1.54 | 1.03E-04 |
| *CARD9* | caspase recruitment domain family member 9 | 1.50 | 7.45E-03 |
| *PEAR1* | platelet endothelial aggregation receptor 1 | 1.49 | 0.000143015 |
| *CPA4* | carboxypeptidase A4 | 1.49 | 1.71E-03 |
| *LGR5* | leucine rich repeat containing G protein-coupled receptor 5 | 1.48 | 0.007978884 |
| *DIO2* | iodothyronine deiodinase 2 | 1.42 | 2.41E-02 |
| *TRPC6* | transient receptor potential cation channel subfamily C member 6 | 1.40 | 0.027381653 |
| *SCUBE3* | signal peptide, CUB domain and EGF like domain containing 3 | 1.37 | 0.033390076 |
| *CDH8* | cadherin 8 | 1.35 | 0.036257298 |
| *ADRA2C* | adrenoceptor alpha 2C | 1.34 | 0.040177627 |
| *ANKRD29* | ankyrin repeat domain 29 | 1.33 | 1.26E-04 |
| *TSPAN8* | tetraspanin 8 | 1.30 | 3.45E-02 |
| *TINAGL1* | tubulointerstitial nephritis antigen like 1 | 1.30 | 0.049441547 |
| *PADI2* | peptidyl arginine deiminase 2 | 1.29 | 4.88E-02 |

| | | | |
|---|---|---|---|
| *MET* | MET proto-oncogene, receptor tyrosine kinase | 1.26 | 0.042189152 |
| *TGM1* | transglutaminase 1 | 1.22 | 4.94E-02 |
| *ISYNA1* | inositol-3-phosphate synthase 1 | 1.18 | 0.036257298 |
| *MGARP* | mitochondria localized glutamic acid rich protein | 1.17 | 4.10E-05 |
| *MIR210HG* | MIR210 host gene | 1.17 | 0.045120682 |
| *BDNF* | brain derived neurotrophic factor | 1.11 | 0.046302206 |
| *THSD4* | thrombospondin type 1 domain containing 4 | 1.11 | 0.033426202 |
| *FHL1* | four and a half LIM domains 1 | 1.11 | 2.07E-02 |
| *XRRA1* | X-ray radiation resistance associated 1 | 1.07 | 9.04E-07 |
| *MYH3* | myosin heavy chain 3 | 1.06 | 0.0416563 |
| *EMP1* | epithelial membrane protein 1 | 1.04 | 0.003956507 |
| *GDNF* | glial cell derived neurotrophic factor | 1.02 | 0.034848749 |
| *ULK4* | unc-51 like kinase 4 | 1.00 | 0.046302206 |
| *SLC1A1* | solute carrier family 1 member 1 | 0.98 | 0.003278171 |
| *FANK1* | fibronectin type III and ankyrin repeat domains 1 | 0.98 | 3.14E-02 |
| *ITGB3BP* | integrin subunit beta 3 binding protein | 0.90 | 1.77E-02 |
| *EXOG* | exo/endonuclease G | 0.90 | 0.046302206 |
| *MAP4K3-DT* | MAP4K3 divergent transcript | 0.89 | 0.043575692 |
| *ANGPT1* | angiopoietin 1 | 0.88 | 2.31E-02 |
| *ENDOD1* | endonuclease domain containing 1 | 0.88 | 8.40E-05 |
| *DNAJC6* | DnaJ heat shock protein family (Hsp40) member C6 | 0.85 | 0.036257298 |
| *LOX* | lysyl oxidase | 0.83 | 0.010401742 |
| *PRRX2* | paired related homeobox 2 | 0.79 | 0.044614199 |
| *TRNP1* | TMF1 regulated nuclear protein 1 | 0.77 | 0.0030316 |
| *PGM1* | phosphoglucomutase 1 | 0.77 | 0.040358227 |
| *ALPK2* | alpha kinase 2 | 0.75 | 0.043575692 |
| *STX6* | syntaxin 6 | 0.75 | 2.31E-02 |

Supplementary Table 6 DEGs from comparison of active beige and white adipocytes carrying FTO risk-free genotype; n=4 of each genotype

| **Symbol** | **Gene Name** | **Log2 Fold Change** | **Adj. P value** |
|---|---|---|---|
| *PCK1* | phosphoenolpyruvate carboxykinase 1 | 3.19 | 6.26E-23 |
| *CD96* | CD96 molecule | 2.97 | 2.95E-14 |
| *BBOX1* | gamma-butyrobetaine hydroxylase 1 | 2.76 | 1.31E-15 |
| *KRT79* | keratin 79 | 2.75 | 4.72E-12 |
| *TMEM132C* | transmembrane protein 132C | 2.70 | 1.32E-14 |
| *PM20D1* | peptidase M20 domain containing 1 | 2.56 | 1.17E-09 |
| *SLC22A12* | solute carrier family 22 member 12 | 2.53 | 5.12E-10 |
| *FCN1* | ficolin 1 | 2.50 | 1.42E-09 |
| *ADAMTS18* | ADAM metallopeptidase with thrombospondin type 1 motif 18 | 2.42 | 2.94E-09 |
| *KCNA1* | potassium voltage-gated channel subfamily A member 1 | 2.37 | 1.36E-12 |
| *SCN4A* | sodium voltage-gated channel alpha subunit 4 | 2.31 | 4.35826E-09 |
| *MGAT3* | beta-1,4-mannosyl-glycoprotein 4-beta-N-acetylglucosaminyltransferase | 2.26 | 1.21E-11 |
| *NA* | NA | 2.26 | 4.05578E-08 |
| *TSHR* | thyroid stimulating hormone receptor | 2.24 | 2.55E-07 |
| *ABCG1* | ATP binding cassette subfamily G member 1 | 2.10 | 2.89E-16 |
| *AP3B2* | adaptor related protein complex 3 subunit beta 2 | 2.08 | 1.36E-12 |
| *CPA4* | carboxypeptidase A4 | 2.02 | 4.71E-07 |
| *PDE1B* | phosphodiesterase 1B | 2.02 | 2.92E-08 |
| *CASZ1* | castor zinc finger 1 | 2.01 | 8.87546E-07 |
| *CD52* | CD52 molecule | 2.00 | 6.53E-06 |
| *LHCGR* | luteinizing hormone/choriogonadotropin receptor | 1.94 | 6.17593E-07 |
| *KCNK12* | potassium two pore domain channel subfamily K member 12 | 1.94 | 1.91E-05 |
| *ANO5* | anoctamin 5 | 1.89 | 1.45464E-05 |
| *PDZD2* | PDZ domain containing 2 | 1.89 | 1.7319E-06 |
| *STX11* | syntaxin 11 | 1.88 | 9.57852E-11 |
| *PPP1R1A* | protein phosphatase 1 regulatory inhibitor subunit 1A | 1.85 | 2.77003E-06 |

| | | | |
|---|---|---|---|
| *LINC02554* | long intergenic non-protein coding RNA 2554 | 1.84 | 6.88E-05 |
| *CA3* | carbonic anhydrase 3 | 1.84 | 6.87E-05 |
| *PLPPR4* | phospholipid phosphatase related 4 | 1.84 | 2.98685E-06 |
| *LPL* | lipoprotein lipase | 1.83 | 1.88E-06 |
| *SLC22A3* | solute carrier family 22 member 3 | 1.82 | 1.71964E-07 |
| *CKMT1B* | creatine kinase, mitochondrial 1B | 1.81 | 2.30E-05 |
| *CYP4F12* | cytochrome P450 family 4 subfamily F member 12 | 1.81 | 3.61872E-07 |
| *ICA1* | islet cell autoantigen 1 | 1.79 | 2.50E-07 |
| *TRHDE-AS1* | TRHDE antisense RNA 1 | 1.78 | 1.43612E-09 |
| *CDH6* | cadherin 6 | 1.78 | 7.29733E-05 |
| *AQP3* | aquaporin 3 (Gill blood group) | 1.73 | 9.46377E-05 |
| *SORL1* | sortilin related receptor 1 | 1.71 | 1.83E-04 |
| *SCAMP5* | secretory carrier membrane protein 5 | 1.69 | 1.89E-05 |
| *FNDC1* | fibronectin type III domain containing 1 | 1.67 | 0.000236687 |
| *FABP4* | fatty acid binding protein 4 | 1.67 | 8.87546E-07 |
| *PLIN5* | perilipin 5 | 1.67 | 3.67898E-05 |
| *PLEKHG6* | pleckstrin homology and RhoGEF domain containing G6 | 1.66 | 9.35461E-06 |
| *CITED1* | Cbp/p300 interacting transactivator with Glu/Asp rich carboxy-terminal domain 1 | 1.64 | 2.16821E-05 |
| *CADM3* | cell adhesion molecule 3 | 1.64 | 6.94E-04 |
| *PALMD* | palmdelphin | 1.64 | 6.88E-05 |
| *EPB41L4B* | erythrocyte membrane protein band 4.1 like 4B | 1.64 | 6.39081E-05 |
| *RBM24* | RNA binding motif protein 24 | 1.63 | 8.57318E-06 |
| *FCHO1* | FCH and mu domain containing endocytic adaptor 1 | 1.60 | 7.13E-05 |
| *RGL3* | ral guanine nucleotide dissociation stimulator like 3 | 1.59 | 1.00E-04 |
| *UCP1* | uncoupling protein 1 | 1.59 | 0.001326054 |
| *IL1B* | interleukin 1 beta | 1.58 | 0.00058207 |
| *TRPM8* | transient receptor potential cation channel subfamily M member 8 | 1.56 | 0.001326054 |
| *TSPAN15* | tetraspanin 15 | 1.54 | 0.000344402 |
| *LRRC15* | leucine rich repeat containing 15 | 1.53 | 0.002078686 |
| *C4orf19* | chromosome 4 open reading frame 19 | 1.51 | 0.0021136 |

| | | | |
|---|---|---|---|
| *ADAM12* | ADAM metallopeptidase domain 12 | 1.51 | 4.52E-05 |
| *THBD* | thrombomodulin | 1.50 | 0.001543075 |
| *GK* | glycerol kinase | 1.50 | 1.7319E-06 |
| *ITGA8* | integrin subunit alpha 8 | 1.50 | 0.001901231 |
| *CLDN2* | claudin 2 | 1.50 | 0.00355501 |
| *LMO2* | LIM domain only 2 | 1.50 | 0.000755093 |
| *TMEM130* | transmembrane protein 130 | 1.50 | 0.003659612 |
| *ADIPOQ* | adiponectin, C1Q and collagen domain containing | 1.49 | 0.002078686 |
| *CTXN1* | cortexin 1 | 1.49 | 8.57318E-06 |
| *C1QTNF3* | C1q and TNF related 3 | 1.49 | 0.001193504 |
| *ECSCR* | endothelial cell surface expressed chemotaxis and apoptosis regulator | 1.48 | 0.004085113 |
| *MEST* | mesoderm specific transcript | 1.48 | 8.87116E-06 |
| *PCSK1* | proprotein convertase subtilisin/kexin type 1 | 1.48 | 8.09148E-05 |
| *RASD1* | ras related dexamethasone induced 1 | 1.48 | 1.26058E-10 |
| *ZNF365* | zinc finger protein 365 | 1.47 | 0.001730505 |
| *RTKN2* | rhotekin 2 | 1.45 | 0.000528277 |
| *RIMS4* | regulating synaptic membrane exocytosis 4 | 1.45 | 0.00207578 |
| *SCN9A* | sodium voltage-gated channel alpha subunit 9 | 1.45 | 0.003049696 |
| *GCOM1* | GRINL1A complex locus 1 | 1.44 | 1.57455E-05 |
| *MOGAT1* | monoacylglycerol O-acyltransferase 1 | 1.42 | 0.004013574 |
| *CDHR1* | cadherin related family member 1 | 1.41 | 1.59E-04 |
| *GRAP* | GRB2 related adaptor protein | 1.40 | 0.003659612 |
| *TMEM200C* | transmembrane protein 200C | 1.39 | 0.007634929 |
| *CIART* | circadian associated repressor of transcription | 1.39 | 3.06911E-05 |
| *PCDH19* | protocadherin 19 | 1.39 | 3.49E-03 |
| *IRF8* | interferon regulatory factor 8 | 1.38 | 0.008630167 |
| *LAMP3* | lysosomal associated membrane protein 3 | 1.37 | 8.86E-03 |
| *RXRG* | retinoid X receptor gamma | 1.36 | 0.001397029 |
| *SLC19A3* | solute carrier family 19 member 3 | 1.35 | 0.000470289 |
| *ADAMTS14* | ADAM metallopeptidase with thrombospondin type 1 motif 14 | 1.35 | 0.008943036 |
| *CYS1* | cystin 1 | 1.34 | 0.000667895 |

| | | | |
|---|---|---|---|
| *ACKR1* | atypical chemokine receptor 1 (Duffy blood group) | 1.34 | 0.012237669 |
| *LOC100507560* | uncharacterized LOC100507560 | 1.34 | 0.011658454 |
| *KLHDC7A* | kelch domain containing 7A | 1.34 | 0.015981012 |
| *APLP1* | amyloid beta precursor like protein 1 | 1.33 | 6.23237E-07 |
| *RGS7BP* | regulator of G protein signaling 7 binding protein | 1.33 | 0.004697797 |
| *TENT5C* | terminal nucleotidyltransferase 5C | 1.33 | 8.35E-03 |
| *CKMT1A* | creatine kinase, mitochondrial 1A | 1.32 | 0.017436439 |
| *CPA2* | carboxypeptidase A2 | 1.32 | 0.017127307 |
| *SRCIN1* | SRC kinase signaling inhibitor 1 | 1.32 | 0.005985029 |
| *HPSE2* | heparanase 2 (inactive) | 1.32 | 1.32E-02 |
| *MGAT4A* | alpha-1,3-mannosyl-glycoprotein 4-beta-N-acetylglucosaminyltransferase A | 1.31 | 0.005533282 |
| *EPHB2* | EPH receptor B2 | 1.30 | 0.002375381 |
| *CCN3* | cellular communication network factor 3 | 1.29 | 0.004008059 |
| *LBP* | lipopolysaccharide binding protein | 1.29 | 0.006272526 |
| *KCNK15* | potassium two pore domain channel subfamily K member 15 | 1.28 | 1.09E-02 |
| *ASS1* | argininosuccinate synthase 1 | 1.28 | 5.00E-12 |
| *FAM151A* | family with sequence similarity 151 member A | 1.27 | 0.025035351 |
| *EGFLAM* | EGF like, fibronectin type III and laminin G domains | 1.27 | 2.46E-03 |
| *SLC7A10* | solute carrier family 7 member 10 | 1.26 | 0.025674126 |
| *PTPN3* | protein tyrosine phosphatase non-receptor type 3 | 1.26 | 0.000752433 |
| *TRHDE* | thyrotropin releasing hormone degrading enzyme | 1.25 | 0.000632671 |
| *C14orf180* | chromosome 14 open reading frame 180 | 1.24 | 0.017017843 |
| *LINC01347* | long intergenic non-protein coding RNA 1347 | 1.23 | 0.03104806 |
| *DLX3* | distal-less homeobox 3 | 1.23 | 0.02712039 |
| *SYNGR2* | synaptogyrin 2 | 1.23 | 0.000643619 |
| *MPP7* | membrane palmitoylated protein 7 | 1.23 | 0.003245163 |
| *TMEM131L* | transmembrane 131 like | 1.23 | 1.87822E-06 |
| *HAS3* | hyaluronan synthase 3 | 1.22 | 0.024592675 |
| *CXADRP3* | CXADR pseudogene 3 | 1.22 | 0.037446998 |
| *CKMT2* | creatine kinase, mitochondrial 2 | 1.21 | 0.007824085 |

| | | | |
|---|---|---|---|
| *LOC100506253* | uncharacterized LOC100506253 | 1.21 | 0.040464703 |
| *ANOS1* | anosmin 1 | 1.21 | 0.040882609 |
| *CDH13* | cadherin 13 | 1.20 | 0.000146848 |
| *LUCAT1* | lung cancer associated transcript 1 | 1.20 | 0.030443356 |
| *PDE1C* | phosphodiesterase 1C | 1.20 | 0.022991064 |
| *L3MBTL4* | L3MBTL histone methyl-lysine binding protein 4 | 1.19 | 0.044001175 |
| *CIDEA* | cell death inducing DFFA like effector a | 1.19 | 0.047144109 |
| *GSG1* | germ cell associated 1 | 1.19 | 0.022143679 |
| *GTF2IP1* | general transcription factor IIi pseudogene 1 | 1.19 | 0.043541916 |
| *PLPP2* | phospholipid phosphatase 2 | 1.19 | 0.013450146 |
| *SAMD10* | sterile alpha motif domain containing 10 | 1.18 | 0.001888824 |
| *SH2D3C* | SH2 domain containing 3C | 1.18 | 0.03156525 |
| *FGF13* | fibroblast growth factor 13 | 1.17 | 0.027468375 |
| *ADGRB2* | adhesion G protein-coupled receptor B2 | 1.17 | 0.006143259 |
| *PPM1H* | protein phosphatase, Mg2+/Mn2+ dependent 1H | 1.16 | 0.003387417 |
| *DPYSL3* | dihydropyrimidinase like 3 | 1.16 | 8.87546E-07 |
| *HCAR3* | hydroxycarboxylic acid receptor 3 | 1.16 | 0.04886069 |
| *PLIN4* | perilipin 4 | 1.16 | 0.04518701 |
| *KRT14* | keratin 14 | 1.15 | 0.012986163 |
| *SLC24A4* | solute carrier family 24 member 4 | 1.15 | 0.048615779 |
| *SELPLG* | selectin P ligand | 1.15 | 0.0064944 |
| *FLVCR2* | FLVCR heme transporter 2 | 1.13 | 0.02712039 |
| *ATP1A3* | ATPase Na+/K+ transporting subunit alpha 3 | 1.13 | 0.04584214 |
| *PTGDS* | prostaglandin D2 synthase | 1.13 | 1.41466E-10 |
| *MMD* | monocyte to macrophage differentiation associated | 1.13 | 0.022316681 |
| *SAMD14* | sterile alpha motif domain containing 14 | 1.12 | 0.002348835 |
| *WARS1* | tryptophanyl-tRNA synthetase 1 | 1.12 | 0.001051673 |
| *MAP3K7CL* | MAP3K7 C-terminal like | 1.12 | 0.004977903 |
| *RGS2* | regulator of G protein signaling 2 | 1.11 | 8.22041E-05 |
| *ABCA3* | ATP binding cassette subfamily A member 3 | 1.11 | 0.022052525 |
| *FBXO27* | F-box protein 27 | 1.10 | 1.46454E-05 |
| *KIAA1549* | KIAA1549 | 1.09 | 0.007517754 |

| | | | |
|---|---|---|---|
| *FYB1* | FYN binding protein 1 | 1.09 | 0.007768415 |
| *HSD17B6* | hydroxysteroid 17-beta dehydrogenase 6 | 1.09 | 0.03104806 |
| *IFI30* | IFI30 lysosomal thiol reductase | 1.08 | 0.007553361 |
| *CNIH3* | cornichon family AMPA receptor auxiliary protein 3 | 1.08 | 0.026170681 |
| *ITIH5* | inter-alpha-trypsin inhibitor heavy chain 5 | 1.07 | 0.032877012 |
| *TENM4* | teneurin transmembrane protein 4 | 1.07 | 0.006670827 |
| *JAG2* | jagged canonical Notch ligand 2 | 1.06 | 0.018880928 |
| *COL8A2* | collagen type VIII alpha 2 chain | 1.06 | 0.000931703 |
| *RUBCNL* | rubicon like autophagy enhancer | 1.04 | 0.018550658 |
| *EPB41* | erythrocyte membrane protein band 4.1 | 1.03 | 0.004416376 |
| *RALGPS1* | Ral GEF with PH domain and SH3 binding motif 1 | 1.02 | 0.011726318 |
| *GPC4* | glypican 4 | 1.02 | 0.006224509 |
| *GPR153* | G protein-coupled receptor 153 | 1.02 | 9.22297E-08 |
| *STAT4* | signal transducer and activator of transcription 4 | 1.01 | 0.004333457 |
| *NHS* | NHS actin remodeling regulator | 1.00 | 0.0254592 |
| *GPC1* | glypican 1 | 0.99 | 1.74E-04 |
| *KIAA1549L* | KIAA1549 like | 0.99 | 0.001193359 |
| *NOCT* | nocturnin | 0.99 | 0.004333457 |
| *SEL1L3* | SEL1L family member 3 | 0.98 | 0.013772341 |
| *PER3* | period circadian regulator 3 | 0.98 | 0.019361196 |
| *CA5B* | carbonic anhydrase 5B | 0.97 | 0.047981213 |
| *GXYLT2* | glucoside xylosyltransferase 2 | 0.97 | 1.89E-02 |
| *FHL1* | four and a half LIM domains 1 | 0.97 | 0.000341183 |
| *ARNTL2* | aryl hydrocarbon receptor nuclear translocator like 2 | 0.96 | 0.020946553 |
| *FNIP2* | folliculin interacting protein 2 | 0.95 | 0.027764502 |
| *BANK1* | B cell scaffold protein with ankyrin repeats 1 | 0.94 | 0.023973182 |
| *CSAD* | cysteine sulfinic acid decarboxylase | 0.94 | 0.039509392 |
| *CHN2* | chimerin 2 | 0.94 | 0.027468375 |
| *GUCY1A2* | guanylate cyclase 1 soluble subunit alpha 2 | 0.94 | 0.005669642 |
| *LACC1* | laccase domain containing 1 | 0.94 | 2.62E-02 |
| *PHLDA1* | pleckstrin homology like domain family A member 1 | 0.93 | 0.023973182 |
| *SLC19A1* | solute carrier family 19 member 1 | 0.93 | 0.00114516 |

| | | | |
|---|---|---|---|
| *SEMA6D* | semaphorin 6D | 0.92 | 0.006948144 |
| *IL20RA* | interleukin 20 receptor subunit alpha | 0.91 | 1.87E-02 |
| *ACAA2* | acetyl-CoA acyltransferase 2 | 0.90 | 4.73E-04 |
| *MYO1B* | myosin IB | 0.89 | 0.002057379 |
| *MYEF2* | myelin expression factor 2 | 0.89 | 3.22E-02 |
| *PER2* | period circadian regulator 2 | 0.89 | 3.93E-02 |
| *GLIS2* | GLIS family zinc finger 2 | 0.88 | 2.62E-05 |
| *TRIM6* | tripartite motif containing 6 | 0.88 | 2.95E-02 |
| *BHLHE41* | basic helix-loop-helix family member e41 | 0.87 | 4.58E-02 |
| *QRICH2* | glutamine rich 2 | 0.87 | 2.53E-02 |
| *FAM227A* | family with sequence similarity 227 member A | 0.86 | 2.74E-02 |
| *PIK3R2* | phosphoinositide-3-kinase regulatory subunit 2 | 0.85 | 8.82E-05 |
| *AKNA* | AT-hook transcription factor | 0.84 | 9.90E-03 |
| *NR1D2* | nuclear receptor subfamily 1 group D member 2 | 0.84 | 5.28E-04 |
| *NUDT14* | nudix hydrolase 14 | 0.84 | 0.027391618 |
| *DTX4* | deltex E3 ubiquitin ligase 4 | 0.84 | 3.50E-02 |
| *MOCOS* | molybdenum cofactor sulfurase | 0.84 | 0.023862149 |
| *ASPHD2* | aspartate beta-hydroxylase domain containing 2 | 0.83 | 0.019396186 |
| *AK4* | adenylate kinase 4 | 0.83 | 1.56E-02 |
| *NCS1* | neuronal calcium sensor 1 | 0.83 | 6.95E-06 |
| *SLC6A9* | solute carrier family 6 member 9 | 0.82 | 3.08E-02 |
| *ITGA4* | integrin subunit alpha 4 | 0.82 | 3.48E-02 |
| *PRRX2* | paired related homeobox 2 | 0.82 | 3.46E-02 |
| *PPIP5K1* | diphosphoinositol pentakisphosphate kinase 1 | 0.82 | 0.024812449 |
| *UHRF1BP1* | UHRF1 binding protein 1 | 0.81 | 4.89E-02 |
| *DOK5* | docking protein 5 | 0.81 | 4.37E-04 |
| *LIMS1* | LIM zinc finger domain containing 1 | 0.80 | 0.000707977 |
| *APOL6* | apolipoprotein L6 | 0.79 | 4.63E-02 |
| *GTPBP2* | GTP binding protein 2 | 0.79 | 0.048615779 |
| *CCSAP* | centriole, cilia and spindle associated protein | 0.79 | 0.032119771 |
| *THY1* | Thy-1 cell surface antigen | 0.78 | 0.018245584 |
| *RGS3* | regulator of G protein signaling 3 | 0.77 | 0.015624435 |
| *SHB* | SH2 domain containing adaptor protein B | 0.77 | 0.048563756 |

| | | | |
|---|---|---|---|
| *PWWP2B* | PWWP domain containing 2B | 0.77 | 0.008943036 |
| *EGR2* | early growth response 2 | 0.76 | 2.80E-02 |
| *MLYCD* | malonyl-CoA decarboxylase | 0.76 | 1.89E-02 |
| *MFSD9* | major facilitator superfamily domain containing 9 | 0.76 | 0.04886069 |
| *COL27A1* | collagen type XXVII alpha 1 chain | 0.76 | 0.015624435 |
| *ATP8B1* | ATPase phospholipid transporting 8B1 | 0.75 | 0.017260247 |
| *TET3* | tet methylcytosine dioxygenase 3 | 0.75 | 4.71E-02 |
| *MET* | MET proto-oncogene, receptor tyrosine kinase | 0.74 | 0.038557109 |
| *XPOT* | exportin for tRNA | 0.74 | 0.037268889 |
| *RBM38* | RNA binding motif protein 38 | 0.73 | 0.035043311 |
| *ALG9* | ALG9 alpha-1,2-mannosyltransferase | 0.73 | 4.84E-02 |
| *SDC3* | syndecan 3 | 0.72 | 0.011536489 |
| *CACNB1* | calcium voltage-gated channel auxiliary subunit beta 1 | 0.72 | 0.011540997 |
| *ZBTB16* | zinc finger and BTB domain containing 16 | -4.38 | 1.14018E-55 |
| *TIMP4* | TIMP metallopeptidase inhibitor 4 | -2.34 | 3.23663E-08 |
| *FKBP5* | FKBP prolyl isomerase 5 | -2.24 | 2.27E-08 |
| *SAA1* | serum amyloid A1 | -2.22 | 3.3072E-07 |
| *PER1* | period circadian regulator 1 | -2.18 | 4.99E-15 |
| *MMP28* | matrix metallopeptidase 28 | -2.11 | 8.02647E-09 |
| *INHBB* | inhibin subunit beta B | -2.04 | 9.22E-08 |
| *GABRA5* | gamma-aminobutyric acid type A receptor subunit alpha5 | -1.97 | 8.87E-06 |
| *ANGPTL8* | angiopoietin like 8 | -1.89 | 3.71154E-05 |
| *APCDD1* | APC down-regulated 1 | -1.86 | 2.3894E-12 |
| *ATP1A2* | ATPase Na+/K+ transporting subunit alpha 2 | -1.81 | 0.000106102 |
| *DPT* | dermatopontin | -1.78 | 8.57318E-06 |
| *TMPRSS5* | transmembrane serine protease 5 | -1.77 | 8.57318E-06 |
| *ABCC2* | ATP binding cassette subfamily C member 2 | -1.76 | 1.74E-04 |
| *PTGS1* | prostaglandin-endoperoxide synthase 1 | -1.73 | 4.79878E-13 |
| *CRLF1* | cytokine receptor like factor 1 | -1.70 | 3.92E-07 |
| *LRP1B* | LDL receptor related protein 1B | -1.70 | 0.000341183 |
| *MAOA* | monoamine oxidase A | -1.69 | 0.000174045 |
| *CRISPLD2* | cysteine rich secretory protein LCCL domain containing 2 | -1.64 | 2.38623E-05 |

| | | | |
|---|---|---|---|
| *SLC16A12* | solute carrier family 16 member 12 | -1.63 | 0.00091505 |
| *TENT5B* | terminal nucleotidyltransferase 5B | -1.61 | 6.86903E-06 |
| *CPM* | carboxypeptidase M | -1.60 | 3.67898E-05 |
| *LMO3* | LIM domain only 3 | -1.58 | 0.000144809 |
| *PILRA* | paired immunoglobin like type 2 receptor alpha | -1.57 | 0.001492983 |
| *ACKR2* | atypical chemokine receptor 2 | -1.57 | 1.27998E-05 |
| *GALNT15* | polypeptide N-acetylgalactosaminyltransferase 15 | -1.55 | 0.002078686 |
| *NEGR1* | neuronal growth regulator 1 | -1.54 | 1.71964E-07 |
| *AVPR1A* | arginine vasopressin receptor 1A | -1.51 | 0.003277674 |
| *SYN2* | synapsin II | -1.49 | 0.003741136 |
| *NKD2* | NKD inhibitor of WNT signaling pathway 2 | -1.48 | 0.000751398 |
| *TMC2* | transmembrane channel like 2 | -1.47 | 0.004416376 |
| *STC1* | stanniocalcin 1 | -1.46 | 0.004329215 |
| *TF* | transferrin | -1.45 | 5.65E-04 |
| *METTL7A* | methyltransferase like 7A | -1.44 | 0.001193359 |
| *HIF3A* | hypoxia inducible factor 3 subunit alpha | -1.41 | 0.004567973 |
| *NA* | NA | -1.41 | 0.007455214 |
| *SERPINA3* | serpin family A member 3 | -1.40 | 0.000242257 |
| *C6* | complement C6 | -1.40 | 0.006601815 |
| *TSC22D3* | TSC22 domain family member 3 | -1.40 | 1.42408E-09 |
| *GLYAT* | glycine-N-acyltransferase | -1.39 | 0.009462246 |
| *RASL11A* | RAS like family 11 member A | -1.38 | 1.19E-03 |
| *GSTA1* | glutathione S-transferase alpha 1 | -1.36 | 1.16E-02 |
| *MGP* | matrix Gla protein | -1.34 | 0.014510646 |
| *HPD* | 4-hydroxyphenylpyruvate dioxygenase | -1.34 | 0.014492172 |
| *CLCN1* | chloride voltage-gated channel 1 | -1.32 | 0.018198762 |
| *CYP8B1* | cytochrome P450 family 8 subfamily B member 1 | -1.31 | 0.00622792 |
| *PKD2L1* | polycystin 2 like 1, transient receptor potential cation channel | -1.31 | 0.018346649 |
| *ACSL6* | acyl-CoA synthetase long chain family member 6 | -1.30 | 0.02049558 |
| *IP6K3* | inositol hexakisphosphate kinase 3 | -1.30 | 0.016867757 |
| *HSPA6* | heat shock protein family A (Hsp70) member 6 | -1.28 | 0.018861605 |
| *ISM1* | isthmin 1 | -1.28 | 0.022052525 |

| | | | |
|---|---|---|---|
| *MAP2K6* | mitogen-activated protein kinase kinase 6 | -1.28 | 0.024547624 |
| *HS3ST2* | heparan sulfate-glucosamine 3-sulfotransferase 2 | -1.27 | 0.024547624 |
| *AACS* | acetoacetyl-CoA synthetase | -1.26 | 4.33E-03 |
| *ARHGEF16* | Rho guanine nucleotide exchange factor 16 | -1.25 | 0.026445094 |
| *CST3* | cystatin C | -1.24 | 0.001901231 |
| *ANGPTL1* | angiopoietin like 1 | -1.23 | 0.018936049 |
| *ALS2CL* | ALS2 C-terminal like | -1.23 | 0.005588454 |
| *PLCH1* | phospholipase C eta 1 | -1.23 | 0.03625776 |
| *GLUL* | glutamate-ammonia ligase | -1.22 | 0.007824085 |
| *DNASE1L3* | deoxyribonuclease 1 like 3 | -1.21 | 0.020704394 |
| *RXFP1* | relaxin family peptide receptor 1 | -1.20 | 0.044001175 |
| *FMO2* | flavin containing dimethylaniline monoxygenase 2 | -1.20 | 0.034567002 |
| *MT1X* | metallothionein 1X | -1.19 | 0.013807091 |
| *SLC2A13* | solute carrier family 2 member 13 | -1.18 | 0.034752967 |
| *IGFBP2* | insulin like growth factor binding protein 2 | -1.17 | 0.04674927 |
| *TRNP1* | TMF1 regulated nuclear protein 1 | -1.16 | 3.02E-07 |
| *KCNK5* | potassium two pore domain channel subfamily K member 5 | -1.14 | 0.047144109 |
| *CYP4B1* | cytochrome P450 family 4 subfamily B member 1 | -1.14 | 0.029373766 |
| *CUTC* | cutC copper transporter | -1.14 | 5.36E-05 |
| *WNT5A* | Wnt family member 5A | -1.12 | 0.03478741 |
| *CTHRC1* | collagen triple helix repeat containing 1 | -1.10 | 0.03070775 |
| *ELANE* | elastase, neutrophil expressed | -1.10 | 0.019245127 |
| *C1QTNF7* | C1q and TNF related 7 | -1.10 | 0.015274773 |
| *ABLIM3* | actin binding LIM protein family member 3 | -1.09 | 0.046759343 |
| *CORO6* | coronin 6 | -1.03 | 0.018861605 |
| *DUSP4* | dual specificity phosphatase 4 | -1.01 | 3.14E-03 |
| *SMOC2* | SPARC related modular calcium binding 2 | -1.01 | 5.95E-03 |
| *KLF9* | Kruppel like factor 9 | -1.00 | 0.005864639 |
| *SPARCL1* | SPARC like 1 | -1.00 | 0.032877012 |
| *HRCT1* | histidine rich carboxyl terminus 1 | -1.00 | 0.016531751 |
| *RGMA* | repulsive guidance molecule BMP co-receptor a | -0.99 | 0.013679865 |

| | | | |
|---|---|---|---|
| *GSN* | gelsolin | -0.98 | 0.037068588 |
| *GPSM2* | G protein signaling modulator 2 | -0.97 | 0.02150243 |
| *ZNF395* | zinc finger protein 395 | -0.96 | 8.95E-08 |
| *SMARCD2* | SWI/SNF related, matrix associated, actin dependent regulator of chromatin, subfamily d, member 2 | -0.96 | 0.000215617 |
| *TMEM59L* | transmembrane protein 59 like | -0.96 | 0.012986163 |
| *HLX* | H2.0 like homeobox | -0.94 | 1.74E-04 |
| *TWF2* | twinfilin actin binding protein 2 | -0.89 | 4.48E-03 |
| *ERRFI1* | ERBB receptor feedback inhibitor 1 | -0.89 | 0.044614151 |
| *DUSP1* | dual specificity phosphatase 1 | -0.85 | 0.002303644 |
| *HMGN5* | high mobility group nucleosome binding domain 5 | -0.82 | 0.025975231 |
| *ADAMTS1* | ADAM metallopeptidase with thrombospondin type 1 motif 1 | -0.77 | 1.03E-02 |
| *FAM43A* | family with sequence similarity 43 member A | -0.73 | 1.82E-02 |
| *GAS1* | growth arrest specific 1 | -0.73 | 0.039012627 |

Supplementary Table 7 DEGs from comparison of active beige and white adipocytes carrying FTO obesity-risk genotype; n=4 of each genotype

| Symbol | Gene Name | Log2 Fold Change | Adj. P value |
|---|---|---|---|
| *CPA4* | carboxypeptidase A4 | 2.21 | 6.62E-07 |
| *LAMP3* | lysosomal associated membrane protein 3 | 1.89 | 1.61E-04 |
| *GAP43* | growth associated protein 43 | 1.85 | 3.30E-04 |
| *FNDC1* | fibronectin type III domain containing 1 | 1.81 | 1.59E-04 |
| *SLC24A2* | solute carrier family 24 member 2 | 1.76 | 3.70E-04 |
| *MAP3K7CL* | MAP3K7 C-terminal like | 1.76 | 2.29E-07 |
| *C1QTNF3* | C1q and TNF related 3 | 1.69 | 1.93E-03 |
| *KRT79* | keratin 79 | 1.67 | 1.19E-03 |
| *THBD* | thrombomodulin | 1.61 | 4.84E-04 |
| *CEMIP* | cell migration inducing hyaluronidase 1 | 1.60 | 2.48E-04 |
| *CTXN1* | cortexin 1 | 1.58 | 0.001179796 |
| *RSPO3* | R-spondin 3 | 1.54 | 4.53E-03 |
| *PRAG1* | PEAK1 related, kinase-activating pseudokinase 1 | 1.53 | 0.000129348 |
| *KCNA1* | potassium voltage-gated channel subfamily A member 1 | 1.52 | 7.22E-03 |
| *KRT16* | keratin 16 | 1.51 | 1.84E-03 |
| *PCK1* | phosphoenolpyruvate carboxykinase 1 | 1.48 | 8.96E-03 |
| *TENT5C* | terminal nucleotidyltransferase 5C | 1.47 | 1.01E-02 |
| *CD96* | CD96 molecule | 1.47 | 1.32E-02 |
| *MEST* | mesoderm specific transcript | 1.45 | 0.000112085 |
| *MYL4* | myosin light chain 4 | 1.44 | 7.86E-03 |
| *CSDC2* | cold shock domain containing C2 | 1.43 | 0.000134369 |
| *AQP3* | aquaporin 3 (Gill blood group) | 1.43 | 1.14E-02 |
| *STAC* | SH3 and cysteine rich domain | 1.43 | 1.67558E-05 |
| *ZNF365* | zinc finger protein 365 | 1.39 | 4.32936E-05 |
| *TAGAP* | T cell activation RhoGTPase activating protein | 1.38 | 0.029446682 |
| *PM20D1* | peptidase M20 domain containing 1 | 1.36 | 0.032278284 |
| *PPP1R14C* | protein phosphatase 1 regulatory inhibitor subunit 14C | 1.34 | 4.18E-02 |
| *KIRREL3* | kirre like nephrin family adhesion molecule 3 | 1.34 | 2.45E-02 |
| *KCNK15* | potassium two pore domain channel subfamily K member 15 | 1.31 | 0.014097184 |

| | | | |
|---|---|---|---|
| *KCNMA1* | potassium calcium-activated channel subfamily M alpha 1 | 1.28 | 2.97E-02 |
| *RTKN2* | rhotekin 2 | 1.25 | 0.007842491 |
| *RBM24* | RNA binding motif protein 24 | 1.21 | 1.93E-07 |
| *SLITRK6* | SLIT and NTRK like family member 6 | 1.20 | 0.046666701 |
| *TNFAIP6* | TNF alpha induced protein 6 | 1.17 | 1.66E-02 |
| *LINGO1* | leucine rich repeat and Ig domain containing 1 | 1.17 | 0.027520144 |
| *CIART* | circadian associated repressor of transcription | 1.16 | 1.35919E-05 |
| *LINC00856* | long intergenic non-protein coding RNA 856 | 1.16 | 0.042053483 |
| *NDUFA4L2* | NDUFA4 mitochondrial complex associated like 2 | 1.15 | 1.06E-02 |
| *NA* | NA | 1.12 | 1.12E-04 |
| *APLP1* | amyloid beta precursor like protein 1 | 1.09 | 0.01078143 |
| *FUCA1* | alpha-L-fucosidase 1 | 1.09 | 8.13092E-05 |
| *DOK5* | docking protein 5 | 1.07 | 9.45044E-05 |
| *CYS1* | cystin 1 | 1.07 | 0.024443729 |
| *EPB41L4B* | erythrocyte membrane protein band 4.1 like 4B | 1.05 | 0.042053483 |
| *KCNG1* | potassium voltage-gated channel modifier subfamily G member 1 | 1.00 | 1.48E-02 |
| *SSC5D* | scavenger receptor cysteine rich family member with 5 domains | 0.99 | 2.45E-02 |
| *RCAN1* | regulator of calcineurin 1 | 0.99 | 0.00039172 |
| *TGM5* | transglutaminase 5 | 0.97 | 0.022767992 |
| *APBA2* | amyloid beta precursor protein binding family A member 2 | 0.94 | 1.22E-02 |
| *RNF122* | ring finger protein 122 | 0.91 | 1.53E-02 |
| *GALNT12* | polypeptide N-acetylgalactosaminyltransferase 12 | 0.88 | 0.00618527 |
| *GPC1* | glypican 1 | 0.87 | 0.019587135 |
| *STARD10* | StAR related lipid transfer domain containing 10 | 0.81 | 0.004855982 |
| *AMACR* | alpha-methylacyl-CoA racemase | 0.75 | 0.000368807 |
| *RFX2* | regulatory factor X2 | 0.71 | 0.027357964 |
| *ZBTB16* | zinc finger and BTB domain containing 16 | -4.95 | 3.4687E-100 |
| *HIF3A* | hypoxia inducible factor 3 subunit alpha | -3.26 | 4.89E-17 |
| *FKBP5* | FKBP prolyl isomerase 5 | -3.00 | 1.36855E-23 |

| | | | |
|---|---|---|---|
| *GALNT15* | polypeptide N-acetylgalactosaminyltransferase 15 | -2.96 | 1.42582E-21 |
| *GPX3* | glutathione peroxidase 3 | -2.72 | 3.01485E-14 |
| *GRIA1* | glutamate ionotropic receptor AMPA type subunit 1 | -2.60 | 1.03957E-26 |
| *CYP4B1* | cytochrome P450 family 4 subfamily B member 1 | -2.39 | 4.25677E-07 |
| *SLC16A12* | solute carrier family 16 member 12 | -2.39 | 1.83324E-07 |
| *C6* | complement C6 | -2.35 | 6.61625E-07 |
| *PER1* | period circadian regulator 1 | -2.31 | 1.22494E-22 |
| *PILRA* | paired immunoglobin like type 2 receptor alpha | -2.25 | 3.83627E-08 |
| *FMO2* | flavin containing dimethylaniline monoxygenase 2 | -2.18 | 2.55939E-06 |
| *GGT5* | gamma-glutamyltransferase 5 | -2.09 | 3.83627E-08 |
| *ABCC2* | ATP binding cassette subfamily C member 2 | -2.08 | 2.37257E-05 |
| *RAPGEF5* | Rap guanine nucleotide exchange factor 5 | -2.05 | 4.0043E-05 |
| *FGD4* | FYVE, RhoGEF and PH domain containing 4 | -2.05 | 3.46366E-08 |
| *MAOA* | monoamine oxidase A | -2.02 | 6.51851E-06 |
| *LRP1B* | LDL receptor related protein 1B | -1.99 | 8.24129E-05 |
| *METTL7A* | methyltransferase like 7A | -1.95 | 3.71531E-05 |
| *NEGR1* | neuronal growth regulator 1 | -1.91 | 1.33964E-11 |
| *TIMP4* | TIMP metallopeptidase inhibitor 4 | -1.87 | 0.000280383 |
| *LMO3* | LIM domain only 3 | -1.84 | 3.85E-06 |
| *F5* | coagulation factor V | -1.84 | 0.000416733 |
| *LEP* | leptin | -1.82 | 6.19839E-05 |
| *ALOX5AP* | arachidonate 5-lipoxygenase activating protein | -1.81 | 0.00048402 |
| *APCDD1* | APC down-regulated 1 | -1.80 | 1.67E-05 |
| *VIT* | vitrin | -1.78 | 2.65961E-13 |
| *PLXNA4* | plexin A4 | -1.78 | 4.29E-07 |
| *GLUL* | glutamate-ammonia ligase | -1.76 | 1.6253E-05 |
| *ANGPTL8* | angiopoietin like 8 | -1.75 | 0.000888082 |
| *ADH1A* | alcohol dehydrogenase 1A (class I), alpha polypeptide | -1.73 | 0.000242433 |
| *ADRA1B* | adrenoceptor alpha 1B | -1.72 | 2.63567E-06 |
| *SAA1* | serum amyloid A1 | -1.71 | 0.00138169 |
| *GPM6B* | glycoprotein M6B | -1.70 | 6.84565E-05 |

| | | | |
|---|---|---|---|
| *NKD2* | NKD inhibitor of WNT signaling pathway 2 | -1.70 | 0.001733466 |
| *CPM* | carboxypeptidase M | -1.66 | 0.003080267 |
| *EPHB6* | EPH receptor B6 | -1.65 | 0.000471449 |
| *HS3ST2* | heparan sulfate-glucosamine 3-sulfotransferase 2 | -1.65 | 3.16E-03 |
| *RASL11A* | RAS like family 11 member A | -1.65 | 0.000471449 |
| *INHBB* | inhibin subunit beta B | -1.64 | 0.000567616 |
| *CRISPLD2* | cysteine rich secretory protein LCCL domain containing 2 | -1.63 | 9.97296E-05 |
| *NA* | NA | -1.62 | 4.21E-03 |
| *PRODH* | proline dehydrogenase 1 | -1.60 | 0.004211143 |
| *APOB* | apolipoprotein B | -1.60 | 0.004638876 |
| *SCARA5* | scavenger receptor class A member 5 | -1.59 | 0.004527457 |
| *DPEP1* | dipeptidase 1 | -1.58 | 0.006228468 |
| *DAAM2-AS1* | DAAM2 antisense RNA 1 | -1.57 | 2.28E-03 |
| *MAOB* | monoamine oxidase B | -1.56 | 5.34E-03 |
| *RGCC* | regulator of cell cycle | -1.56 | 0.006255759 |
| *KIAA0040* | KIAA0040 | -1.56 | 6.83E-03 |
| *TMPRSS5* | transmembrane serine protease 5 | -1.55 | 0.006074323 |
| *LRRN3* | leucine rich repeat neuronal 3 | -1.55 | 0.003577777 |
| *TMC2* | transmembrane channel like 2 | -1.53 | 0.007842491 |
| *ALS2CL* | ALS2 C-terminal like | -1.48 | 0.011813178 |
| *COL4A6* | collagen type IV alpha 6 chain | -1.48 | 0.011623143 |
| *CD34* | CD34 molecule | -1.48 | 0.012930564 |
| *GDF7* | growth differentiation factor 7 | -1.48 | 0.002574005 |
| *PTPN22* | protein tyrosine phosphatase non-receptor type 22 | -1.46 | 0.015300372 |
| *GUCY1A1* | guanylate cyclase 1 soluble subunit alpha 1 | -1.46 | 6.84565E-05 |
| *STC1* | stanniocalcin 1 | -1.46 | 0.016475532 |
| *LINC01088* | long intergenic non-protein coding RNA 1088 | -1.46 | 0.011477841 |
| *NA* | NA | -1.45 | 0.017347738 |
| *RIPOR3* | RIPOR family member 3 | -1.45 | 0.017043932 |
| *ANGPTL1* | angiopoietin like 1 | -1.44 | 0.000553976 |
| *AZGP1* | alpha-2-glycoprotein 1, zinc-binding | -1.44 | 0.018535439 |
| *LINC00482* | long intergenic non-protein coding RNA 482 | -1.44 | 0.016475532 |
| *APOD* | apolipoprotein D | -1.44 | 0.019587135 |
| *TMEM150C* | transmembrane protein 150C | -1.43 | 0.011354892 |

| | | | |
|---|---|---|---|
| *PKD2L1* | polycystin 2 like 1, transient receptor potential cation channel | -1.42 | 0.016475532 |
| *TSC22D3* | TSC22 domain family member 3 | -1.40 | 3.77338E-08 |
| *POM121L9P* | POM121 transmembrane nucleoporin like 9, pseudogene | -1.39 | 0.006228468 |
| *ADH1B* | alcohol dehydrogenase 1B (class I), beta polypeptide | -1.39 | 0.015862261 |
| *MRO* | maestro | -1.37 | 0.033330958 |
| *GUCY1B1* | guanylate cyclase 1 soluble subunit beta 1 | -1.36 | 0.000327154 |
| *CILP* | cartilage intermediate layer protein | -1.34 | 0.03989726 |
| *POU3F3* | POU class 3 homeobox 3 | -1.34 | 0.029446682 |
| *FAM166B* | family with sequence similarity 166 member B | -1.34 | 0.042868397 |
| *SMCO3* | single-pass membrane protein with coiled-coil domains 3 | -1.33 | 0.018465442 |
| *SERPINA3* | serpin family A member 3 | -1.32 | 0.046666701 |
| *NRCAM* | neuronal cell adhesion molecule | -1.29 | 0.026482246 |
| *SLC34A2* | solute carrier family 34 member 2 | -1.27 | 0.049483648 |
| *PRELP* | proline and arginine rich end leucine rich repeat protein | -1.26 | 3.80272E-05 |
| *WASF3* | WASP family member 3 | -1.26 | 3.57256E-14 |
| *RASSF4* | Ras association domain family member 4 | -1.26 | 0.015366962 |
| *ADARB1* | adenosine deaminase RNA specific B1 | -1.26 | 0.000183071 |
| *NA* | NA | -1.25 | 0.033654555 |
| *MGP* | matrix Gla protein | -1.25 | 0.024534915 |
| *DHRS3* | dehydrogenase/reductase 3 | -1.23 | 0.003152659 |
| *MT1X* | metallothionein 1X | -1.17 | 0.012510803 |
| *BOC* | BOC cell adhesion associated, oncogene regulated | -1.17 | 0.005972864 |
| *KLF9* | Kruppel like factor 9 | -1.17 | 1.0158E-06 |
| *SIM1* | SIM bHLH transcription factor 1 | -1.16 | 0.046781488 |
| *CFD* | complement factor D | -1.15 | 0.002412682 |
| *ERRFI1* | ERBB receptor feedback inhibitor 1 | -1.14 | 0.000247582 |
| *RGMA* | repulsive guidance molecule BMP co-receptor a | -1.13 | 0.004979066 |
| *TENT5B* | terminal nucleotidyltransferase 5B | -1.12 | 0.015300372 |
| *IRS1* | insulin receptor substrate 1 | -1.11 | 0.002530607 |
| *CLCN4* | chloride voltage-gated channel 4 | -1.11 | 0.000108217 |
| *SSH2* | slingshot protein phosphatase 2 | -1.08 | 0.000160562 |
| *CST3* | cystatin C | -1.08 | 0.004211143 |

| | | | |
|---|---|---|---|
| *GREB1L* | GREB1 like retinoic acid receptor coactivator | -1.06 | 0.015300372 |
| *RNASE4* | ribonuclease A family member 4 | -1.04 | 1.67319E-05 |
| *NID1* | nidogen 1 | -1.04 | 5.48378E-06 |
| *COL5A3* | collagen type V alpha 3 chain | -1.03 | 0.020208861 |
| *TGFBR2* | transforming growth factor beta receptor 2 | -1.02 | 3.77338E-08 |
| *WNT11* | Wnt family member 11 | -1.01 | 0.015862261 |
| *SOX13* | SRY-box transcription factor 13 | -1.00 | 3.80152E-05 |
| *ZNF395* | zinc finger protein 395 | -1.00 | 0.020208861 |
| *TNS2* | tensin 2 | -1.00 | 0.000678684 |
| *CUTC* | cutC copper transporter | -0.98 | 7.37E-07 |
| *DSEL* | dermatan sulfate epimerase like | -0.97 | 0.027576633 |
| *ALDH16A1* | aldehyde dehydrogenase 16 family member A1 | -0.96 | 0.009618862 |
| *CXCL12* | C-X-C motif chemokine ligand 12 | -0.96 | 2.22858E-05 |
| *TLR4* | toll like receptor 4 | -0.94 | 0.004438033 |
| *CHST2* | carbohydrate sulfotransferase 2 | -0.93 | 0.027979782 |
| *TACC1* | transforming acidic coiled-coil containing protein 1 | -0.92 | 3.74E-03 |
| *SMARCD2* | SWI/SNF related, matrix associated, actin dependent regulator of chromatin, subfamily d, member 2 | -0.90 | 0.00039172 |
| *JADE2* | jade family PHD finger 2 | -0.90 | 0.040374979 |
| *HIPK2* | homeodomain interacting protein kinase 2 | -0.90 | 0.011742751 |
| *ITPR1* | inositol 1,4,5-trisphosphate receptor type 1 | -0.90 | 0.030465421 |
| *ANG* | angiogenin | -0.88 | 0.040374979 |
| *ZNF438* | zinc finger protein 438 | -0.85 | 0.036550321 |
| *MINDY2* | MINDY lysine 48 deubiquitinase 2 | -0.84 | 0.009618862 |
| *HLX* | H2.0 like homeobox | -0.83 | 7.46E-03 |
| *CAT* | catalase | -0.82 | 0.029463934 |
| *TTPAL* | alpha tocopherol transfer protein like | -0.81 | 0.006334513 |
| *DNAJC6* | DnaJ heat shock protein family (Hsp40) member C6 | -0.81 | 0.046666701 |
| *CFLAR* | CASP8 and FADD like apoptosis regulator | -0.79 | 4.86E-03 |
| *FOXO1* | forkhead box O1 | -0.78 | 9.70E-03 |
| *FOXO3* | forkhead box O3 | -0.77 | 9.45044E-05 |
| *FZD4* | frizzled class receptor 4 | -0.76 | 4.12E-02 |
| *PTK2B* | protein tyrosine kinase 2 beta | -0.75 | 6.26E-03 |

Supplementary Table 8 DEGs from comparison of active beige and inactive beige adipocytes carrying FTO risk-free genotype; n=4 of each genotype

| Symbol | Gene Name | Log2 Fold Change | Adj. P value |
|---|---|---|---|
| *PCK1* | phosphoenolpyruvate carboxykinase 1 | 2.41 | 1.04E-15 |
| *CPA4* | carboxypeptidase A4 | 2.32 | 1.65E-14 |
| *SCN4A* | sodium voltage-gated channel alpha subunit 4 | 2.20 | 4.18E-11 |
| *KRT79* | keratin 79 | 2.14 | 2.08E-10 |
| *SLC22A12* | solute carrier family 22 member 12 | 1.99 | 7.43E-09 |
| *NA* | NA | 1.92 | 1.18E-08 |
| *BBOX1* | gamma-butyrobetaine hydroxylase 1 | 1.91 | 7.73E-10 |
| *ADAMTS18* | ADAM metallopeptidase with thrombospondin type 1 motif 18 | 1.77 | 4.53E-07 |
| *FCN1* | ficolin 1 | 1.77 | 5.43E-07 |
| *TRPM8* | transient receptor potential cation channel subfamily M member 8 | 1.66 | 4.26E-06 |
| *SLC22A3* | solute carrier family 22 member 3 | 1.63 | 2.57558E-08 |
| *PPP1R1A* | protein phosphatase 1 regulatory inhibitor subunit 1A | 1.63 | 1.70E-08 |
| *CITED1* | Cbp/p300 interacting transactivator with Glu/Asp rich carboxy-terminal domain 1 | 1.62 | 3.88183E-07 |
| *FABP4* | fatty acid binding protein 4 | 1.59 | 5.18E-09 |
| *PDE1B* | phosphodiesterase 1B | 1.56 | 4.79E-08 |
| *AP3B2* | adaptor related protein complex 3 subunit beta 2 | 1.56 | 4.37E-12 |
| *SORL1* | sortilin related receptor 1 | 1.55 | 2.83E-05 |
| *AQP3* | aquaporin 3 (Gill blood group) | 1.54 | 2.55E-05 |
| *GK* | glycerol kinase | 1.54 | 7.39062E-10 |
| *IL1B* | interleukin 1 beta | 1.50 | 1.70E-05 |
| *CD52* | CD52 molecule | 1.49 | 7.22291E-05 |
| *LPL* | lipoprotein lipase | 1.49 | 1.54E-06 |
| *CLDN2* | claudin 2 | 1.47 | 7.157E-05 |
| *KCNA1* | potassium voltage-gated channel subfamily A member 1 | 1.46 | 6.87598E-05 |
| *CTXN1* | cortexin 1 | 1.45 | 1.22527E-07 |
| *PLEKHG6* | pleckstrin homology and RhoGEF domain containing G6 | 1.44 | 3.88183E-07 |
| *PLIN5* | perilipin 5 | 1.41 | 6.00E-05 |
| *KIRREL3* | kirre like nephrin family adhesion molecule 3 | 1.40 | 6.11E-05 |

| | | | |
|---|---|---|---|
| *PM20D1* | peptidase M20 domain containing 1 | 1.39 | 0.000156088 |
| *CYP4F12* | cytochrome P450 family 4 subfamily F member 12 | 1.37 | 2.10E-05 |
| *KCNK12* | potassium two pore domain channel subfamily K member 12 | 1.37 | 0.000368849 |
| *LHCGR* | luteinizing hormone/choriogonadotropin receptor | 1.37 | 1.68E-04 |
| *ANO5* | anoctamin 5 | 1.36 | 0.000205831 |
| *ADIPOQ* | adiponectin, C1Q and collagen domain containing | 1.34 | 1.98E-04 |
| *LINC02554* | long intergenic non-protein coding RNA 2554 | 1.33 | 0.000503395 |
| *TMEM132C* | transmembrane protein 132C | 1.33 | 0.000234679 |
| *TSHR* | thyroid stimulating hormone receptor | 1.33 | 0.000613382 |
| *CD96* | CD96 molecule | 1.32 | 6.09E-04 |
| *C1QTNF3* | C1q and TNF related 3 | 1.32 | 5.87E-04 |
| *ADAM12* | ADAM metallopeptidase domain 12 | 1.30 | 0.000126762 |
| *CASZ1* | castor zinc finger 1 | 1.29 | 0.001217157 |
| *LUCAT1* | lung cancer associated transcript 1 | 1.28 | 0.001332685 |
| *STX11* | syntaxin 11 | 1.28 | 1.23367E-05 |
| *TRHDE-AS1* | TRHDE antisense RNA 1 | 1.27 | 3.22087E-10 |
| *ABCG1* | ATP binding cassette subfamily G member 1 | 1.27 | 5.95E-06 |
| *EPB41L4B* | erythrocyte membrane protein band 4.1 like 4B | 1.25 | 4.94E-04 |
| *TMEM200C* | transmembrane protein 200C | 1.24 | 0.001474214 |
| *THBD* | thrombomodulin | 1.24 | 0.001938233 |
| *MOGAT1* | monoacylglycerol O-acyltransferase 1 | 1.21 | 3.13E-03 |
| *MAP3K7CL* | MAP3K7 C-terminal like | 1.21 | 6.11E-05 |
| *FNDC1* | fibronectin type III domain containing 1 | 1.21 | 0.00336912 |
| *SLCO2A1* | solute carrier organic anion transporter family member 2A1 | 1.19 | 0.003487693 |
| *LRRC15* | leucine rich repeat containing 15 | 1.19 | 0.00418377 |
| *RUBCNL* | rubicon like autophagy enhancer | 1.18 | 0.000440607 |
| *PCSK1* | proprotein convertase subtilisin/kexin type 1 | 1.17 | 0.001533118 |
| *SCAMP5* | secretory carrier membrane protein 5 | 1.17 | 0.002084765 |
| *ICA1* | islet cell autoantigen 1 | 1.17 | 1.22E-03 |
| *RASD1* | ras related dexamethasone induced 1 | 1.15 | 1.23367E-05 |
| *NUDT14* | nudix hydrolase 14 | 1.14 | 8.99329E-06 |

| | | | |
|---|---|---|---|
| *GRAP* | GRB2 related adaptor protein | 1.14 | 0.005592092 |
| *TENT5C* | terminal nucleotidyltransferase 5C | 1.14 | 0.006357465 |
| *FGR* | FGR proto-oncogene, Src family tyrosine kinase | 1.13 | 6.11052E-05 |
| *ADGRB2* | adhesion G protein-coupled receptor B2 | 1.13 | 0.000831457 |
| *EGFLAM* | EGF like, fibronectin type III and laminin G domains | 1.13 | 0.000673902 |
| *MGAT3* | beta-1,4-mannosyl-glycoprotein 4-beta-N-acetylglucosaminyltransferase | 1.13 | 0.00056257 |
| *DLX3* | distal-less homeobox 3 | 1.13 | 0.008700818 |
| *TRHDE* | thyrotropin releasing hormone degrading enzyme | 1.12 | 0.000268661 |
| *RSPO3* | R-spondin 3 | 1.11 | 0.008700818 |
| *TSPAN15* | tetraspanin 15 | 1.11 | 0.004062811 |
| *FCHO1* | FCH and mu domain containing endocytic adaptor 1 | 1.10 | 0.001715175 |
| *EPB41* | erythrocyte membrane protein band 4.1 | 1.10 | 3.45134E-17 |
| *ADAMTS14* | ADAM metallopeptidase with thrombospondin type 1 motif 14 | 1.10 | 0.011720639 |
| *ECSCR* | endothelial cell surface expressed chemotaxis and apoptosis regulator | 1.09 | 0.010950504 |
| *CYS1* | cystin 1 | 1.09 | 0.001547479 |
| *AATBC* | apoptosis associated transcript in bladder cancer | 1.09 | 0.012342268 |
| *ACSM3* | acyl-CoA synthetase medium chain family member 3 | 1.08 | 0.009975763 |
| *PALMD* | palmdelphin | 1.08 | 3.49E-03 |
| *CMYA5* | cardiomyopathy associated 5 | 1.08 | 0.001618637 |
| *PREX1* | phosphatidylinositol-3,4,5-trisphosphate dependent Rac exchange factor 1 | 1.07 | 0.000241977 |
| *RBM24* | RNA binding motif protein 24 | 1.06 | 0.010697451 |
| *RTKN2* | rhotekin 2 | 1.06 | 1.05E-02 |
| *COL8A2* | collagen type VIII alpha 2 chain | 1.06 | 2.77185E-09 |
| *PER3* | period circadian regulator 3 | 1.06 | 3.85E-10 |
| *ITIH1* | inter-alpha-trypsin inhibitor heavy chain 1 | 1.05 | 0.013689471 |
| *PHLDA1* | pleckstrin homology like domain family A member 1 | 1.05 | 0.00147364 |
| *SV2A* | synaptic vesicle glycoprotein 2A | 1.04 | 0.019965933 |
| *PCDH19* | protocadherin 19 | 1.03 | 0.0159972 |

| | | | |
|---|---|---|---|
| *MEST* | mesoderm specific transcript | 1.03 | 0.005849882 |
| *CKMT1B* | creatine kinase, mitochondrial 1B | 1.03 | 0.022431637 |
| *SH2D3C* | SH2 domain containing 3C | 1.03 | 0.005279943 |
| *KRT14* | keratin 14 | 1.02 | 0.005819803 |
| *LMO2* | LIM domain only 2 | 1.02 | 0.008648955 |
| *PLIN4* | perilipin 4 | 1.02 | 1.92E-02 |
| *SRCIN1* | SRC kinase signaling inhibitor 1 | 1.01 | 0.011926948 |
| *IL4I1* | interleukin 4 induced 1 | 1.01 | 0.028950887 |
| *SLC19A3* | solute carrier family 19 member 3 | 1.00 | 0.00764458 |
| *ABCA3* | ATP binding cassette subfamily A member 3 | 1.00 | 1.36E-02 |
| *GXYLT2* | glucoside xylosyltransferase 2 | 0.99 | 0.000229046 |
| *CIART* | circadian associated repressor of transcription | 0.99 | 0.004139608 |
| *CKMT2* | creatine kinase, mitochondrial 2 | 0.99 | 0.002462438 |
| *RXRG* | retinoid X receptor gamma | 0.99 | 0.004055711 |
| *RGS7BP* | regulator of G protein signaling 7 binding protein | 0.98 | 5.90E-03 |
| *FBXO27* | F-box protein 27 | 0.98 | 8.39E-04 |
| *CIDEA* | cell death inducing DFFA like effector a | 0.98 | 0.032303887 |
| *APLP1* | amyloid beta precursor like protein 1 | 0.98 | 1.41E-04 |
| *NA* | NA | 0.98 | 0.028202928 |
| *CADM3* | cell adhesion molecule 3 | 0.98 | 0.028950887 |
| *RGS2* | regulator of G protein signaling 2 | 0.98 | 0.000229046 |
| *ASS1* | argininosuccinate synthase 1 | 0.97 | 0.000201889 |
| *MACROD2* | mono-ADP ribosylhydrolase 2 | 0.97 | 0.015219886 |
| *PLEKHA7* | pleckstrin homology domain containing A7 | 0.96 | 0.012153597 |
| *AQP7* | aquaporin 7 | 0.96 | 0.000388439 |
| *RIMS4* | regulating synaptic membrane exocytosis 4 | 0.96 | 0.039917984 |
| *DPYSL3* | dihydropyrimidinase like 3 | 0.96 | 0.000860395 |
| *LGI2* | leucine rich repeat LGI family member 2 | 0.96 | 0.041785315 |
| *FAM151A* | family with sequence similarity 151 member A | 0.96 | 0.036439624 |
| *C3orf80* | chromosome 3 open reading frame 80 | 0.95 | 0.045925757 |
| *PDZD2* | PDZ domain containing 2 | 0.95 | 0.032874973 |
| *FYB1* | FYN binding protein 1 | 0.95 | 0.001975791 |
| *PRG4* | proteoglycan 4 | 0.95 | 0.046393557 |
| *GPR153* | G protein-coupled receptor 153 | 0.94 | 1.47895E-09 |

| | | | |
|---|---|---|---|
| *LBP* | lipopolysaccharide binding protein | 0.94 | 0.033799174 |
| *RGL3* | ral guanine nucleotide dissociation stimulator like 3 | 0.94 | 0.029610558 |
| *DHCR7* | 7-dehydrocholesterol reductase | 0.93 | 3.88183E-07 |
| *GCOM1* | GRINL1A complex locus 1 | 0.93 | 0.00955761 |
| *SEL1L3* | SEL1L family member 3 | 0.93 | 0.015561115 |
| *SYNGR2* | synaptogyrin 2 | 0.93 | 0.004062811 |
| *PPM1H* | protein phosphatase, Mg2+/Mn2+ dependent 1H | 0.93 | 0.00122567 |
| *PLXDC1* | plexin domain containing 1 | 0.92 | 0.005849882 |
| *SSC5D* | scavenger receptor cysteine rich family member with 5 domains | 0.92 | 0.006853399 |
| *CA3* | carbonic anhydrase 3 | 0.92 | 0.044090769 |
| *HCAR3* | hydroxycarboxylic acid receptor 3 | 0.91 | 0.036923788 |
| *GPC1* | glypican 1 | 0.90 | 0.000395719 |
| *KIAA1549L* | KIAA1549 like | 0.89 | 0.001248166 |
| *TMEM131L* | transmembrane 131 like | 0.89 | 0.000592826 |
| *FLVCR2* | FLVCR heme transporter 2 | 0.89 | 0.043463848 |
| *MRAP* | melanocortin 2 receptor accessory protein | 0.88 | 0.012868285 |
| *TDRKH* | tudor and KH domain containing | 0.88 | 0.004519579 |
| *SAMD14* | sterile alpha motif domain containing 14 | 0.87 | 0.012941161 |
| *TENM4* | teneurin transmembrane protein 4 | 0.87 | 0.025884593 |
| *PTGER2* | prostaglandin E receptor 2 | 0.87 | 0.046393557 |
| *AFF2* | AF4/FMR2 family member 2 | 0.87 | 0.008077169 |
| *SLC19A1* | solute carrier family 19 member 1 | 0.86 | 6.93268E-05 |
| *ACAA2* | acetyl-CoA acyltransferase 2 | 0.86 | 0.000388058 |
| *NR3C2* | nuclear receptor subfamily 3 group C member 2 | 0.86 | 0.002004975 |
| *TTC9* | tetratricopeptide repeat domain 9 | 0.86 | 0.039296395 |
| *DOK5* | docking protein 5 | 0.85 | 1.39055E-05 |
| *DIRAS1* | DIRAS family GTPase 1 | 0.85 | 0.037709719 |
| *ASPHD2* | aspartate beta-hydroxylase domain containing 2 | 0.84 | 0.003345262 |
| *NR1D2* | nuclear receptor subfamily 1 group D member 2 | 0.84 | 5.04452E-12 |
| *FAR2* | fatty acyl-CoA reductase 2 | 0.83 | 0.021325265 |
| *DOCK10* | dedicator of cytokinesis 10 | 0.82 | 0.000727254 |
| *GUCY1A2* | guanylate cyclase 1 soluble subunit alpha 2 | 0.82 | 0.000319778 |

| | | | |
|---|---|---|---|
| *MKNK2* | MAPK interacting serine/threonine kinase 2 | 0.81 | 0.000759138 |
| *LACC1* | laccase domain containing 1 | 0.81 | 0.024633811 |
| *ACOT2* | acyl-CoA thioesterase 2 | 0.79 | 0.021010834 |
| *ATP8B1* | ATPase phospholipid transporting 8B1 | 0.78 | 1.23367E-05 |
| *MLYCD* | malonyl-CoA decarboxylase | 0.77 | 0.002234209 |
| *PPIP5K1* | diphosphoinositol pentakisphosphate kinase 1 | 0.76 | 0.002268177 |
| *NCS1* | neuronal calcium sensor 1 | 0.76 | 1.23367E-05 |
| *LAIR1* | leukocyte associated immunoglobulin like receptor 1 | 0.76 | 0.018875982 |
| *ANGPTL4* | angiopoietin like 4 | 0.76 | 0.02339336 |
| *L3MBTL2-AS1* | L3MBTL2 antisense RNA 1 | 0.76 | 0.040013166 |
| *CAMKK1* | calcium/calmodulin dependent protein kinase kinase 1 | 0.76 | 0.000696505 |
| *PTGDS* | prostaglandin D2 synthase | 0.75 | 9.57E-03 |
| *HILPDA* | hypoxia inducible lipid droplet associated | 0.75 | 0.001810421 |
| *LRBA* | LPS responsive beige-like anchor protein | 0.73 | 0.000130463 |
| *PIK3R2* | phosphoinositide-3-kinase regulatory subunit 2 | 0.73 | 0.000140823 |
| *ITGA4* | integrin subunit alpha 4 | 0.73 | 0.041203422 |
| *PEX11A* | peroxisomal biogenesis factor 11 alpha | 0.71 | 0.036796195 |
| *SHB* | SH2 domain containing adaptor protein B | 0.71 | 3.64E-02 |
| *ORAI2* | ORAI calcium release-activated calcium modulator 2 | 0.70 | 0.027435891 |
| *ZBTB16* | zinc finger and BTB domain containing 16 | -3.87 | 6.25789E-60 |
| *PER1* | period circadian regulator 1 | -2.46 | 1.256E-100 |
| *FKBP5* | FKBP prolyl isomerase 5 | -2.07 | 2.89729E-11 |
| *GABRA5* | gamma-aminobutyric acid type A receptor subunit alpha5 | -1.94 | 2.12939E-08 |
| *TIMP4* | TIMP metallopeptidase inhibitor 4 | -1.79 | 3.84076E-07 |
| *HIF3A* | hypoxia inducible factor 3 subunit alpha | -1.79 | 3.14282E-07 |
| *MMP28* | matrix metallopeptidase 28 | -1.77 | 5.85E-08 |
| *INHBB* | inhibin subunit beta B | -1.76 | 3.97426E-08 |
| *ACKR2* | atypical chemokine receptor 2 | -1.73 | 6.90845E-13 |

| | | | |
|---|---|---|---|
| *PILRA* | paired immunoglobin like type 2 receptor alpha | -1.72 | 1.14664E-06 |
| *ANGPTL8* | angiopoietin like 8 | -1.70 | 2.08E-06 |
| *TMC2* | transmembrane channel like 2 | -1.68 | 1.11E-06 |
| *TENT5B* | terminal nucleotidyltransferase 5B | -1.65 | 6.41146E-10 |
| *NKD2* | NKD inhibitor of WNT signaling pathway 2 | -1.59 | 3.45E-17 |
| *DPT* | dermatopontin | -1.59 | 3.14E-07 |
| *C6* | complement C6 | -1.58 | 1.45E-05 |
| *ABCC2* | ATP binding cassette subfamily C member 2 | -1.58 | 1.30E-05 |
| *CRLF1* | cytokine receptor like factor 1 | -1.57 | 9.39E-09 |
| *APCDD1* | APC down-regulated 1 | -1.55 | 2.66E-10 |
| *CRISPLD2* | cysteine rich secretory protein LCCL domain containing 2 | -1.52 | 3.84E-07 |
| *LMO3* | LIM domain only 3 | -1.50 | 3.88E-07 |
| *CPM* | carboxypeptidase M | -1.49 | 2.22E-07 |
| *SAA1* | serum amyloid A1 | -1.49 | 3.08E-05 |
| *MAOA* | monoamine oxidase A | -1.47 | 2.94745E-05 |
| *NEGR1* | neuronal growth regulator 1 | -1.47 | 1.18E-08 |
| *SYN2* | synapsin II | -1.45 | 0.000125717 |
| *ATP1A2* | ATPase Na+/K+ transporting subunit alpha 2 | -1.41 | 0.000229046 |
| *PTGS1* | prostaglandin-endoperoxide synthase 1 | -1.40 | 1.09E-08 |
| *RASL11A* | RAS like family 11 member A | -1.37 | 1.57E-05 |
| *TSC22D3* | TSC22 domain family member 3 | -1.37 | 3.69E-14 |
| *AVPR1A* | arginine vasopressin receptor 1A | -1.34 | 6.05E-04 |
| *ANGPTL1* | angiopoietin like 1 | -1.24 | 6.11E-05 |
| *ISM1* | isthmin 1 | -1.23 | 0.002115835 |
| *TRNP1* | TMF1 regulated nuclear protein 1 | -1.22 | 1.76E-10 |
| *GLUL* | glutamate-ammonia ligase | -1.21 | 8.41E-05 |
| *CYP8B1* | cytochrome P450 family 8 subfamily B member 1 | -1.18 | 0.004345624 |
| *METTL7A* | methyltransferase like 7A | -1.17 | 1.47E-03 |
| *SLC16A12* | solute carrier family 16 member 12 | -1.16 | 0.004232885 |
| *GALNT15* | polypeptide N-acetylgalactosaminyltransferase 15 | -1.16 | 0.00514905 |
| *STC1* | stanniocalcin 1 | -1.15 | 0.005819803 |
| *FMO2* | flavin containing dimethylaniline monoxygenase 2 | -1.11 | 0.008659854 |
| *PLXNA4* | plexin A4 | -1.11 | 7.07688E-06 |

| | | | |
|---|---|---|---|
| *TMPRSS5* | transmembrane serine protease 5 | -1.10 | 0.010950504 |
| *SPARCL1* | SPARC like 1 | -1.10 | 3.29E-03 |
| *SMOC2* | SPARC related modular calcium binding 2 | -1.10 | 1.91E-04 |
| *LRP1B* | LDL receptor related protein 1B | -1.09 | 0.00514905 |
| *HPD* | 4-hydroxyphenylpyruvate dioxygenase | -1.08 | 0.013383227 |
| *GLYAT* | glycine-N-acyltransferase | -1.08 | 0.008749031 |
| *ELANE* | elastase, neutrophil expressed | -1.07 | 1.88E-04 |
| *F5* | coagulation factor V | -1.07 | 0.015627188 |
| *MT1X* | metallothionein 1X | -1.07 | 0.001130893 |
| *SERPINA3* | serpin family A member 3 | -1.06 | 0.013154664 |
| *PKD2L1* | polycystin 2 like 1, transient receptor potential cation channel | -1.06 | 1.19E-02 |
| *MGP* | matrix Gla protein | -1.05 | 0.016714274 |
| *WNT5A* | Wnt family member 5A | -1.05 | 0.010950504 |
| *DUSP4* | dual specificity phosphatase 4 | -1.04 | 0.000171262 |
| *PRODH* | proline dehydrogenase 1 | -1.04 | 0.018449532 |
| *KLF9* | Kruppel like factor 9 | -1.04 | 2.17E-07 |
| *GGT5* | gamma-glutamyltransferase 5 | -1.03 | 0.021127555 |
| *TMEM59L* | transmembrane protein 59 like | -1.03 | 9.34E-04 |
| *TF* | transferrin | -1.03 | 0.019267474 |
| *GSN* | gelsolin | -1.02 | 6.79E-03 |
| *PIK3R1* | phosphoinositide-3-kinase regulatory subunit 1 | -1.01 | 5.90E-04 |
| *CST3* | cystatin C | -1.00 | 0.010233937 |
| *RGMA* | repulsive guidance molecule BMP co-receptor a | -0.99 | 9.37536E-12 |
| *KIAA0040* | KIAA0040 | -0.98 | 0.033799174 |
| *CORO6* | coronin 6 | -0.98 | 0.010092135 |
| *TTYH1* | tweety family member 1 | -0.98 | 0.037970196 |
| *TAFA2* | TAFA chemokine like family member 2 | -0.97 | 1.25E-02 |
| *TRARG1* | trafficking regulator of GLUT4 (SLC2A4) 1 | -0.97 | 0.011926948 |
| *RASSF4* | Ras association domain family member 4 | -0.96 | 2.66E-04 |
| *SMARCD2* | SWI/SNF related, matrix associated, actin dependent regulator of chromatin, subfamily d, member 2 | -0.96 | 1.41963E-08 |
| *MAP2K6* | mitogen-activated protein kinase kinase 6 | -0.95 | 0.048141751 |

| | | | |
|---|---|---|---|
| *CACNB2* | calcium voltage-gated channel auxiliary subunit beta 2 | -0.95 | 0.009011383 |
| *TC2N* | tandem C2 domains, nuclear | -0.94 | 0.024554373 |
| *RAPGEF5* | Rap guanine nucleotide exchange factor 5 | -0.93 | 0.042389959 |
| *ERRFI1* | ERBB receptor feedback inhibitor 1 | -0.93 | 0.001332685 |
| *PRRT4* | proline rich transmembrane protein 4 | -0.93 | 0.003253894 |
| *HLX* | H2.0 like homeobox | -0.92 | 1.57858E-05 |
| *ANPEP* | alanyl aminopeptidase, membrane | -0.92 | 0.01006536 |
| *C1QTNF7* | C1q and TNF related 7 | -0.92 | 0.02339336 |
| *SLC2A13* | solute carrier family 2 member 13 | -0.91 | 0.046393557 |
| *BOC* | BOC cell adhesion associated, oncogene regulated | -0.91 | 0.032358485 |
| *NA* | NA | -0.91 | 0.048654705 |
| *SAMHD1* | SAM and HD domain containing deoxynucleoside triphosphate triphosphohydrolase 1 | -0.90 | 0.022109898 |
| *LAMA2* | laminin subunit alpha 2 | -0.89 | 0.000651072 |
| *ZNF395* | zinc finger protein 395 | -0.89 | 3.46852E-17 |
| *AACS* | acetoacetyl-CoA synthetase | -0.88 | 3.07E-02 |
| *CUTC* | cutC copper transporter | -0.87 | 0.008083892 |
| *STK17B* | serine/threonine kinase 17b | -0.87 | 0.020867511 |
| *AOX1* | aldehyde oxidase 1 | -0.86 | 0.045804255 |
| *OLFM2* | olfactomedin 2 | -0.86 | 0.000184629 |
| *GRK5* | G protein-coupled receptor kinase 5 | -0.86 | 0.008659854 |
| *LAMA3* | laminin subunit alpha 3 | -0.86 | 0.015627188 |
| *TMEM64* | transmembrane protein 64 | -0.86 | 0.009583854 |
| *PDE8B* | phosphodiesterase 8B | -0.83 | 6.81E-03 |
| *CDKN1C* | cyclin dependent kinase inhibitor 1C | -0.83 | 2.90E-02 |
| *HRCT1* | histidine rich carboxyl terminus 1 | -0.82 | 0.034232774 |
| *DNASE1L3* | deoxyribonuclease 1 like 3 | -0.82 | 0.045292682 |
| *TNS2* | tensin 2 | -0.81 | 2.57558E-08 |
| *SSH2* | slingshot protein phosphatase 2 | -0.81 | 0.02224363 |
| *GNG2* | G protein subunit gamma 2 | -0.80 | 0.032675774 |
| *IQCH-AS1* | IQCH antisense RNA 1 | -0.79 | 0.002346016 |
| *SOX13* | SRY-box transcription factor 13 | -0.79 | 0.016610242 |
| *SYNE2* | spectrin repeat containing nuclear envelope protein 2 | -0.79 | 0.047078744 |
| *WASF3* | WASP family member 3 | -0.79 | 2.61234E-05 |
| *RNASE4* | ribonuclease A family member 4 | -0.78 | 0.029687428 |
| *TGFBR2* | transforming growth factor beta receptor 2 | -0.77 | 4.43326E-10 |

| | | | |
|---|---|---|---|
| *NID1* | nidogen 1 | -0.74 | 1.94E-03 |
| *DUSP1* | dual specificity phosphatase 1 | -0.73 | 0.001559477 |
| *ZFP36* | ZFP36 ring finger protein | -0.73 | 0.001938233 |
| *MFGE8* | milk fat globule EGF and factor V/VIII domain containing | -0.72 | 0.002011495 |
| *CAB39L* | calcium binding protein 39 like | -0.72 | 0.039039489 |
| *DHRS3* | dehydrogenase/reductase 3 | -0.72 | 0.00122567 |
| *SLC25A10* | solute carrier family 25 member 10 | -0.71 | 0.047718206 |
| *EVA1C* | eva-1 homolog C | -0.71 | 0.048862069 |

Supplementary Table 9 DEGs from comparison of active beige and inactive beige adipocytes carrying FTO obesity-risk genotype; n=4 of each genotype

| Symbol | Gene Name | Log2 Fold Change | Adj. P value |
|---|---|---|---|
| *CPA4* | carboxypeptidase A4 | 1.53 | 2.12E-04 |
| *MAP3K7CL* | MAP3K7 C-terminal like | 1.42 | 2.60E-05 |
| *C3orf80* | chromosome 3 open reading frame 80 | 1.34 | 1.51E-03 |
| *FNDC1* | fibronectin type III domain containing 1 | 1.31 | 4.24E-03 |
| *MYL4* | myosin light chain 4 | 1.20 | 1.14E-02 |
| *KCNK15* | potassium two pore domain channel subfamily K member 15 | 1.19 | 1.17E-02 |
| *C1QTNF3* | C1q and TNF related 3 | 1.09 | 4.03E-02 |
| *ADGRG1* | adhesion G protein-coupled receptor G1 | 1.08 | 3.84E-02 |
| *GAP43* | growth associated protein 43 | 1.05 | 4.67E-02 |
| *E2F7* | E2F transcription factor 7 | 1.05 | 9.31E-03 |
| *RPLP0P2* | ribosomal protein lateral stalk subunit P0 pseudogene 2 | 1.03 | 0.048507883 |
| *SELPLG* | selectin P ligand | 1.01 | 3.45E-02 |
| *DOK5* | docking protein 5 | 0.98 | 1.31478E-05 |
| *MEST* | mesoderm specific transcript | 0.98 | 2.45E-02 |
| *CIART* | circadian associated repressor of transcription | 0.93 | 5.76E-03 |
| *STARD10* | StAR related lipid transfer domain containing 10 | 0.87 | 1.42E-02 |
| *SSC5D* | scavenger receptor cysteine rich family member with 5 domains | 0.86 | 3.89E-02 |
| *PLAU* | plasminogen activator, urokinase | 0.80 | 5.04E-03 |
| *ZBTB16* | zinc finger and BTB domain containing 16 | -4.46 | 1.60471E-74 |
| *GALNT15* | polypeptide N-acetylgalactosaminyltransferase 15 | -2.85 | 2.52E-26 |
| *FKBP5* | FKBP prolyl isomerase 5 | -2.75 | 1.25837E-38 |
| *HIF3A* | hypoxia inducible factor 3 subunit alpha | -2.57 | 1.20E-13 |
| *GGT5* | gamma-glutamyltransferase 5 | -2.45 | 1.5993E-19 |
| *LEP* | leptin | -2.44 | 2.21481E-18 |
| *PILRA* | paired immunoglobin like type 2 receptor alpha | -2.38 | 1.20227E-15 |
| *PER1* | period circadian regulator 1 | -2.27 | 2.21481E-18 |
| *GRIA1* | glutamate ionotropic receptor AMPA type subunit 1 | -2.26 | 1.63E-17 |

| | | | |
|---|---|---|---|
| *RAPGEF5* | Rap guanine nucleotide exchange factor 5 | -2.17 | 5.04E-09 |
| *ADRA1B* | adrenoceptor alpha 1B | -2.00 | 4.29943E-14 |
| *NEGR1* | neuronal growth regulator 1 | -2.00 | 8.53E-15 |
| *MAOA* | monoamine oxidase A | -1.92 | 1.28694E-09 |
| *CYP4B1* | cytochrome P450 family 4 subfamily B member 1 | -1.89 | 7.91E-07 |
| *CRISPLD2* | cysteine rich secretory protein LCCL domain containing 2 | -1.85 | 6.85088E-09 |
| *LMO3* | LIM domain only 3 | -1.82 | 3.94E-08 |
| *FGD4* | FYVE, RhoGEF and PH domain containing 4 | -1.82 | 5.34309E-09 |
| *GPX3* | glutathione peroxidase 3 | -1.78 | 7.0809E-06 |
| *PLXNA4* | plexin A4 | -1.76 | 5.03299E-08 |
| *INHBB* | inhibin subunit beta B | -1.76 | 2.25E-07 |
| *FMO2* | flavin containing dimethylaniline monoxygenase 2 | -1.69 | 2.02E-05 |
| *RASL11A* | RAS like family 11 member A | -1.69 | 9.70807E-07 |
| *SLC16A12* | solute carrier family 16 member 12 | -1.67 | 4.15847E-05 |
| *DPEP1* | dipeptidase 1 | -1.61 | 0.000103228 |
| *TIMP4* | TIMP metallopeptidase inhibitor 4 | -1.61 | 9.50598E-05 |
| *PRODH* | proline dehydrogenase 1 | -1.61 | 0.000103678 |
| *POM121L9P* | POM121 transmembrane nucleoporin like 9, pseudogene | -1.58 | 9.71E-06 |
| *ADARB1* | adenosine deaminase RNA specific B1 | -1.55 | 5.04E-09 |
| *NA* | NA | -1.52 | 0.000303833 |
| *VIT* | vitrin | -1.52 | 5.03299E-08 |
| *EPHB6* | EPH receptor B6 | -1.50 | 1.04E-04 |
| *ANGPTL1* | angiopoietin like 1 | -1.49 | 1.73E-06 |
| *LRRN3* | leucine rich repeat neuronal 3 | -1.47 | 0.000466681 |
| *MGP* | matrix Gla protein | -1.44 | 0.00060328 |
| *METTL7A* | methyltransferase like 7A | -1.43 | 0.000212293 |
| *ACKR2* | atypical chemokine receptor 2 | -1.42 | 0.000617651 |
| *IGF2* | insulin like growth factor 2 | -1.40 | 0.00111228 |
| *ART4* | ADP-ribosyltransferase 4 (inactive) (Dombrock blood group) | -1.37 | 0.001870234 |
| *HECW1* | HECT, C2 and WW domain containing E3 ubiquitin protein ligase 1 | -1.37 | 1.24E-04 |
| *SCARA5* | scavenger receptor class A member 5 | -1.37 | 0.002350735 |
| *NPR3* | natriuretic peptide receptor 3 | -1.35 | 5.81045E-06 |
| *GLUL* | glutamate-ammonia ligase | -1.32 | 2.94977E-06 |

| | | | |
|---|---|---|---|
| *KIAA0040* | KIAA0040 | -1.31 | 0.003788274 |
| *TSC22D3* | TSC22 domain family member 3 | -1.31 | 9.11628E-09 |
| *APCDD1* | APC down-regulated 1 | -1.30 | 0.002783793 |
| *SLA* | Src like adaptor | -1.30 | 0.004192613 |
| *LRP1B* | LDL receptor related protein 1B | -1.30 | 0.003937068 |
| *CRLF1* | cytokine receptor like factor 1 | -1.27 | 0.000138219 |
| *WASF3* | WASP family member 3 | -1.27 | 3.109E-15 |
| *CYP8B1* | cytochrome P450 family 8 subfamily B member 1 | -1.27 | 0.00566502 |
| *ERRFI1* | ERBB receptor feedback inhibitor 1 | -1.26 | 4.03827E-07 |
| *KLF9* | Kruppel like factor 9 | -1.25 | 6.00664E-11 |
| *NA* | NA | -1.24 | 0.000621609 |
| *GPM6B* | glycoprotein M6B | -1.24 | 0.008587625 |
| *GDF7* | growth differentiation factor 7 | -1.23 | 0.009956326 |
| *TENT5B* | terminal nucleotidyltransferase 5B | -1.23 | 0.000730265 |
| *WNT5A* | Wnt family member 5A | -1.23 | 0.003057797 |
| *STC1* | stanniocalcin 1 | -1.22 | 0.011366869 |
| *SOX13* | SRY-box transcription factor 13 | -1.22 | 4.41E-09 |
| *NGFR* | nerve growth factor receptor | -1.21 | 0.009122867 |
| *GREB1L* | GREB1 like retinoic acid receptor coactivator | -1.21 | 0.000467389 |
| *ADH1A* | alcohol dehydrogenase 1A (class I), alpha polypeptide | -1.21 | 0.011366869 |
| *RGCC* | regulator of cell cycle | -1.21 | 1.14E-02 |
| *CD34* | CD34 molecule | -1.20 | 0.014474888 |
| *ALOX5AP* | arachidonate 5-lipoxygenase activating protein | -1.20 | 1.14E-02 |
| *NEXN* | nexilin F-actin binding protein | -1.19 | 2.01059E-05 |
| *GUCY1A1* | guanylate cyclase 1 soluble subunit alpha 1 | -1.19 | 0.001682657 |
| *TACC1* | transforming acidic coiled-coil containing protein 1 | -1.18 | 2.21085E-07 |
| *APOB* | apolipoprotein B | -1.17 | 0.020311162 |
| *CPM* | carboxypeptidase M | -1.16 | 0.019372744 |
| *SMCO3* | single-pass membrane protein with coiled-coil domains 3 | -1.16 | 0.017528199 |
| *NEBL* | nebulette | -1.15 | 0.020962114 |
| *NID1* | nidogen 1 | -1.15 | 1.8104E-08 |
| *OMD* | osteomodulin | -1.14 | 0.023125643 |
| *MAOB* | monoamine oxidase B | -1.14 | 1.29E-02 |
| *NA* | NA | -1.12 | 0.006808232 |
| *CFD* | complement factor D | -1.12 | 0.001282946 |

| | | | |
|---|---|---|---|
| *C6* | complement C6 | -1.12 | 0.017236773 |
| *DSEL* | dermatan sulfate epimerase like | -1.12 | 1.76E-03 |
| *RGMA* | repulsive guidance molecule BMP co-receptor a | -1.11 | 0.000206401 |
| *LINC01088* | long intergenic non-protein coding RNA 1088 | -1.11 | 0.034162437 |
| *GUCY1B1* | guanylate cyclase 1 soluble subunit beta 1 | -1.10 | 0.015107554 |
| *TRNP1* | TMF1 regulated nuclear protein 1 | -1.10 | 0.00120906 |
| *TCEAL4* | transcription elongation factor A like 4 | -1.10 | 2.78E-06 |
| *TMC2* | transmembrane channel like 2 | -1.09 | 4.25E-02 |
| *DPT* | dermatopontin | -1.09 | 0.042483615 |
| *HLX* | H2.0 like homeobox | -1.08 | 2.23E-06 |
| *CDKN1C* | cyclin dependent kinase inhibitor 1C | -1.08 | 0.024503934 |
| *OLAH* | oleoyl-ACP hydrolase | -1.08 | 0.040638212 |
| *PIK3R1* | phosphoinositide-3-kinase regulatory subunit 1 | -1.08 | 0.040327671 |
| *NRCAM* | neuronal cell adhesion molecule | -1.08 | 0.04190168 |
| *IRS2* | insulin receptor substrate 2 | -1.07 | 0.000329481 |
| *RSPO1* | R-spondin 1 | -1.07 | 0.045696184 |
| *RNF144B* | ring finger protein 144B | -1.06 | 0.037658048 |
| *DNAJC6* | DnaJ heat shock protein family (Hsp40) member C6 | -1.05 | 0.000467389 |
| *ITPR1* | inositol 1,4,5-trisphosphate receptor type 1 | -1.05 | 0.00930819 |
| *DHRS3* | dehydrogenase/reductase 3 | -1.05 | 0.008570022 |
| *CST3* | cystatin C | -1.05 | 2.4235E-06 |
| *FOXO1* | forkhead box O1 | -1.04 | 0.00043938 |
| *RAB40A* | RAB40A, member RAS oncogene family | -1.04 | 0.042358847 |
| *TMEM119* | transmembrane protein 119 | -1.03 | 0.008587625 |
| *STK17B* | serine/threonine kinase 17b | -1.03 | 0.040327671 |
| *NAV2* | neuron navigator 2 | -1.02 | 0.003057797 |
| *TLR4* | toll like receptor 4 | -1.02 | 0.004507392 |
| *PLCL1* | phospholipase C like 1 (inactive) | -1.00 | 0.009287195 |
| *PRELP* | proline and arginine rich end leucine rich repeat protein | -1.00 | 0.014096681 |
| *COBLL1* | cordon-bleu WH2 repeat protein like 1 | -0.99 | 0.001219936 |
| *NFASC* | neurofascin | -0.99 | 2.21085E-07 |
| *TGFBR2* | transforming growth factor beta receptor 2 | -0.99 | 9.94915E-16 |
| *ANGPT1* | angiopoietin 1 | -0.98 | 0.017236773 |

| | | | |
|---|---|---|---|
| *SSH2* | slingshot protein phosphatase 2 | -0.97 | 0.000502181 |
| *ZNF395* | zinc finger protein 395 | -0.97 | 0.00159517 |
| *MINDY2* | MINDY lysine 48 deubiquitinase 2 | -0.96 | 0.000108694 |
| *TNS2* | tensin 2 | -0.96 | 0.000161324 |
| *PDE3A* | phosphodiesterase 3A | -0.96 | 0.011413361 |
| *SH3D19* | SH3 domain containing 19 | -0.95 | 0.013368087 |
| *CALCOCO2* | calcium binding and coiled-coil domain 2 | -0.95 | 0.000566645 |
| *ARMC2* | armadillo repeat containing 2 | -0.93 | 0.00159517 |
| *RASSF4* | Ras association domain family member 4 | -0.93 | 0.006062503 |
| *BMPR1B* | bone morphogenetic protein receptor type 1B | -0.93 | 0.022685107 |
| *CAB39L* | calcium binding protein 39 like | -0.92 | 0.00519468 |
| *SETBP1* | SET binding protein 1 | -0.91 | 0.000206401 |
| *ZHX3* | zinc fingers and homeoboxes 3 | -0.91 | 0.003813157 |
| *PARD3B* | par-3 family cell polarity regulator beta | -0.91 | 0.000354793 |
| *MT1X* | metallothionein 1X | -0.91 | 0.009984076 |
| *IRS1* | insulin receptor substrate 1 | -0.91 | 0.000114999 |
| *LAMA2* | laminin subunit alpha 2 | -0.90 | 0.038677568 |
| *COL5A3* | collagen type V alpha 3 chain | -0.90 | 7.90824E-05 |
| *JAK2* | Janus kinase 2 | -0.89 | 0.012378855 |
| *FOXO3* | forkhead box O3 | -0.89 | 2.4235E-06 |
| *ITGA1* | integrin subunit alpha 1 | -0.88 | 0.007903682 |
| *HPS5* | HPS5 biogenesis of lysosomal organelles complex 2 subunit 2 | -0.88 | 0.047592551 |
| *SPART* | spartin | -0.88 | 0.00281968 |
| *SERPING1* | serpin family G member 1 | -0.87 | 0.022949188 |
| *FAT4* | FAT atypical cadherin 4 | -0.86 | 0.038604148 |
| *EPN2* | epsin 2 | -0.86 | 0.009815046 |
| *ANG* | angiogenin | -0.86 | 0.000138725 |
| *PDGFRA* | platelet derived growth factor receptor alpha | -0.85 | 0.028791744 |
| *SMARCD2* | SWI/SNF related, matrix associated, actin dependent regulator of chromatin, subfamily d, member 2 | -0.84 | 1.81995E-05 |
| *FBLN2* | fibulin 2 | -0.83 | 0.028745433 |
| *NA* | NA | -0.83 | 0.005039655 |
| *RNASE4* | ribonuclease A family member 4 | -0.82 | 0.003862282 |
| *FOXN3* | forkhead box N3 | -0.80 | 5.5559E-06 |
| *ARRDC2* | arrestin domain containing 2 | -0.80 | 0.018317278 |

| | | | |
|---|---|---|---|
| *PRRG1* | proline rich and Gla domain 1 | -0.79 | 0.042711244 |
| *TP53I11* | tumor protein p53 inducible protein 11 | -0.78 | 0.012793999 |
| *FAM13A* | family with sequence similarity 13 member A | -0.77 | 3.14E-02 |
| *CUTC* | cutC copper transporter | -0.77 | 0.000125811 |
| *PTK2B* | protein tyrosine kinase 2 beta | -0.77 | 0.000103678 |
| *SIDT2* | SID1 transmembrane family member 2 | -0.77 | 9.76834E-05 |
| *LRP4* | LDL receptor related protein 4 | -0.76 | 0.011613502 |
| *PLEKHA2* | pleckstrin homology domain containing A2 | -0.75 | 0.029976761 |
| *CASTOR3* | CASTOR family member 3 | -0.74 | 1.13E-03 |
| *RELL1* | RELT like 1 | -0.73 | 0.000206401 |
| *FEZ2* | fasciculation and elongation protein zeta 2 | -0.73 | 0.001474034 |
| *AFF1* | AF4/FMR2 family member 1 | -0.73 | 0.040703416 |
| *CXCL12* | C-X-C motif chemokine ligand 12 | -0.71 | 0.009984076 |
| *PCYOX1* | prenylcysteine oxidase 1 | -0.71 | 0.002231091 |
